# Independent evolution of flavour and parthenocarpy in fig, one of the earliest domesticated fruit trees

**DOI:** 10.64898/2026.08.09.743143

**Authors:** Hidetoshi Ikegami, Takeshi Hayashi, Shiori Yabe, Kenta Shirasawa, Masaru Sato, Hideyuki Suzuki, Kosuke Tashiro, Kazuki Mori, Hiroshi Yakushiji, Ikue Yoshikawa, Misaki Ishibashi, Katsuhiro Shiratake, Chiharu Hirata, Hitoshi Nogata

## Abstract

Domestication often modifies multiple traits in concert, but whether fruit quality and reproductive mode share a genetic basis in perennial crops has not been tested at multi-omics resolution. In fig (*Ficus carica*), one of the earliest domesticated fruit trees and a crop with archaeobotanical evidence of cultivation over 11,000 years ago, domestication yielded both distinctive flavour and parthenocarpy, fruit development without pollination. We integrated whole-genome resequencing, transcriptomics, volatile and metabolite profiling, and taste sensor data across a diverse accession panel, using Bayesian GWAS/TWAS and multi-omics factor analysis. Flavour emerges as an integrated sensory system led by aroma, which modulates taste through cross-modal interactions, with sugar–acid balance as a secondary axis; these quality traits map to numerous independent loci. Parthenocarpy instead traces to a single perfectly concordant chromosome 04 haplotype (*Eden*) centred on *FcMYB101-like*, carrying a selective-sweep signature and a derived allele fixed in cultivated figs but rare across the genus. Flavour and reproductive mode thus evolved along separate genomic trajectories, allowing each to be improved independently.

## Introduction

Figs (*Ficus carica* L.) are among the earliest domesticated fruit trees, with archaeobotanical evidence from the Jordan Valley indicating intentional cultivation as early as 11,400 years ago^1^—predating cereal domestication and positioning fig as a pivotal model for the origins of agriculture and the domestication of perennial crops. Cultivated Common-type figs are characterised by parthenocarpy—fruit development without pollination—a trait that enabled stable production independent of the obligate fig wasp mutualism and facilitated the crop’s spread across broad agricultural environments^2–5^. The genetic basis of this reproductive innovation remains only partly resolved: classical segregation studies inferred a single dominant locus^2,6^, whereas a recent comparative genomic analysis reported contraction of the s-domain receptor-like kinase family, which mediates pollen recognition, in *F. carica* relative to wild *Ficus* species^7^—a species-level feature that cannot by itself account for the segregation of pollination dependence among cultivars. Possibly building on this reproductive foundation, human selection has generated extensive phenotypic diversity in fruit quality, encompassing sugars, organic acids, pigments, aroma compounds, and taste attributes^3,8^, yet the integrated architecture of this sensory diversity remains uncharacterised. Because domestication often modifies multiple traits in concert, through pleiotropy or through clustering of selected loci^9^, whether fruit quality and reproductive mode in fig share a genetic basis or evolved through independent selective processes remains untested.

Flavour itself is increasingly understood not as a set of independent chemical traits but as a single sensory system in which aroma, taste, and pigmentation co-vary^10^. Aroma compounds perceived retronasally modulate taste intensity and quality, and sugar–acid balance shapes the hedonic response to volatile compounds^10–12^. In annual crops such as tomato, integrative approaches have begun to reveal how these dimensions interact at the metabolic and genetic level^12,13^. In perennial fruit crops, however, systematic multi-omics characterisation of flavour as an integrated phenotype has not been achieved, in part because the large genome sizes, long generation times, and high heterozygosity of tree fruits present substantial methodological challenges^14^. Studies of fig fruit quality have accordingly addressed individual metabolite classes—sugars, organic acids, or volatiles—in isolation^15,16^, leaving the cross-modal interactions and shared genetic architecture that define flavour as a whole unresolved.

Here we address this gap by applying an integrated multi-omics strategy to a diverse panel of *Ficus carica* accessions. We combine whole-genome resequencing (n = 482), RNA-seq transcriptomics, GC–MS volatile profiling, GC–MS quantification of primary metabolites, probe electrospray ionization tandem mass spectrometry (PESI-MS/MS) quantification of anthocyanins, and taste sensor measurements with Bayesian GWAS, transcriptome-wide association study (TWAS), and multi-omics factor analysis (MOFA+)^17^. This design allows us to characterise fruit flavour at the sensory, biochemical, and genomic levels simultaneously, resolving both the latent structure linking omics layers and the loci underlying individual quality traits. In parallel, we apply the same genomic framework to parthenocarpy, identifying a perfectly concordant haplotype on chromosome 04 (locus symbol *Eden*) centred on the candidate gene *FcMYB101-like*. This haplotype carries a selective-sweep signature and, based on genotypes recovered from wild *Ficus* outgroup species, a derived allele that reaches fixation in cultivated *F. carica* but is otherwise rare across the genus, consistent with positive selection for parthenocarpy during domestication. Together, our results reveal that fig fruit flavour is organised as an integrated sensory system in which aroma is the dominant axis among sensory omics layers, and that fruit quality and parthenocarpy represent evolutionarily independent axes of fig domestication governed by distinct genomic layers.

## Results

### Genetic and phenotypic diversity of the fig accession panel

To capture the breadth of genetic diversity in cultivated fig, we performed whole-genome resequencing of 482 accessions spanning the major horticultural classes of *F. carica* (Common-, San Pedro-, and Smyrna-type, and Caprifig), F1 progeny from crosses within *F. carica*, *F. carica* × *F. erecta* interspecific hybrid populations, and wild and closely related *Ficus* species, including 27 previously published genomes^18,19^; the full panel composition is given in Supplementary Data 1 and Methods. After quality filtering, 3,275,911 high-confidence SNPs were identified (Ts/Tv = 1.66), distributed predominantly in intergenic (61.2%) and intronic (22.9%) regions, with 15.1% exonic; SnpEff classified 0.3% (n = 10,031) of sites as high-effect variants (Supplementary Fig. 1). This dataset provides the genomic foundation for the association analyses below.

### Population structure and inter-accession relationships

To examine relationships among accessions, we performed principal component analysis (PCA) and neighbour-joining tree analysis on the full quality-filtered SNP dataset (3,275,911 SNPs), using a pairwise 1−IBS genetic distance matrix computed in PLINK^20^ and the nj() function in the R package ape^21^. The first two principal components explained 23.59% and 19.6% of total genetic variance, respectively (Figure 1A).

**Figure 1.**
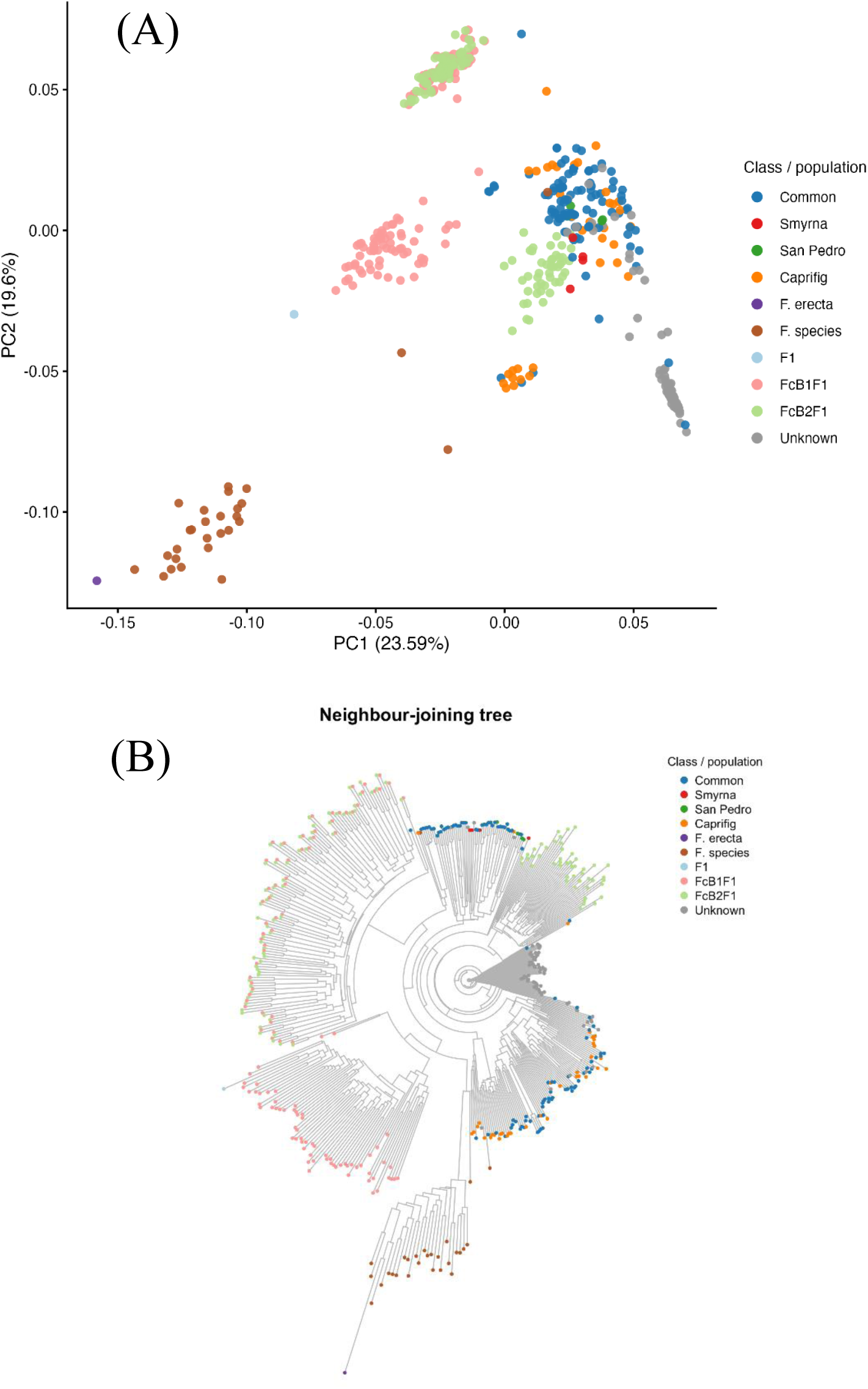
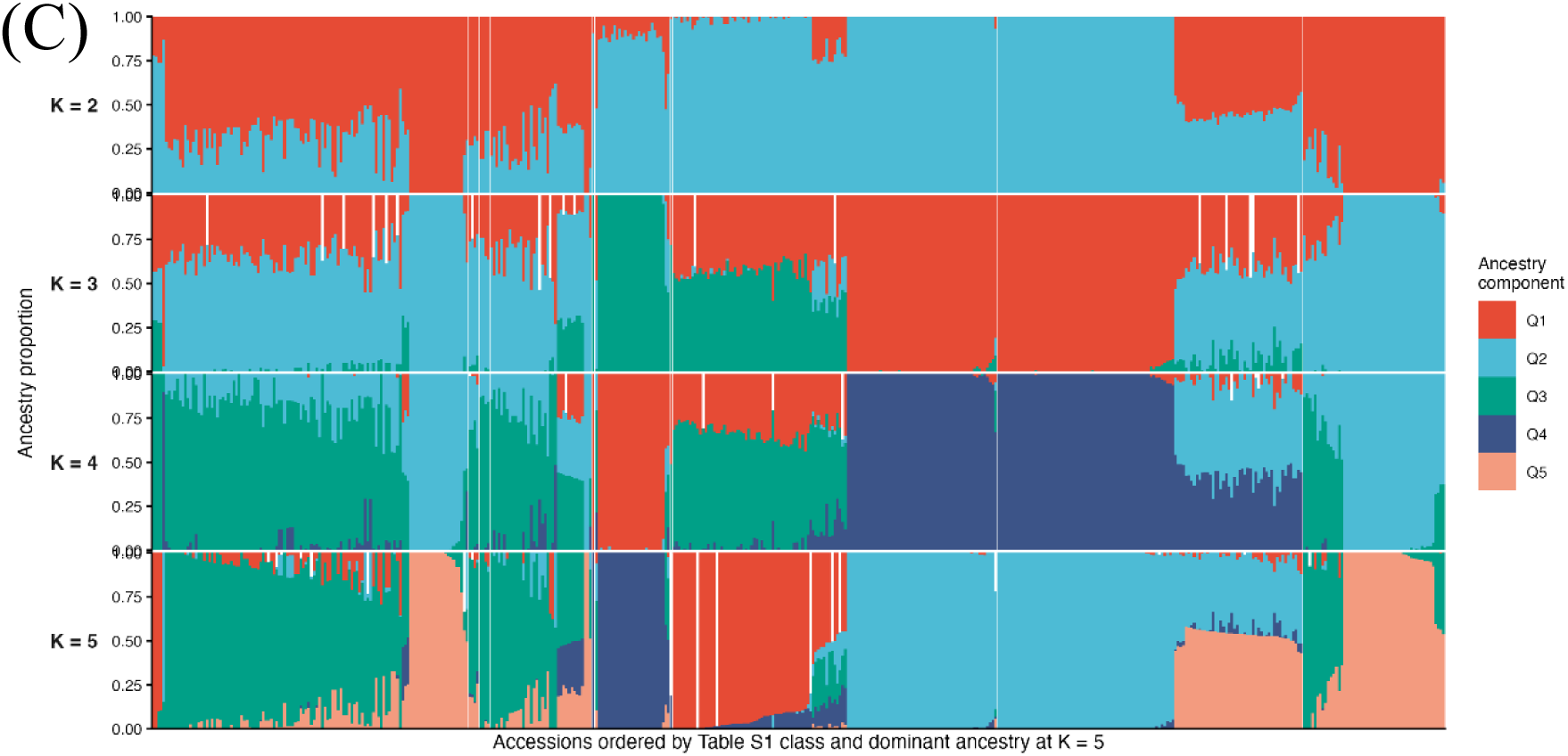
Genetic diversity and population structure of the *Ficus* accession panel. (A) Principal component analysis (PCA) of 482 accessions based on the LD-pruned SNP dataset (164,933 SNPs). Each point is one accession, coloured by horticultural class (Common, Smyrna, San Pedro, Caprifig, *F. erecta*, Hybrid breeding line), progeny population (F1, FcB1F1, FcB2F1), or designation (Parent, Published_Ficus). PC1 and PC2 explained 23.59% and 19.6% of total variance. The Published_Ficus category comprises 27 previously published *Ficus* genomes^18,19^: one *F. carica* accession (FcarWG116), which clusters within the *F. carica* clade, and 26 accessions of 13 wild *Ficus* species forming a distinct cluster along PC1. The *F. palmata* accession (Fc_31) clusters near Caprifig accessions. Horticultural classes of *F. carica* were broadly co-distributed without discrete separation. (B) Neighbour-joining tree constructed from a pairwise 1−IBS genetic distance matrix computed from the same LD-pruned dataset. Branch tips are coloured as in (A). FcB1F1 and FcB2F1 accessions occupy positions intermediate between *F. carica* and *F. erecta*; the published *Ficus* accessions form a distinct clade with the exception of FcarWG116, and *F. palmata* (Fc_31) clusters near Caprifig accessions. (C) ADMIXTURE ancestry proportions for K = 2–5 based on the same LD-pruned dataset. Each vertical bar is one accession, coloured by inferred ancestry component (Q1–Q5); accessions are ordered by horticultural class (Supplementary Data 1) and by dominant component at K = 5. K = 2 separates the interspecific hybrid populations from *F. carica*; K = 3–4 progressively resolve substructure within the cultivated gene pool; K = 5 reveals additional fine-scale differentiation. Full results for K = 2–10 are shown in Supplementary Fig. 2A.

Common-type, Smyrna-type, San Pedro-type, and Caprifig accessions were broadly co-distributed in PCA space without clear separation, indicating that genetic divergence among horticultural classes within *F. carica* is quantitative rather than discrete (Figure 1A). Caprifig accessions—maintained as pollen donors in close association with cultivated lines—shared a largely overlapping genetic background with Common- and Smyrna-type accessions. Published *Ficus* genomes formed a clearly separated cluster along PC1, whereas the F1 hybrid populations (FcB1F1 and FcB2F1) occupied intermediate positions between the parental *F. carica* and *F. erecta* accessions in both PCA space and the neighbour-joining tree, confirming the expected admixture of parental haplotypes (Figure 1A, 1B).

ADMIXTURE analysis of a linkage-disequilibrium-pruned SNP set (164,933 SNPs retained after pruning with PLINK --indep-pairwise 50 10 0.2) resolved a hierarchical ancestry structure across the panel (Figure 1C; Supplementary Fig. 2). At K = 2, the FcB1F1 and FcB2F1 hybrid populations separated from *F. carica* as a discrete ancestry component, reflecting their interspecific origin; K = 3–4 progressively resolved substructure within the cultivated *F. carica* gene pool, and K = 5 revealed additional fine-scale differentiation. Cross-validation error decreased monotonically across K = 2–10 without identifying an optimal value (Supplementary Fig. 2B), indicating fine-scale substructure beyond the tested range and consistent with the continuous, admixed nature of variation within cultivated *F. carica* documented by PCA and the neighbour-joining tree. The full range of K = 2–10 is provided in Supplementary Fig. 2A.

These patterns indicate that phenotypic diversification in fruit quality has proceeded against a background of continuous, multi-layered genetic variation, with no discrete genomic boundary separating horticultural classes—motivating the quantitative association framework applied below.

### Phenotypic diversity across fruit quality layers

To characterise the phenotypic landscape of fruit quality, we profiled four complementary omics layers—primary metabolites (sugars, organic acids, and amino acids), volatile aroma compounds, peel colorimetric parameters and anthocyanins, and taste sensor attributes—and applied hierarchical clustering and PCA to each dataset independently. Across all four layers, phenotypic variation among accessions was quantitative and continuous, with genotypes distributed along smooth gradients in PCA space rather than forming discrete clusters (Figure 2; Supplementary Fig. 3).

**Figure 2.**
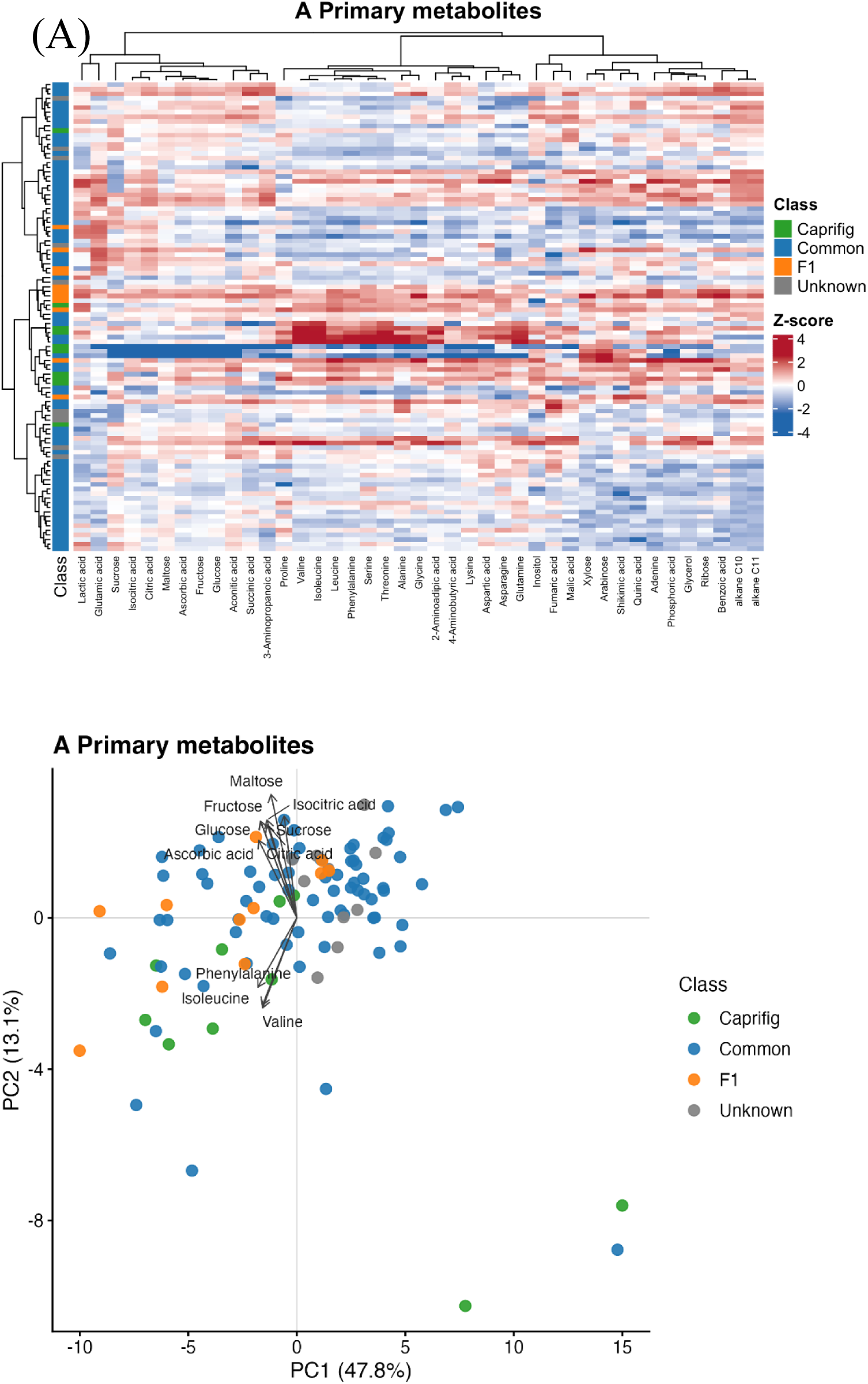
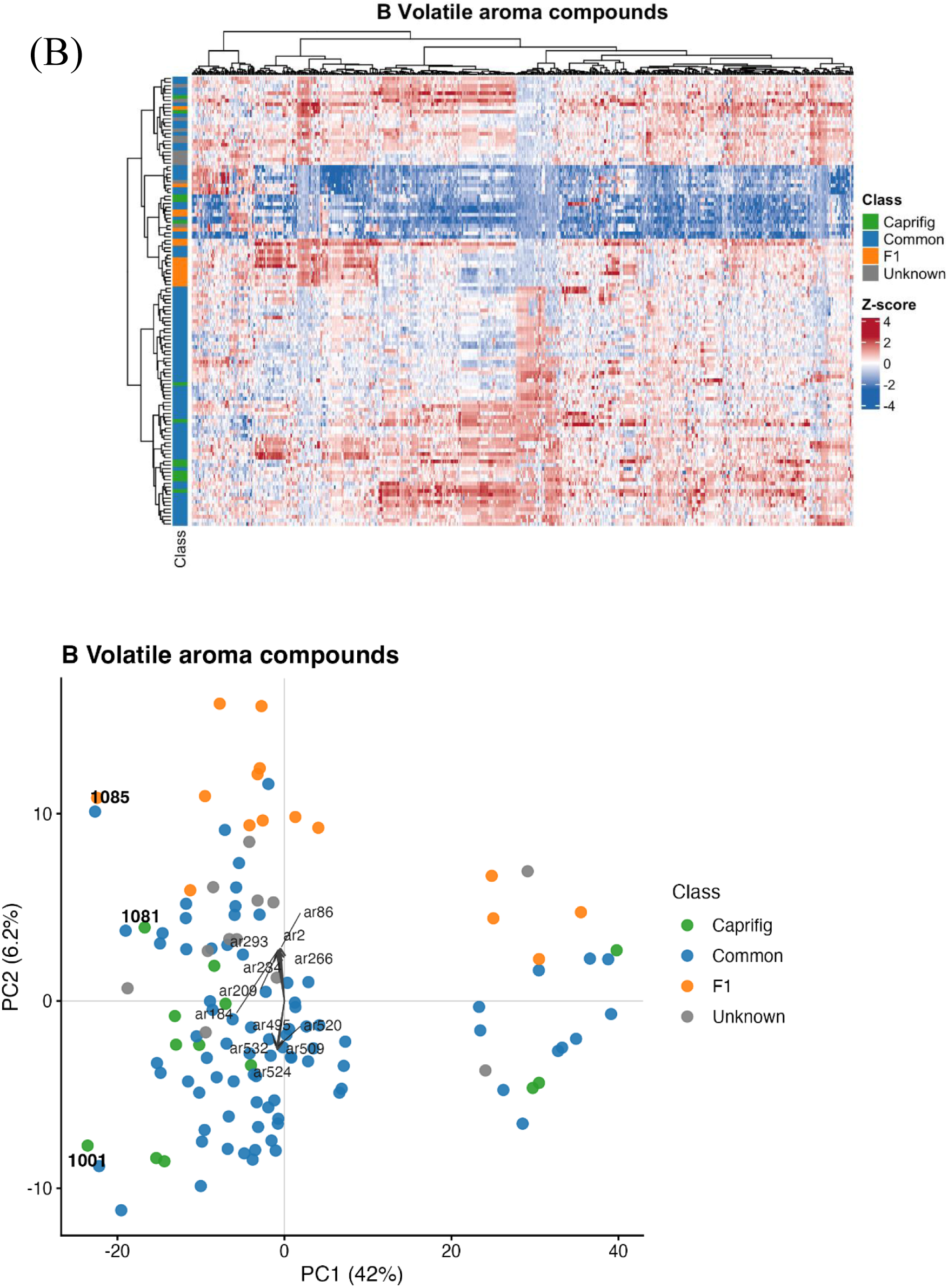
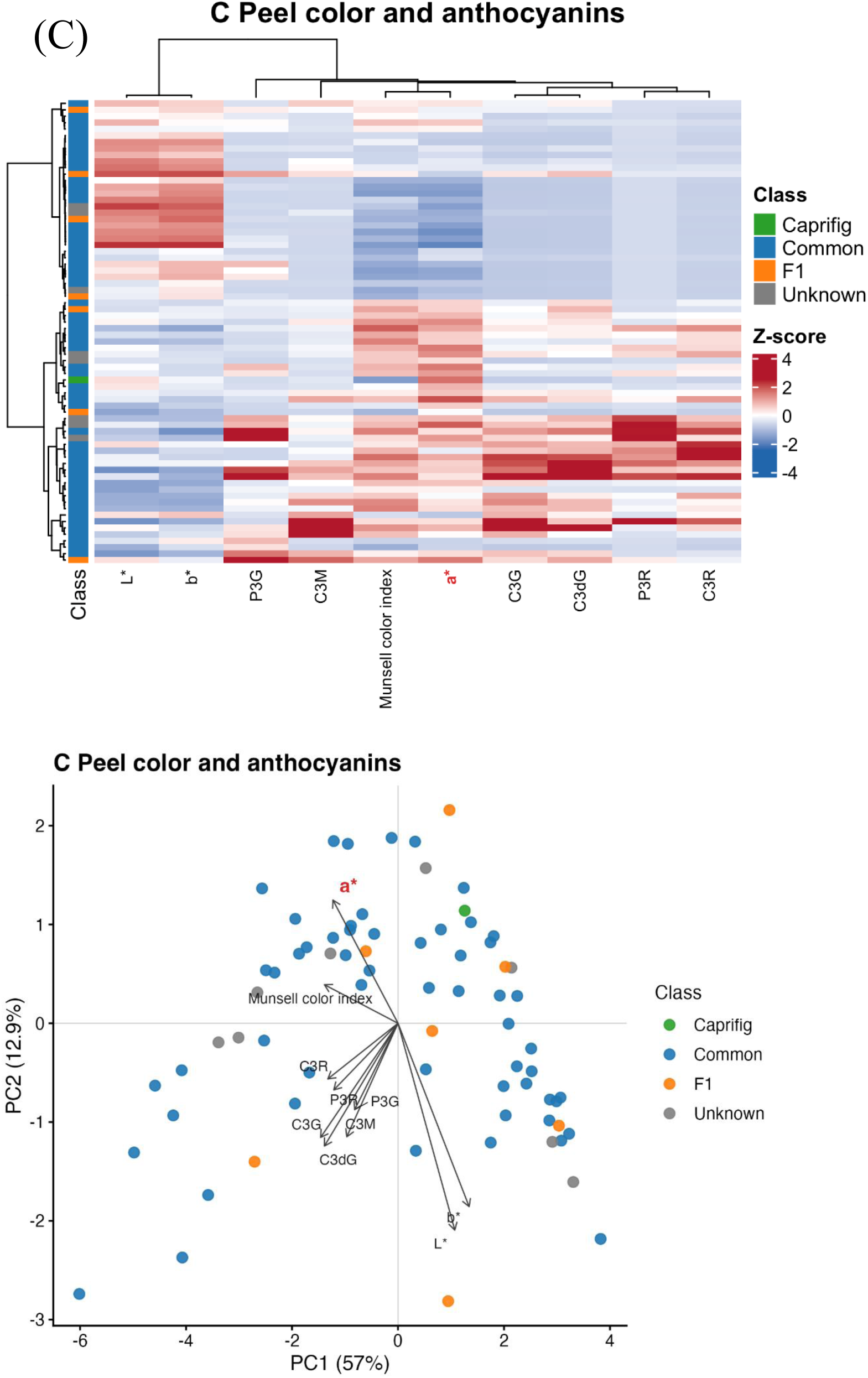
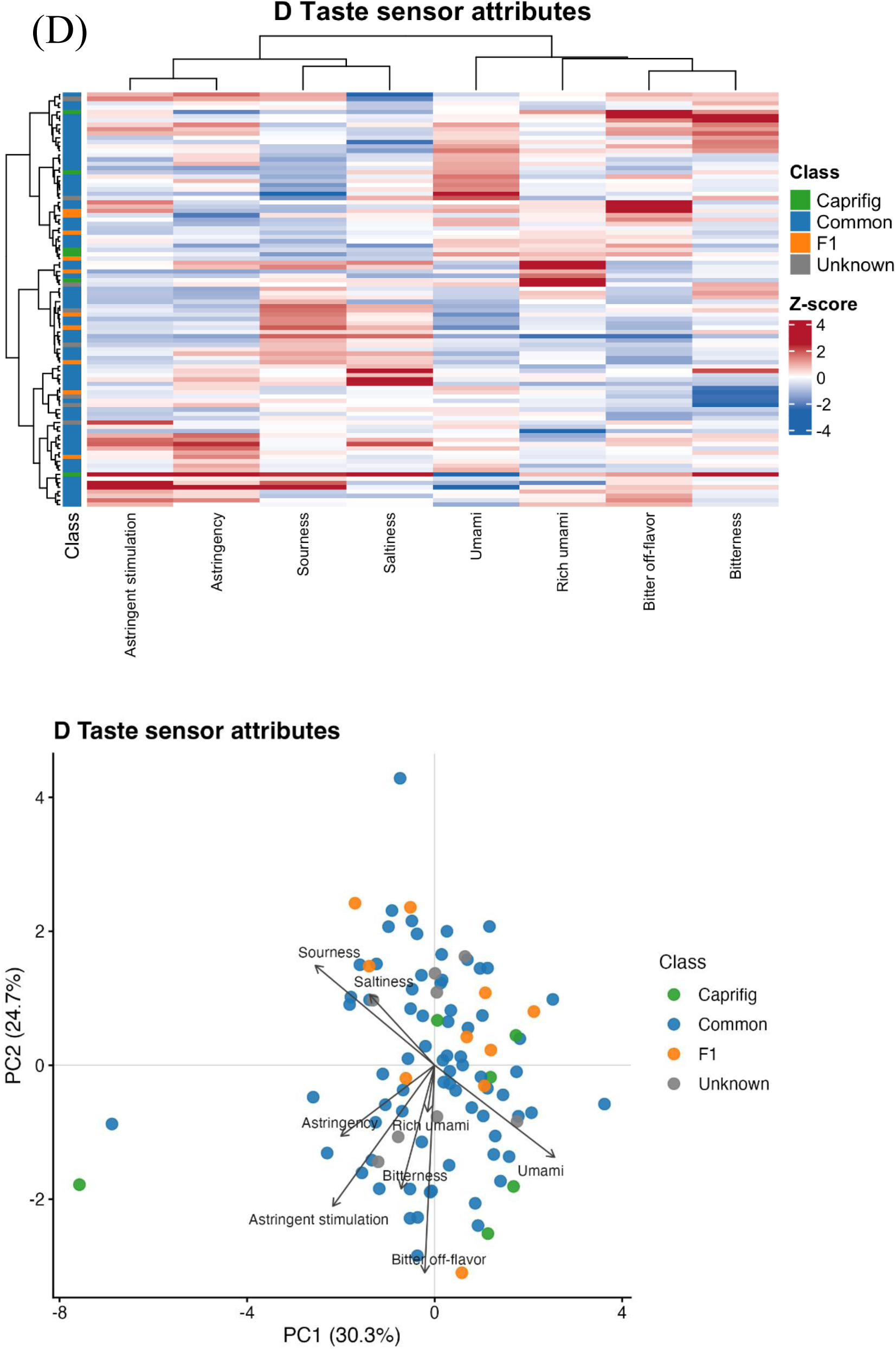
Phenotypic diversity across four fruit-quality omics layers in the *Ficus carica* accession panel. PCA biplots and hierarchical clustering heatmaps for four fruit-quality omics layers. In all PCA plots each point is one accession, coloured by horticultural class (Common-type, Caprifig, F1, or Unknown); loading vectors indicate the direction and relative magnitude of trait contributions to each principal component. Heatmaps show Z-score-scaled trait values, clustered by Ward.D2 linkage. (A) Primary metabolites (n = 102 accessions; 41 parameters). PC1 (47.8%) was dominated by sugars (fructose, glucose, maltose, sucrose) and organic acids (citric, isocitric, ascorbic acid); PC2 (13.1%) by branched-chain and aromatic amino acids (phenylalanine, isoleucine, valine), an independent axis with lower inter-accession variance. Caprifig accessions separated from Common-type accessions along PC1. (B) Volatile aroma (n = 121 accessions; 549 features). PC1 (42.0%) captured the primary axis of volatile diversity. Most accessions occupied a central cluster, whereas accessions 1081, 1001, and 1085 segregated as high-impact aroma variants with disproportionate accumulation of ester-class volatiles (red in the violin plot; Supplementary Fig. 3B) and marked displacement along PC1 and PC2. (C) Peel colorimetry and anthocyanins (n = 82 accessions; 10 parameters). PC1 (57.0%) contrasted anthocyanin species (P3G, P3R, C3G, C3R, C3dG, C3M; negative loadings) with lightness and yellowness (L*, b*; positive loadings), reflecting the red-to-green pigmentation gradient. PC2 (12.9%) was dominated by a* (the red–green colour axis; highlighted in red). (D) Taste sensor profiles (n = 96 accessions; 8 dimensions). PC1 (30.3%) contrasted umami and rich umami (positive loadings) against astringency, bitterness, astringent stimulation, and bitter off-flavor (negative loadings), reflecting a palatability gradient. PC2 (24.7%) contrasted sourness and saltiness against bitter off-flavor and astringent stimulation. PC1 and PC2 together explained 55.0% of total taste variance.

Within this shared pattern, each layer displayed characteristic axes of variation. For primary metabolites, PC1 (47.8%) was dominated by sugars and organic acids (fructose, glucose, maltose, sucrose, citric and isocitric acid), whereas amino acids (phenylalanine, isoleucine, valine) loaded on PC2 (13.1%), an independent axis with markedly lower inter-accession variance; Caprifig accessions separated distinctly from Common-type accessions along PC1 (Figure 2A; Supplementary Fig. 3A). For aroma, PC1 (42%) captured the primary axis of volatile diversity, with most accessions occupying a central cluster and three accessions (1081, 1001, 1085) segregating as high-impact variants characterised by disproportionate accumulation of ester-class volatiles (Figure 2B; Supplementary Fig. 3B). For colour, PC1 (57%) contrasted the six anthocyanin species with lightness and yellowness (L*, b*), reflecting the red-to-green pigmentation gradient, while PC2 (12.9%) was dominated by a*; hierarchical clustering placed a*, Munsell colour index, and all six anthocyanins in a single trait cluster distinct from L* and b* (Figure 2C; Supplementary Fig. 3C). For taste, PC1 (30.3%) contrasted umami and rich umami against astringency, bitterness, astringent stimulation, and bitter off-flavor, defining a palatability gradient, and PC2 (24.7%) contrasted sourness and saltiness against aversive aftertaste; clustering resolved a corresponding astringency–sourness–saltiness group and an umami–bitterness group (Figure 2D; Supplementary Fig. 3D).

No single layer captured the full extent of flavour diversity, and the relationships among layers—how aroma variation co-varies with taste attributes, and how primary metabolite composition interacts with volatile profiles—could not be resolved by single-layer analyses, motivating the integrated multi-omics approach applied below.

### A perfectly concordant haplotype centred on MYB101-like on chromosome 04 marks parthenocarpy in Ficus carica

Parthenocarpy—fruit development without pollination—is the defining reproductive innovation of cultivated Common-type figs and a prerequisite for agricultural production independent of the fig wasp mutualism. To identify its genomic basis, we performed a genotype–phenotype concordance scan comparing 118 non-Smyrna parthenocarpic accessions (114 Common-type, 4 San Pedro-type) against 4 Smyrna-type accessions, using SNPs mapped independently to both the female-derived (FCAr2.3) and male-derived (FCA6085_r1.1) assemblies; the small Smyrna-type sample precluded quantitative association analysis, motivating this concordance-based approach.

The scan identified 10 perfectly concordant SNPs genome-wide: eight clustered within a single chromosome 04 interval (Figure 3A–C), and two were isolated singletons on chromosomes 01a and 05 without regional clustering or functional support (Supplementary Fig. 4A, B). All 118 non-Smyrna cases carried dosage 0 at each chr04 SNP, while all 4 Smyrna-type controls were heterozygous at each position—perfect genotype–phenotype separation across the panel (Figure 3C; Supplementary Fig. 4E), implying that parthenocarpic fruit set requires d-type homozygosity (see Discussion). The 4 San Pedro-type accessions, whose breba crop sets parthenocarpically while the main crop requires caprification, carried the non-Smyrna haplotype at all 8 SNPs, indicating that the haplotype tracks the capacity for parthenocarpic fruit set rather than its crop-specific expression. The concordant SNPs were confirmed in the non-imputed dataset (Supplementary Fig. 4D), reproduced identically using the FCA6085_r1.1 assembly (Figure 3A), and supported by permutation testing (10,000 replicates; empirical P = 0.0001 for chromosome 04 and P = 0.0003 genome-wide; Figure 4A).

**Figure 3.**
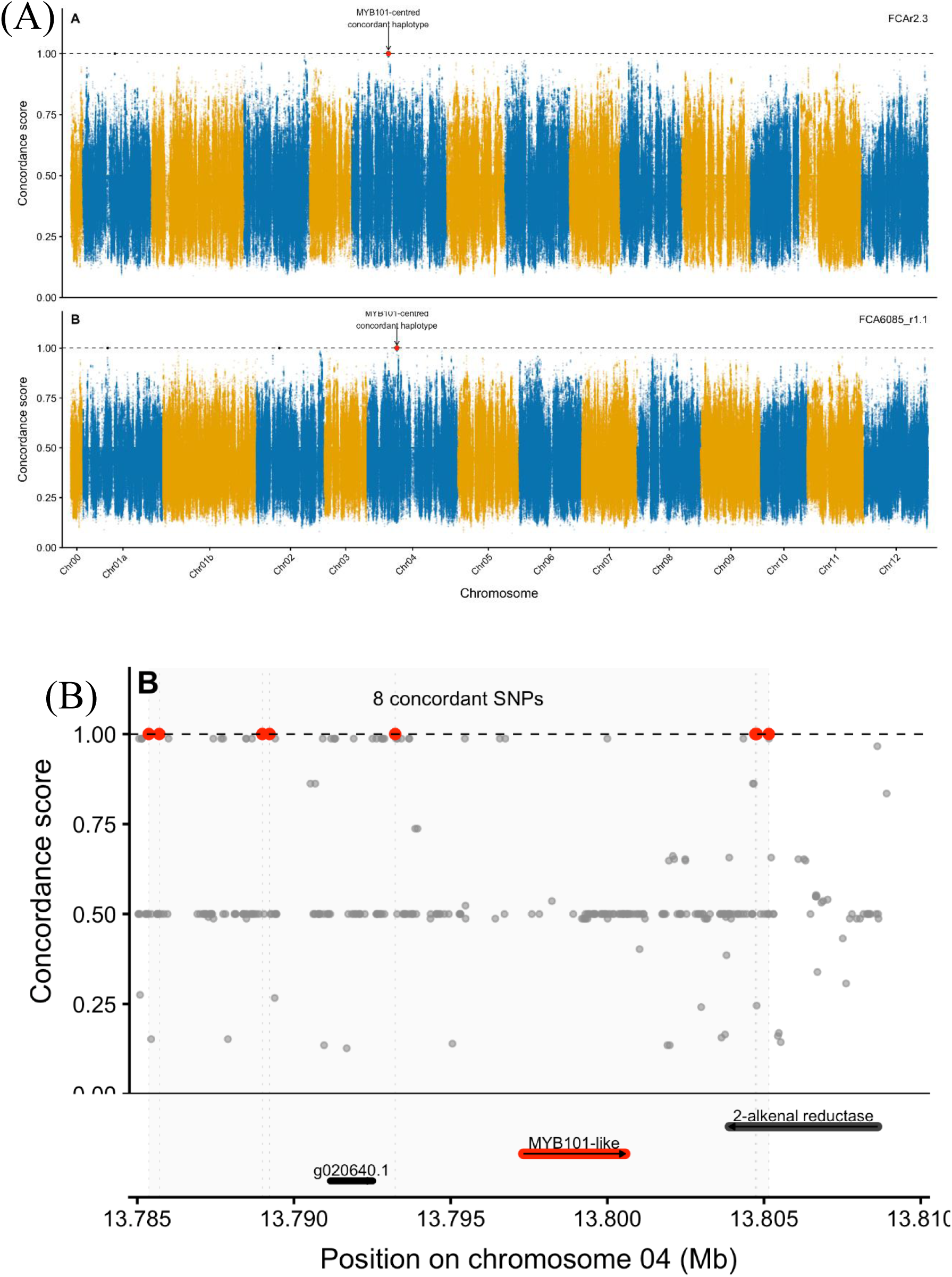
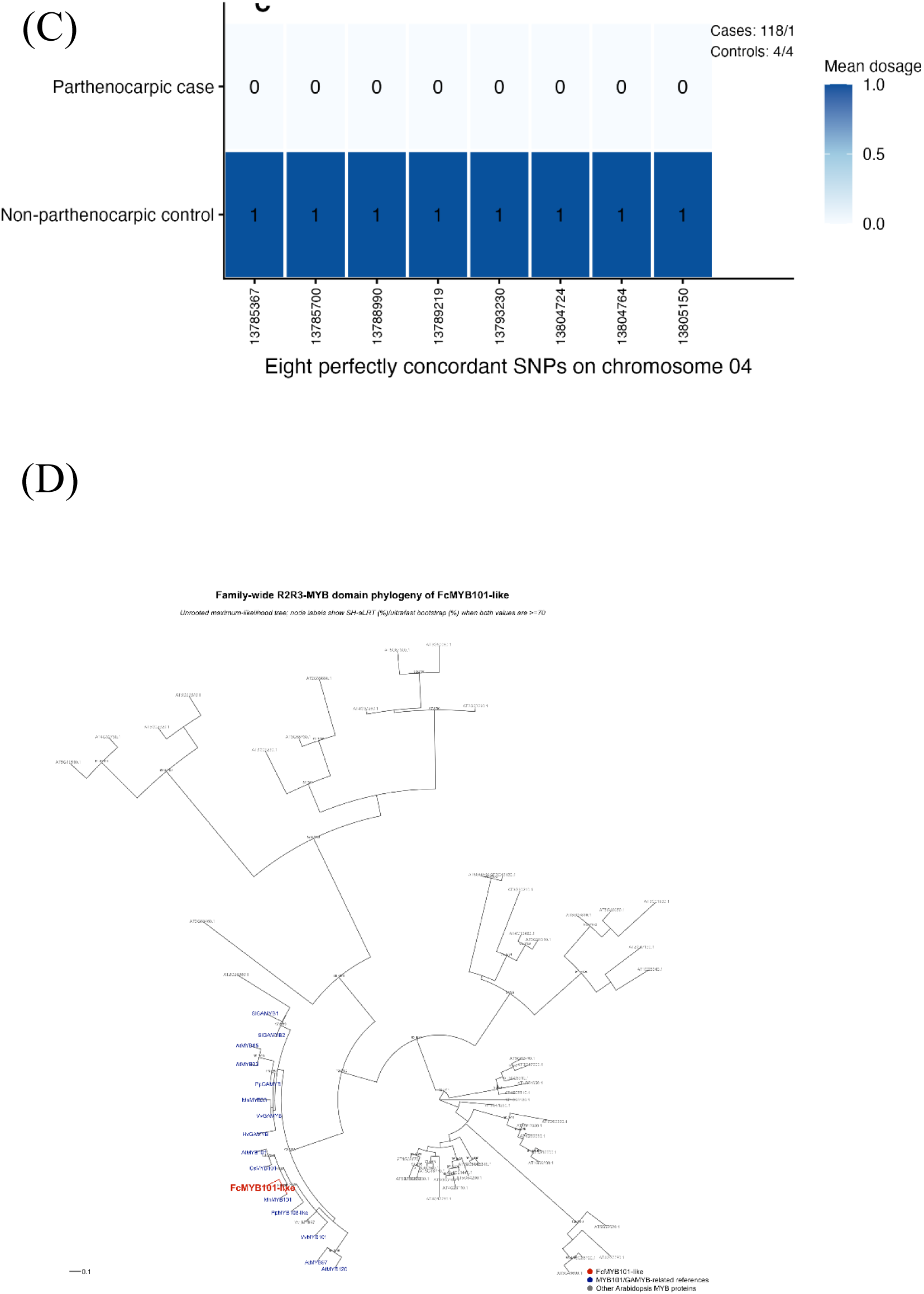
A perfectly concordant haplotype centred on *MYB101-like* on chromosome 04 marks parthenocarpy in *Ficus carica*. (A) Genome-wide genotype–phenotype concordance scans for parthenocarpy, using SNP datasets mapped independently to the FCAr2.3 (upper) and FCA6085_r1.1 (lower) assemblies. x-axis, chromosomal position; y-axis, concordance score between SNP dosage and a binary parthenocarpy model. Perfectly concordant SNPs (score = 1.0) are red. A cluster of 8 perfectly concordant SNPs was detected on chromosome 04 in both assemblies (permutation test, Figure 4A); two isolated concordant SNPs on chromosomes 01a and 05 lack regional clustering or annotation support. (B) Zoomed view of the chromosome 04 candidate interval (13.785–13.810 Mb in FCAr2.3). Upper, SNP-level concordance scores across the region, with the 8 perfectly concordant SNPs in red; shading marks the candidate interval. Lower, annotated gene models in the interval: g020640.1, *MYB101-like* (g020650.1), and 2-alkenal reductase (g020660.1). One concordant SNP lies approximately 4.1 kb upstream of *MYB101-like*; no concordant SNP was detected within the *MYB101-like* coding sequence in the filtered dataset (see Supplementary Fig. 4F). (C) Mean dosage of the counted allele at the eight concordant chromosome 04 SNPs across phenotyped accessions, grouped by reproductive phenotype. All 118 parthenocarpic cases carried dosage 0 at each SNP, whereas all 4 Smyrna-type controls were heterozygous (dosage 1), giving complete genotype–phenotype separation (cases 118/118; controls 4/4). SNP positions (FCAr2.3 coordinates) are shown on the x-axis. (D) Maximum-likelihood phylogeny of the R2R3 DNA-binding domain of *MYB101-like* (Fca_r2.3chr04_g020650.1; labelled FcMYB101-like), with 56 representative *Arabidopsis thaliana* R2R3-MYB members spanning the major subclades and 11 previously characterised MYB101/GAMYB reference sequences from six other species. The tree was inferred using IQ-TREE with the best-fit substitution model selected by ModelFinder; node labels show SH-aLRT (%)/ultrafast bootstrap (%) where both are ≥70%. *MYB101-like* (red) falls within a well-supported MYB101 subclade (blue), separated from the other *Arabidopsis* R2R3-MYB subfamilies sampled here (grey). The tree is unrooted; basal branching order should not be interpreted as ancestral. Scale bar, substitutions per site.

**Figure 4.**
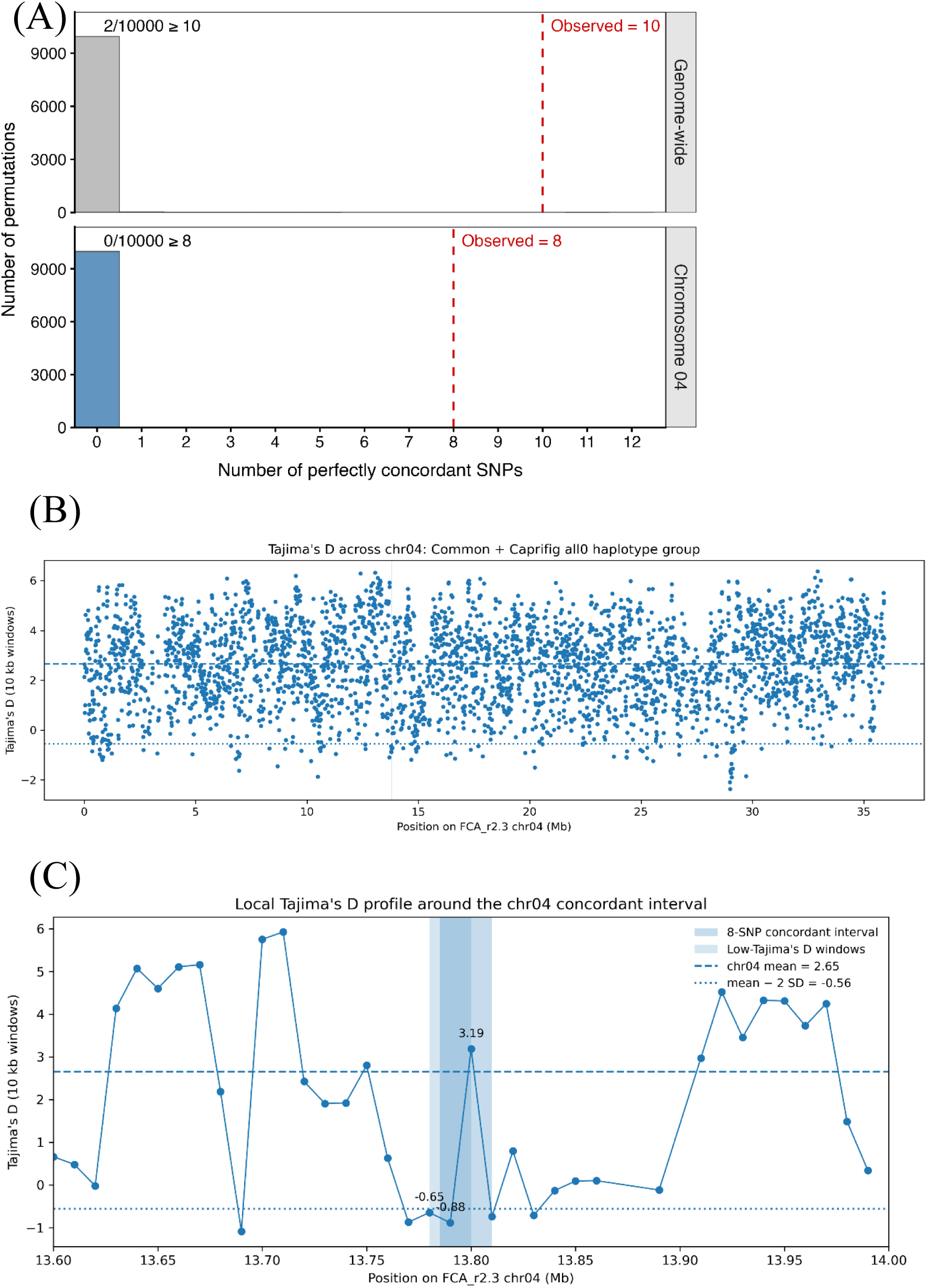
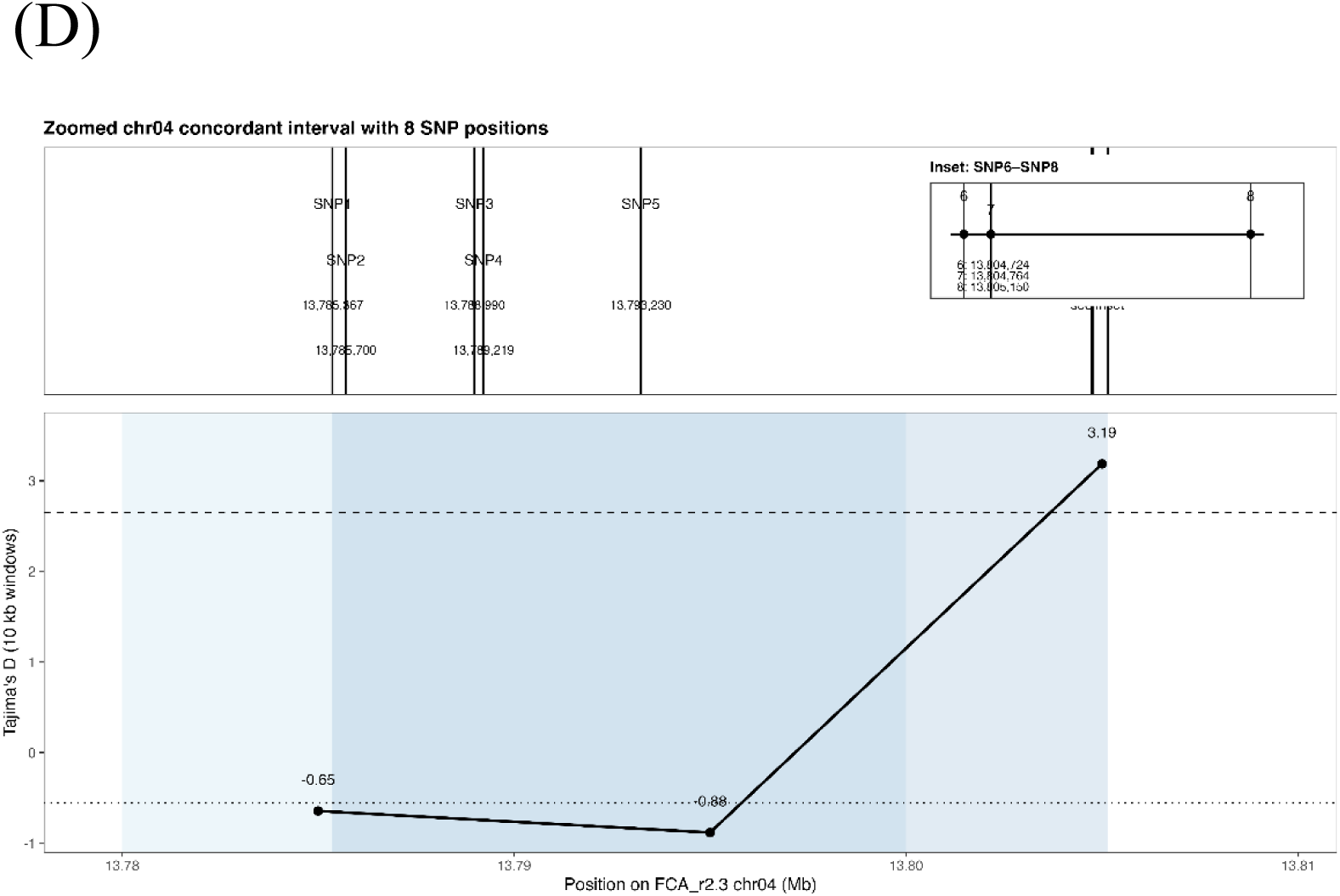
Statistical validation and selective-sweep signature of the *Eden* locus concordant haplotype. (A) Permutation test for the chromosome 04 *MYB101-like*-centred concordant haplotype. Phenotype labels were permuted 10,000 times at the observed case–control ratio (118 non-Smyrna cases : 4 Smyrna-type controls), counting perfectly concordant SNPs on chromosome 04 (blue, lower panel) and genome-wide (grey, upper panel). Red dashed lines, observed values (chr04, 8 SNPs; genome-wide, 10 SNPs); empirical P = 1.0 × 10⁻⁴ and 3.0 × 10⁻⁴, respectively. (B) Tajima’s D across chromosome 04 in Common-type and Caprifig accessions homozygous for the d-type haplotype, in non-overlapping 10 kb windows (n = 3,196). Each point is one window; dashed line, chr04-wide mean (2.65); dotted line, genome-wide mean − 2 SD (−0.405); vertical line, left boundary of the 8-SNP concordant interval (13.785 Mb). (C) Local Tajima’s D profile across 13.60–14.00 Mb; dark blue shading, the 8-SNP concordant interval (13.785–13.810 Mb); lines as in (B), with mean − 2 SD = −0.56. Windows at 13.780 and 13.790 Mb show values of −0.645 and −0.884, the latter in the bottom 0.90th percentile genome-wide (n = 28,202 windows; 2.726 ± 1.566; Z = −2.31). The elevated value at 13.800 Mb (3.19) lies in the repeat-rich 2-alkenal reductase sub-region (g020660.1), where outgroup calls at SNP6 and SNP7 were ambiguous (Supplementary Table 3). (D) The eight concordant SNPs (chr04:13,785,367–13,805,150) defining the *Eden* haplotype, on the same physical scale as the profile from (C) reproduced below. SNP6–SNP8, spanning only 426 bp, are expanded in the inset.

The 8 concordant SNPs span a 13.785–13.810 Mb interval containing three gene models: g020640.1 (no functional annotation available), *MYB101-like* (g020650.1), and 2-alkenal reductase (g020660.1) (Figure 3B; Supplementary Fig. 4C). Of the two annotated genes, *FcMYB101-like* (hereafter *MYB101-like*) is the most plausible candidate: MYB101 belongs to the GAMYB subfamily of R2R3-MYB transcription factors, which regulate reproductive development and hormone signalling associated with parthenocarpy in other crops (Figure 3D). Read-based inspection of the *MYB101-like* gene body in the Smyrna-type control Fc_57 (Sari Lop; 38.9× depth) identified two exonic SNPs, two splice-proximal SNPs, and one intronic INDEL (Supplementary Fig. 4F), none of which showed panel-wide concordance. The eight concordant SNPs are instead predominantly non-coding, including one 4.1 kb upstream of *MYB101-like*, indicating a haplotype defined principally by potentially cis-regulatory variation.

Phylogenetic analysis of the R2R3 DNA-binding domain (56 *Arabidopsis* R2R3-MYB members plus curated MYB101/GAMYB reference sequences) placed *MYB101-like* within a well-supported MYB101 subclade (Figure 3D), grouping with *Morus notabilis* MnMYB101 (SH-aLRT/UFBoot = 83.8/80) and then with *Arabidopsis* AtMYB101 and *Citrus sinensis* CsMYB101 (83/77); support for uniting this subclade with GAMYB sensu stricto genes was weak (47.3/57). *MYB101-like* is therefore a bona fide MYB101-subclade member functionally related to GAMYB, which regulates ovule development and fruit set in tomato^22^ and gibberellin-responsive reproductive development more broadly^23^; GA treatment induces parthenocarpic fruit set in San Pedro-type fig^24^.

The chromosome 04 *MYB101-like* concordant haplotype showed no co-localisation with any GWAS or TWAS locus for fruit quality traits, indicating that parthenocarpy and fruit quality diversification are governed by architecturally distinct genomic layers.

### Fruit flavour in fig is organised as an aroma-dominated integrated sensory system

To characterise the integrated architecture of fruit flavour variation, we applied Multi-Omics Factor Analysis (MOFA+) across four omics layers—volatile aroma profiles (549 features), sugar–acid composition (41 parameters), taste sensor measurements (8 dimensions), and RNA expression (45,900 genes)—in 81 accessions; colorimetric and anthocyanin traits were excluded as they reflect visual rather than flavour quality and are addressed separately by GWAS and TWAS. The model converged on 15 biologically interpretable factors. RNA expression contributed the largest fraction of total explained variance (41.0%), followed by volatile aroma (25.4%), sugar–acid composition (18.5%), and taste sensor profiles (15.1%) (Figure 5A; Supplementary Fig. 6); the transcriptomic contribution likely reflects that layer’s higher dimensionality rather than greater biological influence on sensory output. Among the layers directly reflecting sensory output, aroma contributed the largest share (43.1% of the variance explained by the three sensory layers combined), establishing volatile aroma as the primary axis of flavour differentiation across accessions.

**Figure 5.**
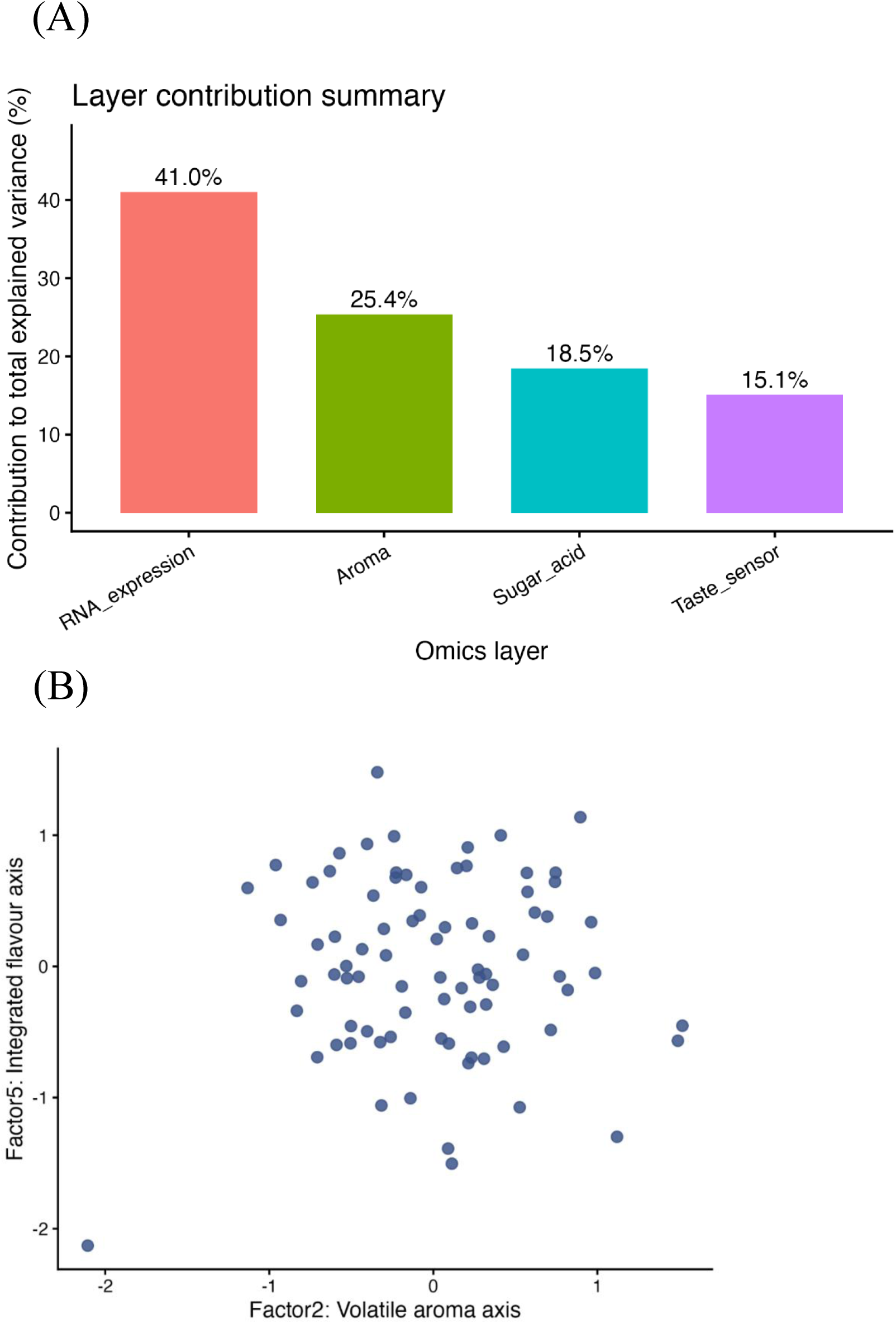
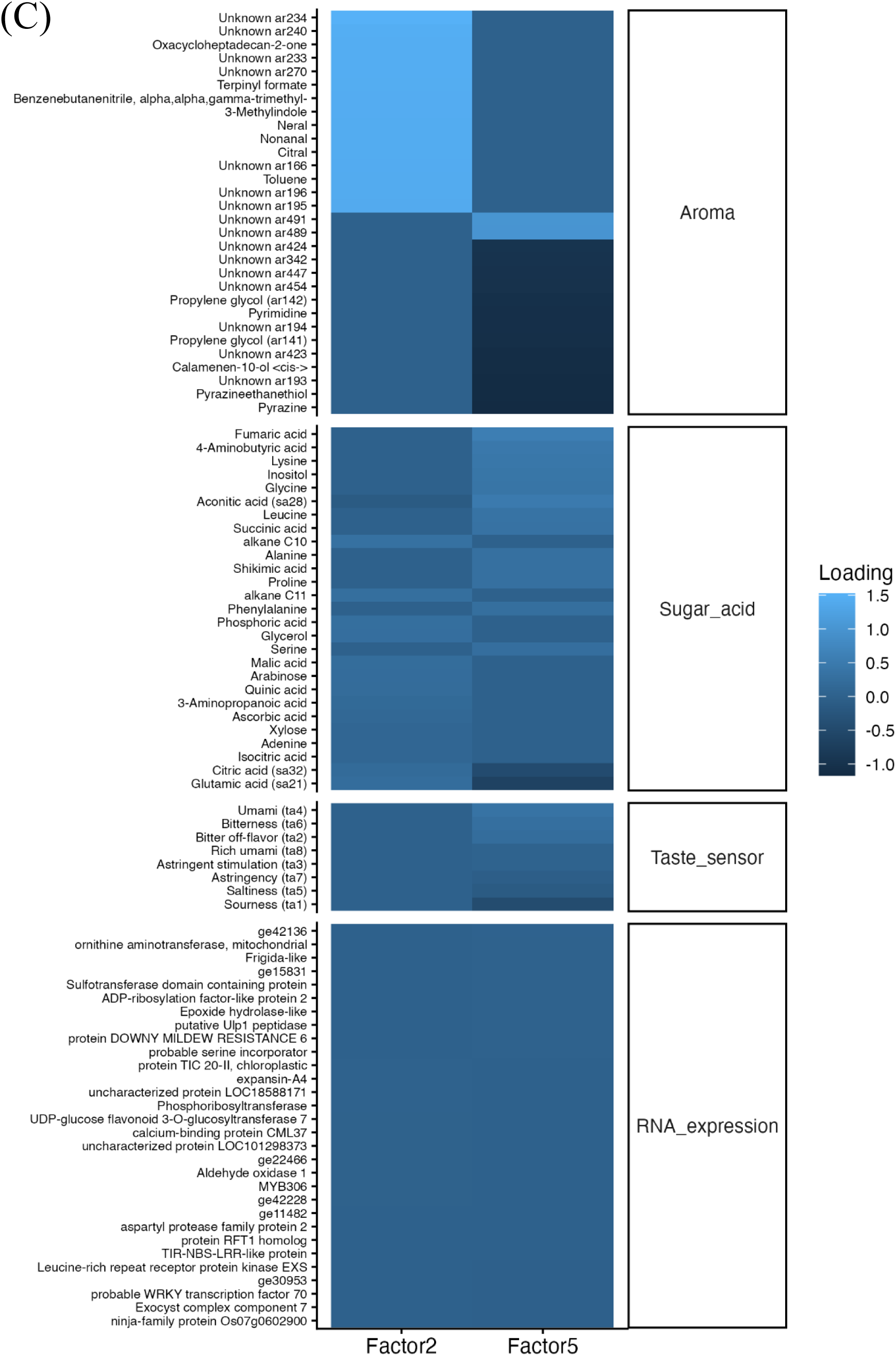
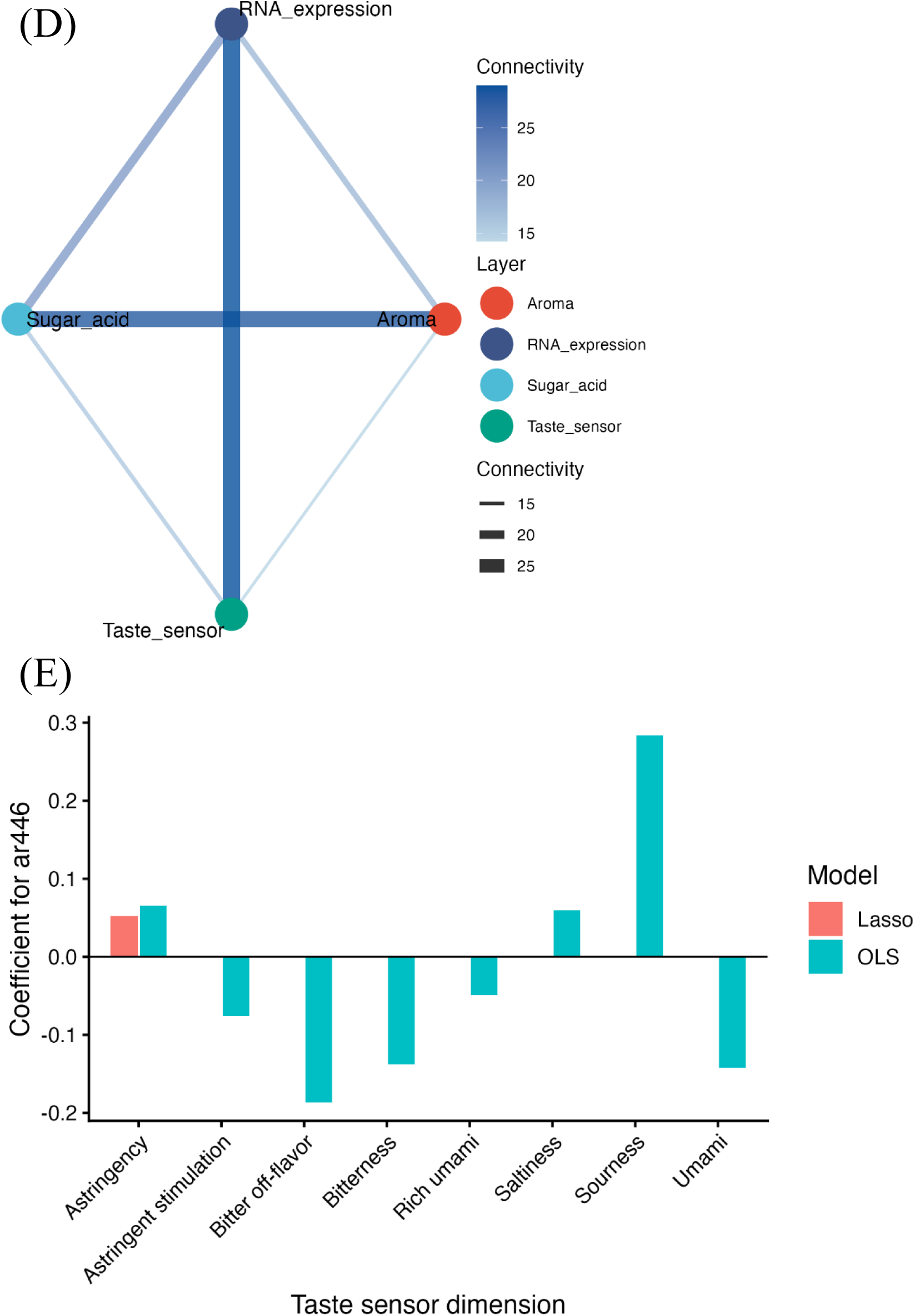
Multi-omics factor analysis reveals an aroma-dominated integrated sensory architecture underlying fig fruit flavour. (A) Total variance explained by MOFA+ for each of the four omics layers: RNA expression 41.0%, volatile aroma 25.4%, sugar–acid composition 18.5%, and taste sensor profiles 15.1%. Among the three sensory layers, aroma accounted for 43.1% of their combined explained variance. (B) Accession scores on Factor 2 (Volatile Aroma Axis; x-axis) and Factor 5 (Integrated Flavour Axis; y-axis) for 81 accessions. Factor 2 was dominated by volatile aroma features (18.3% of total explained variance); Factor 5 integrated volatile aroma (7.2%), sugar–acid composition (4.5%), and taste sensor profiles (3.3%) (Supplementary Fig. 6). (C) Heatmap of factor loadings for the top contributing features of Factors 2 and 5. For Factor 2, volatile aroma features showed strong positive loadings, with citrus- and terpene-associated compounds prominent; sugar–acid features showed moderate loadings, and taste sensor and RNA expression features near-zero loadings. For Factor 5, loadings were distributed across the volatile aroma, sugar–acid, and taste sensor layers. (D) Network diagram of inter-layer connectivity through shared MOFA+ latent factors. Edge width is proportional to the strength of shared latent factor associations between omics layer pairs. All four layers were interconnected. (E) Lasso and ordinary least squares (OLS) regression coefficients for the volatile feature ar446 against each of the eight taste sensor dimensions. Under Lasso regularisation, ar446 was the sole selected predictor of astringency (positive coefficient); under OLS it showed positive associations with sourness and astringency and negative associations with bitter off-flavor, bitterness, and umami.

Factor 2 was dominated by volatile aroma features, defining the primary Volatile Aroma Axis along which accessions with the greatest aroma diversity are distributed (Figure 5B, 5C). Factor 5 integrated volatile aroma, sugar–acid composition, and taste sensor profiles simultaneously, defining an Integrated Flavour Axis that captures coordinated variation across sensory layers. Sugar–acid parameters were associated with both axes: organic acids and amino acids loaded on Factor 2, indicating that aroma variation is coupled with primary metabolite composition, whereas citric acid and glutamic acid loaded more strongly on Factor 5. Network-based analysis showed all four omics layers interconnected through shared latent factors, indicating coordinated rather than independent variation (Figure 5D). This structure was independently supported by correlation network analysis using ConfeitoGUI^25^: among 86 communities spanning multiple layers, the majority linked features from different omics layers rather than forming layer-specific modules (Supplementary Fig. 9). Together, these patterns support a model in which fig flavour is organised as an integrated sensory system with volatile aroma as the dominant axis of differentiation among sensory layers.

The volatile feature ar446 showed consistent regression coefficients across multiple modelling approaches including MOFA+, Lasso, and OLS regression (Figure 5E). Under Lasso regularisation it was selected as the sole predictor of astringency (positive coefficient); under OLS it showed the strongest positive association with sourness and a positive association with astringency, while showing negative associations with bitter off-flavor, bitterness, and umami. ar446 therefore functions as a taste-attribute-specific cross-modal modulator, consistently enhancing sourness and astringency while reducing bitter off-flavor and bitterness. Because the negative association with umami reflects suppression of a generally pleasant rather than an aversive attribute, these effects represent a reshaping of the overall taste balance rather than simple suppression of aversive tastes, consistent with retronasal integration acting independently of absolute concentration^26^.

Fruit flavour in fig is thus not a collection of independent metabolic traits but an organised sensory system in which volatile aroma constitutes the dominant axis of differentiation among sensory layers, with aroma–taste cross-modal interaction (exemplified by ar446) and coupling with sugar–acid balance as additional components of the integrated structure.

### Genetic architecture of individual fruit quality traits within the sensory system

#### Genetic basis of primary metabolite composition: GWAS and TWAS identify regulators of sugar accumulation and amino acid diversity

GWAS identified g048100.1 (chr04) as the primary locus for amino acid composition, associated with eight amino acids (PP = 0.64–0.99), and g012770.1 (chr02) associated with xylose (PP = 0.99) and asparagine (PP = 0.53) (Figure 6A). Neither candidate has a functional annotation that directly explains its broad multi-trait signal—g012770.1 encodes a disease-resistance protein of unclear relevance to sugar and amino acid metabolism—suggesting that both act as upstream regulators rather than pathway-specific enzymes. Notably, the chr04 signal for g048100.1 maps to an interval distinct from the parthenocarpy candidate locus, confirming that primary metabolite composition and reproductive mode are controlled by non-overlapping genomic regions on the same chromosome (Supplementary Fig. 5A).

**Figure 6.**
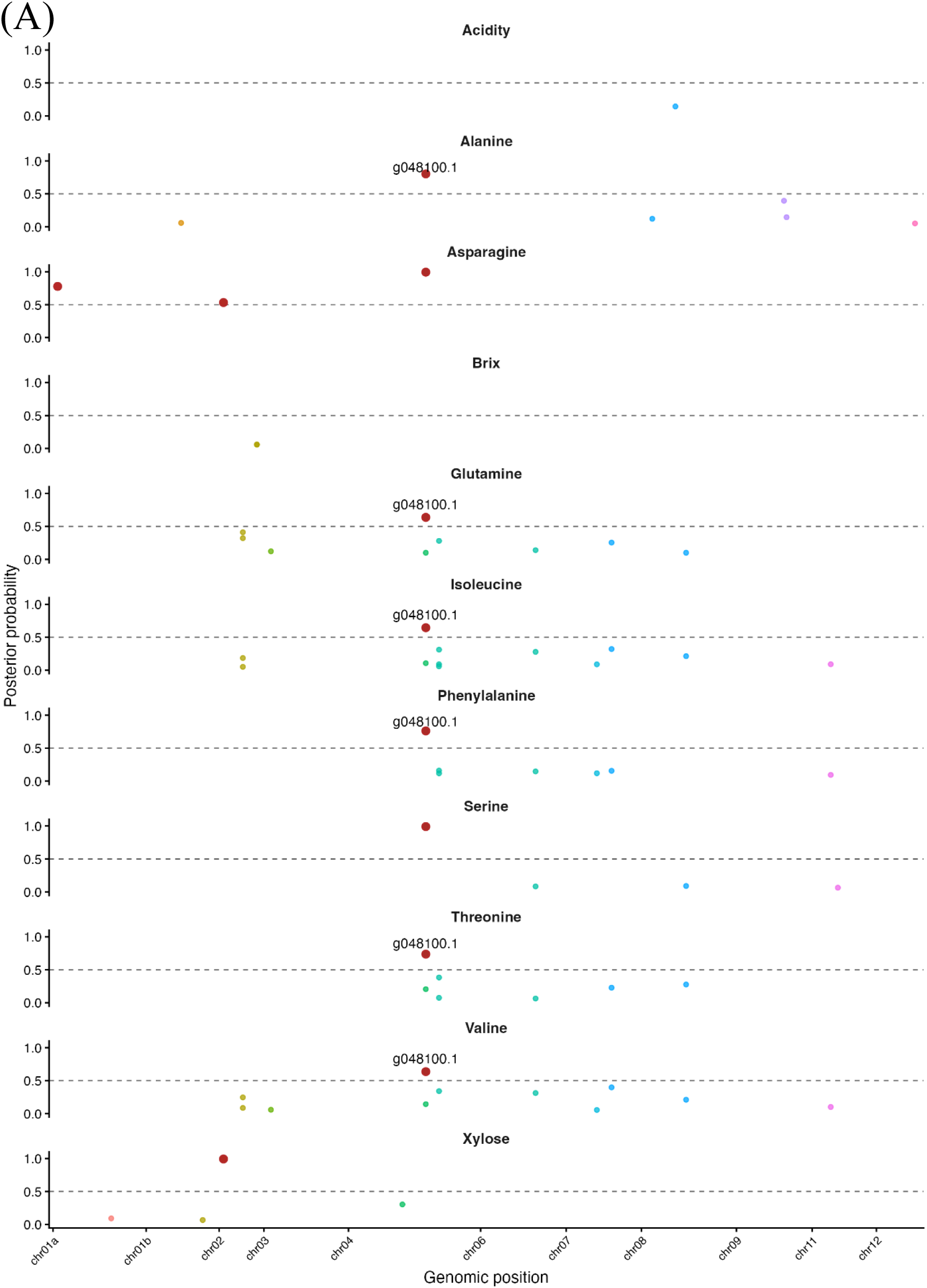
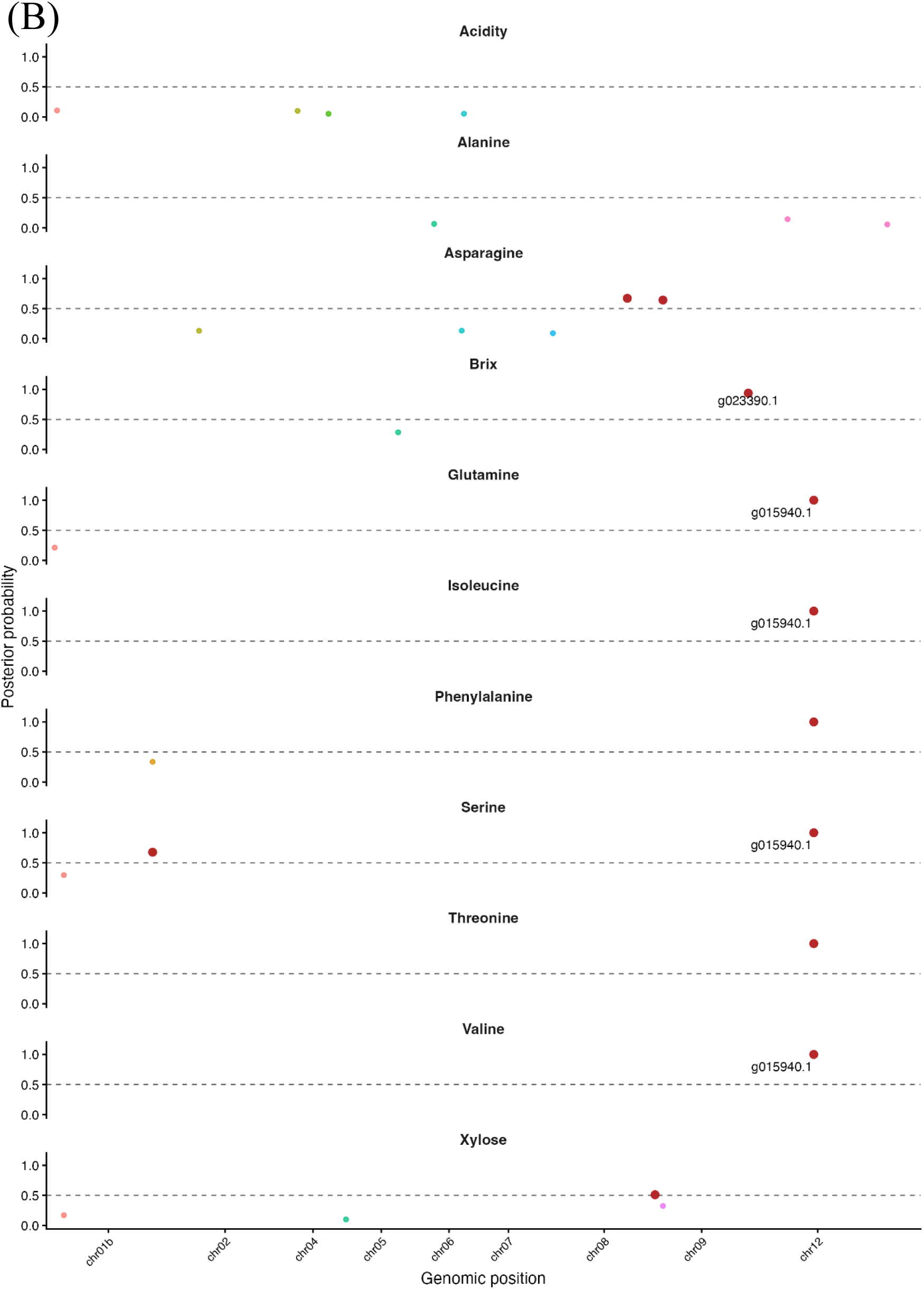
Genetic basis of primary metabolite composition: GWAS and TWAS identify candidate regulators of sugar accumulation and amino acid diversity. (A) Posterior probability plot from Bayesian GWAS for representative primary metabolite traits, including xylose and selected amino acids. Candidate loci with posterior probability (PP) > 0.5 are highlighted. A chromosome 04 locus corresponding to g048100.1 showed high-confidence associations with eight amino acid traits (alanine, asparagine, glutamine, isoleucine, phenylalanine, serine, threonine, and valine; PP = 0.64–0.99). A chromosome 02 locus corresponding to g012770.1 (Disease resistance protein; Data file 2) was associated with xylose (PP = 0.99) and asparagine (PP = 0.53). The dashed line indicates the PP = 0.5 threshold. (B) Transcriptome-wide posterior probability plot from TWAS for the same representative traits. Candidate genes with PP > 0.5 are highlighted. The leading candidates are g023390.1 (phosphoglycerate mutase-like protein; Brix, PP = 0.937) and g015940.1 (geraniol 8-hydroxylase; glutamine, isoleucine, phenylalanine, serine, threonine, valine PP = 1.000; 2-aminoadipic acid PP = 0.802). A third TWAS candidate, g024230.1 (Embryogenesis-associated protein), was associated with xylose (PP = 0.510; chr08).

TWAS identified g023390.1 (chr09, phosphoglycerate mutase-like) as the leading transcriptional regulator of sugar accumulation (Brix; PP = 0.937), consistent with a role in glycolytic carbon partitioning^27^. g015940.1 (chr12, geraniol 8-hydroxylase) showed strong associations (PP = 1.00) across six amino acids plus 2-aminoadipic acid (PP = 0.80)—substantially overlapping the GWAS amino-acid signal at g048100.1 and indicating that both structural variation and transcriptional regulation shape amino acid diversity. A third candidate, g024230.1 (chr08), was associated with xylose (PP = 0.51), complementing the GWAS signal at a distinct locus (Figure 6B, S5B; Supplementary Table 1).

Within the MOFA+ sensory architecture, sugar–acid parameters (fructose, glucose, sucrose) loaded as secondary modulatory axes coupled to the aroma-dominated Factor 2 and the Integrated Flavour Axis (Factor 5), positioning g023390.1 in particular as a genomic node linking primary metabolite composition to the broader flavour system.

### Genetic basis of secondary metabolite diversity: transporter and signaling networks drive aroma variation and glycosylation controls anthocyanin pigmentation

Aroma compounds and anthocyanin pigments are the two dominant secondary-metabolite classes shaping fig sensory and visual quality; we performed integrated GWAS and TWAS for both trait classes.

For aroma, GWAS identified three chromosomal hotspots (PP > 0.5; Figure 7A): chr01b, a candidate-rich region for diverse volatiles; chr05, anchored by pleiotropic hub locus g025980.1 (PP = 1.0 for 1-Pentanol; PP > 0.98 across 12 further compounds); and a secondary chr10 region. TWAS identified 84 candidate genes across 115 gene–trait associations, dominated by transporters (n = 27) and kinase/signalling genes (n = 19), followed by oxidoreductases (n = 7) and lipid/acyl/esterase genes (n = 5)—the latter mechanistically linked to fatty-acid-derived volatile precursor supply^28,29^ (Figure 7B). GWAS–TWAS integration identified three co-localised regions within ±100 kb, on chr01b (11.78–11.83 Mb) and at two chr05 loci, indicating that SNP variation partly acts through local expression changes (Supplementary Fig. 7; Supplementary Data 2).

**Figure 7.**
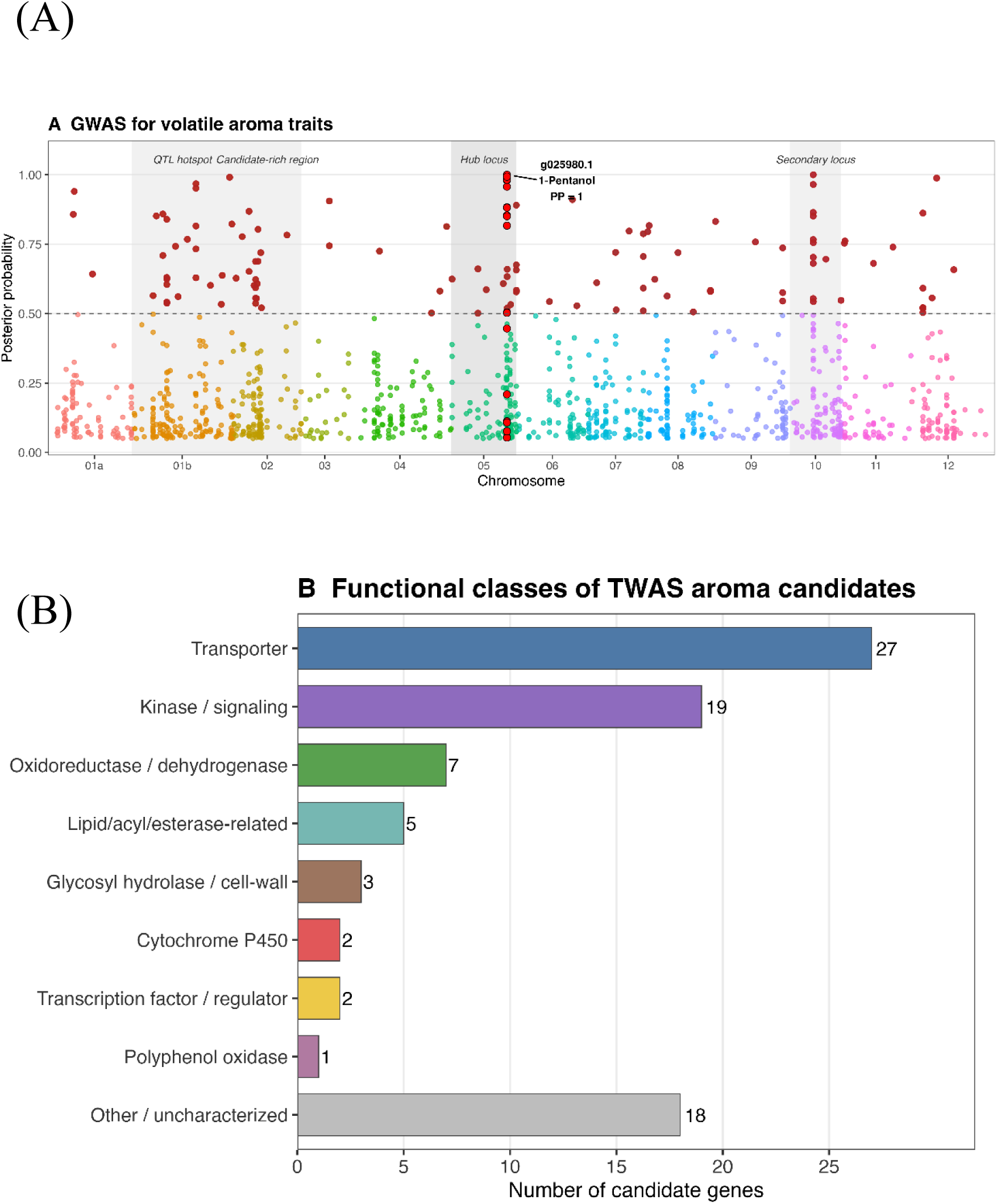
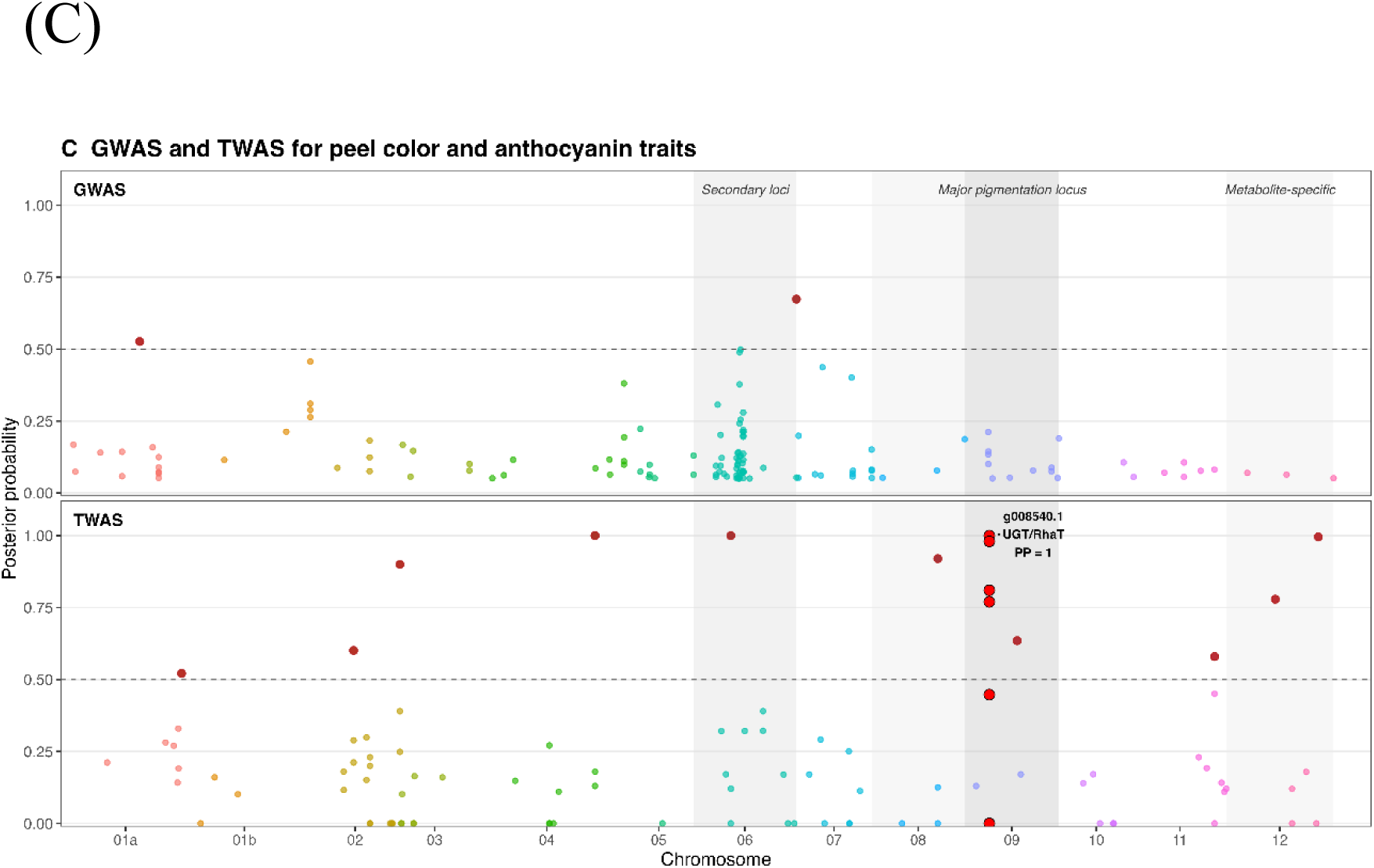
Genetic basis of secondary metabolite diversity: transporter and signaling networks drive aroma variation, whereas anthocyanin glycosylation controls peel pigmentation. (A) Genome-wide posterior probability plot from Bayesian GWAS for volatile aroma traits. Candidates with PP ≥ 0.5 are shown in red; the dashed horizontal line marks PP = 0.5. Three regions are shaded: a QTL hotspot and candidate-rich region on chromosome 01b, a hub locus on chromosome 05, and a secondary locus on chromosome 10. The pleiotropic chromosome 05 candidate g025980.1—associated with 1-Pentanol (PP = 1.0) and 12 additional volatile compounds—is labelled. (B) Functional classification of 84 unique TWAS candidate genes for volatile aroma traits (PP > 0.5; 115 gene–trait associations), assigned from product annotations and KEGG. Transporter-related genes (n = 27) and kinase/signalling components (n = 19) were the largest categories, followed by oxidoreductases/dehydrogenases (n = 7), lipid/acyl/esterase-related genes (n = 5), glycosyl hydrolases and cell-wall-related genes (n = 3), cytochrome P450s (n = 2), transcription factors/regulators (n = 2), polyphenol oxidases (n = 1), and other or uncharacterised genes (n = 18). (C) Genome-wide posterior probability plots from GWAS (upper) and TWAS (lower) for peel colorimetric and anthocyanin traits; candidates with PP ≥ 0.5 are shown in red. Three regions are indicated: secondary loci (chr05–06), the major pigmentation locus (chr09), and metabolite-specific loci (chr11–12). The chromosome 09 candidate g008540.1, encoding a UDP-rhamnose:anthocyanidin-3-glucoside rhamnosyltransferase (UGT/RhaT), was the dominant TWAS candidate for peel colorimetric traits (Munsell colour index PP = 1.0; a* PP = 0.98; b* PP = 0.81; L* PP = 0.77). GWAS identified no high-confidence candidate at the chromosome 09 locus; a secondary GWAS candidate region was detected on chromosome 06 (maximum PP ≈ 0.67).

For peel colour and anthocyanins, TWAS identified a single dominant locus—UDP-rhamnose:anthocyanidin-3-glucoside rhamnosyltransferase g008540.1 (chr09)—with PP = 1.0 for Munsell index and consistently high PP across colorimetric parameters (a* = 0.98, b* = 0.81, L* = 0.77); GWAS detected no strong candidate at this locus (secondary chr06 signal, PP ≈ 0.67), indicating that the association is captured primarily at the expression level (Figure 7C). Fruit colouring ratio and individual anthocyanin species mapped to distinct loci, including kinase and glycosyl transferase candidates (PP = 1.00) and, for P3R, geraniol 8-hydroxylase g015940.1 and NAC25 g020600.1 (Supplementary Data 2). Together these results point to g008540.1-centred glycosylation as the primary driver of pigmentation, modulated by a broader network of ion, redox, and transcriptional regulators.

Aroma and anthocyanin architectures share a common principle: a small number of high-impact hub loci—transporter and signalling genes for aroma, glycosylation regulators for anthocyanins—govern broad trait classes, while trait-specific loci modulate individual compounds. Within the MOFA+ framework, the chr01b and chr05 aroma hotspots and the predominance of transporter and signalling gene expression represent the genomic entry points into the aroma-dominated flavour system.

### Genetic basis of taste attributes and agronomic quality traits: defence-palatability antagonism in taste and multi-layer control of fruit morphology

Taste attributes are the sensory layer most directly experienced by consumers, while fruit morphology defines marketability and postharvest performance; we applied integrated GWAS and TWAS to both trait categories.

For taste, GWAS identified only suggestive loci (PP < 0.5), including TIR-NBS-LRR-like protein g010380.1 (chr08, bitterness) and receptor-like kinases—gene families typically linked to pathogen resistance^30,31^ (Figure 8A). TWAS identified stronger candidates (PP > 0.5), led by g012990.1 (chr11, acidity, PP = 0.99) and g013910.1 (chr07, astringent stimulus, PP = 0.91) (Supplementary Table 2). Two hotspots emerged: on chr03 (14.3–15.7 Mb), GWAS candidates for bitter acidity (g020730.1) and astringency co-occurred with TWAS candidate vinorine synthase g022920.1 (Supplementary Fig. 8A); on chr08 (∼7.6–8.4 Mb), a GWAS bitterness candidate (g010380.1) and a TWAS full-bodied-flavour candidate (g011520.1) co-localised, suggesting a region controlling both aversive and favourable taste dimensions (Supplementary Fig. 8B). Aversive-taste genes were predominantly defence-related, whereas favourable-attribute genes were enriched for transposon-related and uncharacterised genes (Figure 8B)—a defence–palatability antagonism suggesting that domestication relaxed selection on defensive biosynthesis^13^, leaving favourable-flavour genetics largely uncharacterised. These attributes map onto the MOFA+ Factor 5 Integrated Flavour Axis.

**Figure 8.**
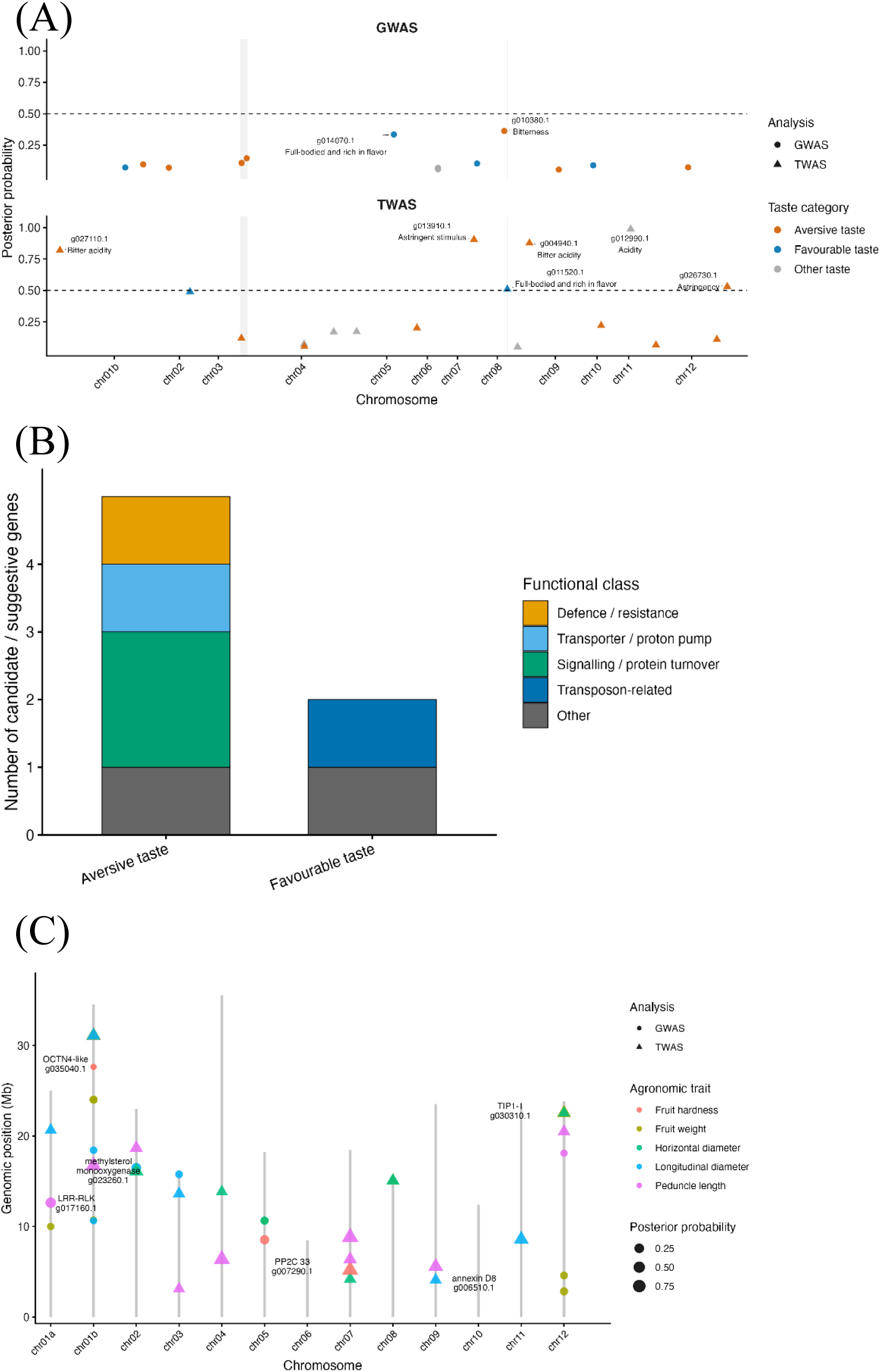
Genetic basis of taste attributes and agronomic quality traits: defence–palatability antagonism and multi-layer control of fruit morphology. (A) Genome-wide posterior probability plots from GWAS (upper) and TWAS (lower) for taste sensor attributes. Loci are colour-coded by taste category: aversive taste attributes (orange; bitterness, bitter acidity, astringency, astringent stimulus), favourable taste attributes (blue; full-bodied and rich flavour), and other taste attributes (grey; acidity). The dashed horizontal line indicates a posterior probability of PP = 0.5. High-confidence TWAS candidates (PP > 0.5) include g027110.1 (chr01a; Bitter acidity), g013910.1 (chr07; Astringent stimulus), g004940.1 (chr09; Bitter acidity), g012990.1 (chr11; Acidity), g011520.1 (chr08; Full-bodied and rich in flavor), and g026730.1 (chr12; Astringency). A candidate-rich region on chromosome 03 (14.3–15.7 Mb) and a candidate region on chromosome 08 (7.6–8.5 Mb) are highlighted. (B) Functional classification of candidate and suggestive GWAS/TWAS genes associated with aversive and favourable taste attributes. Aversive taste-associated candidates (n = 5) included genes in the following functional classes: defence/resistance (n = 1), transporter/proton pump (n = 1), signalling/protein turnover (n = 2), and other/uncharacterised (n = 1). Favourable taste-associated candidates (n = 2) included transposon-related (n = 1) and other/uncharacterised (n = 1) genes. (C) Summary diagram of GWAS and TWAS candidate loci for agronomic quality traits, including fruit weight, longitudinal and horizontal diameter, peduncle length, and fruit hardness. Circle symbols represent GWAS candidates; triangle symbols represent TWAS candidates. Symbol size is proportional to posterior probability. Candidate genes are labelled with gene ID and functional annotation where available, including annexin D8 g006510.1 (chr09; fruit weight GWAS), aquaporin TIP1-1 g030310.1 (chr12; fruit weight TWAS), methylsterol monooxygenase g023260.1 (chr01b; longitudinal diameter TWAS), protein phosphatase 2C 33 g007290.1 (chr07; fruit hardness TWAS), OCTN4-like transporter g035040.1 (chr01b; fruit hardness GWAS), and LRR receptor-like kinase g017160.1 (chr01a; peduncle length GWAS).

For agronomic traits (Figure 8C; Supplementary Table 2), fruit weight was linked to annexin D8 g006510.1 (chr09, GWAS) and aquaporin TIP1-1 g030310.1 (chr12, PP = 0.53, TWAS), implicating membrane and water transport^32^. Longitudinal diameter mapped to multiple TWAS candidates—g012210.1 (chr07, PP = 0.96, RGA3-like), g009410.1 (chr04, PP = 0.90, PI-PLC X domain), and g021570.1 (chr01b, PP = 0.80, TBC1 domain)—reflecting polygenic control; g008540.1, the anthocyanin hub locus, also associated with longitudinal diameter (PP = 0.69), indicating pleiotropy across secondary-metabolic and morphological traits. Peduncle length was led by LRR receptor-like kinase g017160.1 (chr01a, PP = 0.52; GWAS), and fruit hardness by protein phosphatase 2C 33 g007290.1 (chr07, PP = 0.89; TWAS), consistent with ABA-mediated cell-wall regulation^33^, and OCTN4-like transporter g035040.1 (chr01b; GWAS).

Across agronomic traits, GWAS and TWAS identify largely non-overlapping candidates spanning structural variation, transcriptional regulation, hormone signalling, and membrane dynamics—consistent with the polygenic architecture reported for morphological traits in other domesticated fruit crops^34^—and operate largely independently of the aroma-dominated flavour system.

## Discussion

The results presented here converge on two principal findings. First, fruit flavour in *Ficus carica* is not a collection of independent metabolic traits but an organised sensory system in which aroma constitutes the dominant axis of variation, modulating taste perception through cross-modal interaction and coupling with sugar–acid balance as a secondary layer—consistent with multi-omics dissections of flavour in other fruit crops, in which volatile-focused GWAS integrated with eQTL and structural-variant data have similarly resolved flavour into interacting molecular layers^35^. Second, the genetic architecture underlying this sensory system is entirely distinct from the chromosome 04 concordant haplotype associated with parthenocarpy: polygenic loci controlling sugars, organic acids, aroma compounds, anthocyanins, and taste attributes show no co-localisation with the *MYB101-like*-centred candidate interval (Figure 9). This near-Mendelian, single-locus architecture contrasts with the hormone-signalling networks underlying parthenocarpy in other crop lineages, including cucurbits and tomato, where a linked two-gene module (TSP4a/TSP4b) governing auxin- and AINTEGUMENTA-mediated fruit set has recently been characterised^36,37^, indicating that the genetic route to pollinator-independent fruit set is not uniform across angiosperms. Fig domestication therefore involved at least two genetically separable dimensions—reproductive innovation and sensory diversification—that proceeded under distinct selective regimes.

**Figure 9.**
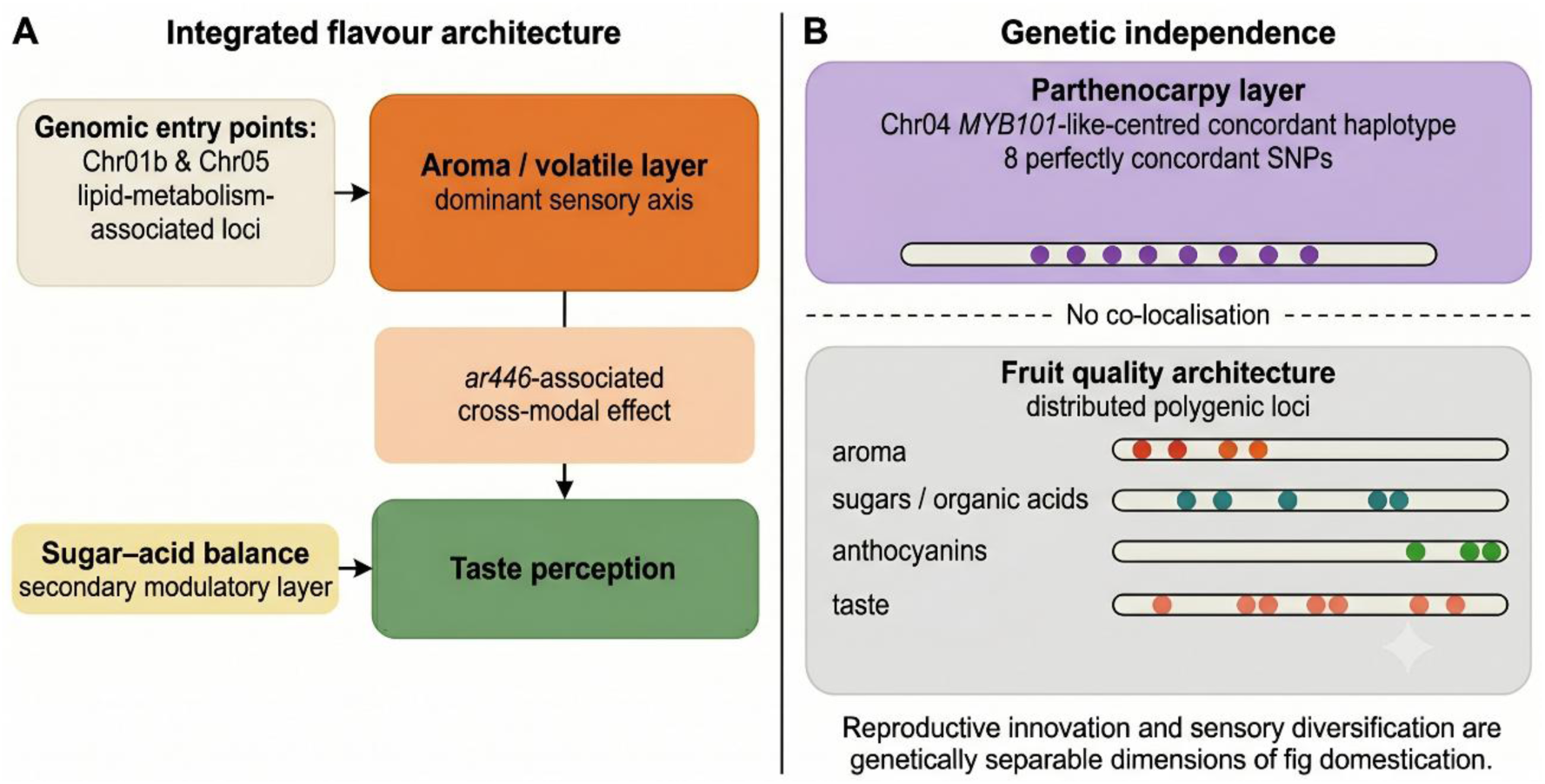
Integrated model of fig fruit flavour architecture and the genetic independence of parthenocarpy from fruit quality. Schematic overview of the two principal findings of this study. (A) Integrated flavour architecture inferred from multi-omics factor analysis (MOFA+) across volatile aroma, sugar–acid, taste-sensor, and RNA-expression layers (Figure 5). The aroma/volatile layer represents the dominant axis of flavour variation and is linked to taste perception through cross-modal effects, including an association with the volatile feature ar446. Sugar–acid balance contributes to taste perception as a coupled component of the integrated system. (B) Genetic independence of parthenocarpy from fruit-quality architecture. A chromosome 04 MYB101-like-centred concordant haplotype, defined by eight perfectly concordant SNPs that distinguish non-Smyrna and Smyrna-type accessions, is shown as a parthenocarpy-associated genomic layer distinct from the distributed polygenic loci controlling fruit-quality traits, including sugars, organic acids, aroma compounds, anthocyanins, and taste attributes. The absence of co-localisation between this chromosome 04 parthenocarpy-associated interval and fruit-quality loci supports a model in which reproductive innovation and sensory diversification represent separable dimensions of fig domestication shaped by distinct selective regimes. Outgroup *Ficus* species outside subsect. *Ficus* carry alleles corresponding to the Smyrna-type (e-type) haplotype at these positions, supporting a model in which the parthenocarpic (d-type) haplotype was driven to fixation in cultivated *F. carica* by strong positive selection during domestication. *F. palmata*, as the sister species of *F. carica* within subsect. *Ficus*, also carries this haplotype—predominantly homozygous at the eight diagnostic SNPs but heterozygous across roughly one quarter of the broader *Eden* interval, a pattern more consistent with long-term maintenance of ancestral polymorphism than with recent gene flow—and therefore does not serve as an independent outgroup for polarity inference.

Among sensory omics layers, aroma accounts for 43.1% of the variance explained by sensory layers combined, consistent with the psychophysical primacy of retronasal olfaction in flavour perception^26,38,39^ and with aroma’s role as a cross-modal modulator of taste^40^. The disproportionate influence of the single volatile feature ar446 shows that this aroma–taste interaction is mediated by discrete high-impact compounds rather than being diffuse. At the genetic level, the concentration of aroma QTLs on chromosomes 01b and 05 and the predominance of transporter and signalling TWAS candidates indicate that substrate transport, signal transduction, and fatty-acid-derived precursor supply are the primary levers linking genotype to flavour, supporting the case that flavour-improvement breeding should prioritise this aroma entry point rather than optimising individual metabolite classes in isolation (Figure 9).

The genetic independence of parthenocarpy from fruit-quality loci supports a model of distinct selection pressures acting at different stages or in different genetic backgrounds. The chromosome 04 *MYB101-like* concordant haplotype (8 SNPs, 13.785–13.810 Mb, complete genotype–phenotype separation across 122 accessions, empirical P = 0.0001) represents a qualitatively different class of genomic signal from the distributed polygenic architecture of sensory traits, consistent with discrete major-effect regulation by a transcription factor with established roles in reproductive development and hormone signalling. The 25 kb interval contains two other gene models: an uncharacterised gene (g020640.1) and 2-alkenal reductase (g020660.1). The two SNP positions closest to the latter (S6, S7) fall within a repeat-rich subregion and were excluded from the primary polarity analysis (Supplementary Table 3); their available calls show the opposite pattern to the other six positions, with most outgroup species carrying the derived (d-type) rather than the ancestral (e-type) allele. This argues against the 2-alkenal reductase flanking region sharing the domestication-associated selective signal seen across the rest of the haplotype, disfavouring it relative to *MYB101-like*. Fine-mapping in recombinant individuals will be required to confirm candidate gene identity. We propose the locus symbol *Eden*—reflecting the entomophilous (e) and domestication-associated (d) allele states—for this chromosome 04 interval, with alleles designated *Eden-e* (ancestral, associated with obligate pollination dependence) and *Eden-d* (derived, required in homozygous state for parthenocarpic fruit set), following the convention established for other *F. carica* reproductive trait loci^2,41^. *FcMYB101-like* is proposed as the leading positional candidate gene underlying the *Eden* locus, pending phylogenetic resolution of its precise orthology and functional validation.

A recent comparative genomic analysis linked contraction of the s-domain receptor-like kinase (RLK) family, which mediates pollen recognition, to parthenocarpy in *F. carica*^7^. Being a species-level feature, this contraction cannot explain the segregation of pollination dependence within *F. carica*, and no s-domain RLK lies within 1 Mb of *Eden* (Supplementary Data 3). The two may act at different levels, with reduced pollen recognition providing a permissive background for allelic variation at *Eden*.

All four Smyrna-type accessions were heterozygous at each of the 8 concordant positions rather than homozygous for the alternative allele, indicating that they carry one copy of the parthenocarpic d-type haplotype alongside a divergent e-type haplotype matching outgroup *Ficus* species. Parthenocarpic fruit set therefore requires d-type homozygosity, a single e-type copy being sufficient to confer pollination dependence—consistent with the classical model of a single dominant parthenocarpy mutation, P^1,2,6^. The sweep fixing the d-type haplotype in Common-type, Caprifig, and San Pedro accessions thus required a transition from d/e heterozygosity, retained in Smyrna, to d/d homozygosity, likely facilitated by fig’s clonal propagation system.

Tajima’s D analysis at the *Eden* locus (10 kb windows, Common-type and Caprifig accessions carrying the d-type haplotype) supports a selective sweep: the window overlapping the concordant interval fell in the bottom 0.90th percentile of the genome-wide distribution (Z = −2.31; Figure 4B–D). This signature was locus-specific rather than a general feature of fig domestication: Tajima’s D at 202 candidate windows for other fruit-quality loci was statistically indistinguishable from the genome-wide background (Mann–Whitney U test, P = 0.49), with only 1.0% reaching as low a percentile as *Eden*—the proportion expected by chance alone. Parthenocarpy and fruit-quality diversification were therefore subject to distinct, independent selective processes.

Genotypes recovered from wild *Ficus* resequencing data^18,19,42^ confirm the evolutionary polarity of this haplotype. At four of the eight concordant positions with reliable outgroup calls, non-Smyrna *F. carica* (homozygous d-type) shared this state with no wild species, whereas fourteen wild *Ficus* species outside subsect. *Ficus* consistently carried the e-type allele (Supplementary Table 3). The d-type haplotype is therefore derived and the e-type ancestral, consistent with the archaeobotanical dating of parthenocarpic fig cultivation to ∼11,400 years B.P. in the Jordan Valley^1^. *F. palmata*, though homozygous for the d-type haplotype at the eight diagnostic SNPs, is *F. carica*’s sister species within subsect. *Ficus*^43^ and therefore cannot serve as an independent outgroup; polarity rests on the consensus outside subsect. *Ficus*. Across the broader 25 kb *Eden* interval, this *F. palmata* accession is heterozygous at approximately one quarter of variant sites outside the diagnostic positions, matching the Smyrna-type rather than the d-type pattern, while genome-wide *D*-statistics showed no significant excess allele sharing with d-type *F. carica* (D = −0.017, Z = −1.49). These patterns argue against recent genome-wide introgression and are most consistent with long-term maintenance of ancestral (trans-species) polymorphism at *Eden* predating the *F. carica*–*F. palmata* split. [9b] Because *F. palmata* is itself a species complex spanning arid to subtropical humid climates⁴⁵ and has a long history of cultivation and deliberate hybridisation with *F. carica*, resolving whether the *Eden* configuration in this accession reflects ancestral polymorphism or introgression will require denser sampling across the complex. Smyrna-type accessions—the only cultivated class retaining a heterozygous e-type copy—thus represent the sole surviving bridge between the ancestral pollination-dependent state and the derived parthenocarpic state now dominant in cultivated *F. carica*. Fine-mapping using the *F. carica* × *F. erecta* backcross population (FcB1F1; n = 121, including 43 recombinants within the 8-SNP interval) will further resolve the causal variant.

The parthenocarpic figs recovered from Gilgal I have been interpreted as evidence of deliberate propagation predating cereal domestication^1^, an interpretation contested on the grounds that parthenocarpic trees also set seeded fruit and can reproduce spontaneously^44^. Our results cannot date the onset of selection or resolve this archaeobotanical question, but they specify what the trait requires. Fruit set in the absence of pollination depends not on the presence of the *Eden*-d haplotype but on homozygosity for it, since a single *Eden*-e copy confers pollination dependence. Homozygosity is fixed across all cultivated classes examined here and coincides with a locus-specific sweep signature, whereas Smyrna-type accessions remain heterozygous. Spontaneous reproduction alone would be expected to maintain segregation rather than fixation at this locus, a consideration that keeps the deliberate-propagation hypothesis open.

Several limitations should be noted. The cross-sectional accession panel does not provide direct evidence for the temporal sequence of trait evolution or for causality between parthenocarpy and quality diversification, and it does not include spontaneous *F. carica* populations^45^, leaving the frequency of *Eden*-d outside cultivated germplasm untested. The *MYB101-like* concordant haplotype also remains functionally unvalidated: although the concordant SNPs are predominantly non-coding, consistent with a cis-regulatory mechanism, the causal variant has not been experimentally confirmed. Reanalysis of public hormone-treatment RNA-seq data from San Pedro-type fig^24,46,47^ showed a directionally consistent increase in *MYB101-like* expression under cytokinin across all six tissue × timepoint comparisons—matching validated cytokinin-responsive controls—but no coherent response to GA₃ (Supplementary Data 4; Supplementary Fig. 10), providing exploratory support for a role in cytokinin-mediated reproductive development, though the single-library design precludes statistical testing.

Notwithstanding these limitations, this study establishes a multi-omics framework for characterising fruit flavour as an integrated sensory phenotype and provides the most comprehensive genomic dissection of fruit quality in fig to date. It identifies transporter and signalling network variation on chromosomes 01b and 05, including lipid metabolism pathway components, as priority targets for flavour-oriented breeding, and shows that reproductive innovation and sensory diversification are genetically separable dimensions of fig domestication. One plausible hypothesis arising from this separation is that reproductive innovation preceded fruit quality diversification, consistent with the classical distinction between early-fixed domestication traits and later-selected improvement traits^9^; testing it will require temporal evidence beyond the cross-sectional design used here. More broadly, the framework provides a template for analogous analyses in other perennial fruit crops.

## Methods

### Plant material and growth conditions

A total of 483 accessions are catalogued in Supplementary Data 1, of which 482 were subjected to whole-genome resequencing; one *F. carica* analytical reference sample was catalogued but not resequenced. The resequenced panel comprised 217 newly sequenced *F. carica* accessions (165 cultivars and parental lines and 52 F1 progeny from crosses within *F. carica*; by horticultural class, 118 Common-type, 38 Caprifig, 4 San Pedro-type, 4 Smyrna-type and 53 of unassigned class), 236 *F. carica* × *F. erecta* interspecific hybrids (FcB1F1, n = 121; FcB2F1, n = 114; and the F1 hybrid FEBN7), 1 *F. erecta*, 1 *F. palmata*, and 27 previously published *Ficus* genomes^18,19^.

For the parthenocarpy concordance scan, accessions were classified by fruit habit into non-Smyrna accessions capable of parthenocarpic fruit set (cases) and Smyrna-type accessions requiring caprification (controls); the composition of each group, and the accessions excluded from the scan, are detailed under the concordance scan below (Supplementary Data 1).

For fruit quality and multi-omics profiling, morphological traits were measured for 81 accessions; the accession sets used for each omics analysis are given in the relevant subsections below. Fruits were harvested at the ripe stage in 2020 for sugar – acid composition, volatile aroma, taste sensor, and transcriptome analyses; sampling seasons for colorimetric and anthocyanin traits are given below. Morphological traits—fruit weight (g), longitudinal and horizontal diameter (cm), fruit colouring ratio (%), peduncle length (cm), and fruit hardness (N)—were measured by standard methods in 2017–2018.

### Whole-genome resequencing and variant calling

Whole-genome resequencing data and variant datasets (VCF files) used in this study are identical to those reported in Ikegami et al.^42^, to which readers are referred for full details of sequencing, mapping, and variant calling. Briefly, 482 accessions were resequenced at a mean depth of 15.23×, and variants were called against the female-derived reference assembly FCAr2.3 and the male-derived draft assembly FCA6085_r1.1. After quality filtering, 3,275,911 high-confidence SNPs were retained, comprising 2,044,654 transitions and 1,231,257 transversions (Ts/Tv = 1.66; BCFtools stats). Functional impact of SNPs was predicted using SnpEff^48^. Chromosome numbering follows the Horaishi-based assemblies^41,42^; correspondence to the Dottato assembly^49,50^ is given in Supplementary Table 4.

### Population structure analysis

To characterise genetic relationships among accessions, principal component analysis (PCA), neighbour-joining tree analysis, and ADMIXTURE analysis were performed on a linkage-disequilibrium-pruned subset of the quality-filtered biallelic SNP dataset (3,275,911 SNPs). After conversion of non-standard chromosome names to numeric codes and assignment of unique SNP identifiers, LD pruning in PLINK v1.9 (--indep-pairwise 50 10 0.2) retained 164,933 SNPs. PCA was performed in PLINK on this dataset. The neighbour-joining tree was constructed from a pairwise 1−IBS genetic distance matrix computed in PLINK from the same SNP set, with topology inferred using the nj() function in the R package ape. ADMIXTURE v1.3.0 was run for K = 2 to K = 10 with five-fold cross-validation. Population stratification in GWAS was assessed by calculating the genomic inflation factor (λGC).

#### Transcriptome profiling

Total RNA was extracted from syconium tissues at the second developmental stage using the Fruit-mate for RNA Purification reagent (Takara Bio, Shiga, Japan) following the protocol optimised for *Ficus* tissues^51^. RNA quality was assessed on an Agilent Bioanalyzer, and libraries were constructed with the QuantSeq 3′ mRNA-Seq kit (Lexogen, Vienna, Austria). A total of 102 transcriptome libraries were constructed and sequenced on an Illumina NextSeq 500. Clean reads were mapped to the FCAr2.3 reference genome using HISAT2^52^, and transcript abundance was quantified as expected counts using RSEM^53^, which assigns multi-mapping reads probabilistically via expectation–maximisation. Expression levels were quantified for 45,900 genes comprising 45,608 nuclear, 117 chloroplast, and 175 mitochondrial genes. Genes with no detectable expression (read count = 0) across the dataset were excluded, retaining 30,081 expressed genes. Expected counts for the retained genes were normalized using the trimmed mean of M-values (TMM) method as implemented in edgeR^54^, followed by averaging across technical/biological replicates prior to downstream multi-omics integration. The resulting normalized expression matrix is provided in Supplementary Data 5; full gene-level functional annotation for all 45,900 gene models is provided in Supplementary Data 3.

#### Sugar and acid compounds profiling

To profile sugar and organic acid components of fig fruit (syconia), 102 accessions were analysed by GC–MS. Fruits harvested at the ripe stage in 2020 were cut into wedge-shaped pieces, weighed, and homogenised with four volumes of methanol using 5 mm-diameter zirconia beads. After centrifugation, the supernatant was passed through a Monospin C18 column (GL Sciences, Japan); the unbound fraction was collected, dried under a nitrogen stream, and derivatised by methoxymation followed by trimethylsilylation. Analysis used a GCMS-TQ8050NX system with an AOC-30i autoinjector (Shimadzu, Japan), helium carrier gas at a linear velocity of 39 cm/s, splitless injection (0.5 μL, inlet 250°C), and an oven programme of 80°C for 2 min, ramped at 12°C/min to 320°C and held for 10 min. Separation used an Rxi-5Sil MS column with Integra-Guard (30 m × 0.25 mm ID, 0.5 μm film thickness; RESTEK). Compounds were identified by matching retention time and ion transitions against an in-house MRM library constructed from authentic standards, and 41 sugar and organic acid parameters were quantified.

#### Aroma compounds profiling

Aroma compounds were profiled in 121 accessions by headspace sampling with MonoTrap RCC18 (GL Sciences, Japan) coupled to GC–MS, yielding 549 volatile features for downstream analysis. Fruits were cut into wedge-shaped pieces, weighed, and homogenised with an equal volume of ultrapure water. A 5 mL aliquot of the homogenate was combined with 2 g sodium chloride in a 20 mL headspace vial, and a MonoTrap RCC18 disc was positioned in the headspace to collect volatiles at 60°C for 16 h. The disc was then immersed in 120 μL dichloromethane to elute the captured volatiles, and 1 μL of the eluate was injected. Analysis used a GCMS-TQ8040 system with an AOC-20i autoinjector (Shimadzu, Japan), helium carrier gas at a linear velocity of 30 cm/s, splitless mode, inlet temperature 250°C, and an oven programme from 50°C ramped at 3°C/min to 200°C followed by a bake-out step at 260°C for 8 min. Separation used a SUPELCOWAX10 column (30 m × 0.25 mm ID, 0.25 μm film thickness; SUPELCO). Volatiles were tentatively identified using the FFNSC3 mass spectral library, with retention indices calculated against both a C7–C33 n-alkane series^55^ and a C4–C24 fatty acid ethyl ester (FAEE) series. Analyses were conducted during the 2020 growing season.

#### Anthocyanin and colorimetric profiling

Fruit peel colour was characterised by visual classification according to the Munsell colour system and by colorimetric measurement with a Minolta CR-300 chromameter recording CIE L*a*b* parameters; values represent the mean of measurements taken in 2018 and 2019. Peel samples for anthocyanin analysis were collected from 82 accessions at the ripe stage in 2020 and 2021. Anthocyanins were analysed according to the method of Ishibashi et al.^56^: peel tissue (approximately 100 mg) was homogenised in 1 mL of 50% (v/v) ethanol containing 0.1% (v/v) formic acid, centrifuged at 15,000 × g for 5 min at 4°C, and the extract diluted 10-fold in the same solvent before analysis on a PESI-MS/MS system (LCMS-8045 tandem mass spectrometer with a DPiMS-8060 probe electrospray ionisation source; Shimadzu, Kyoto, Japan). Data acquisition and analysis used PESI MS Solution v2.0.8 and LabSolutions v5.86 (Shimadzu). Anthocyanin contents were quantified from extracted ion intensities and normalised to sample fresh weight. Six anthocyanin species were quantified: pelargonidin-3-glucoside (P3G), pelargonidin-3-rutinoside (P3R), cyanidin-3-glucoside (C3G), cyanidin-3-rutinoside (C3R), cyanidin-3-dioxalylglucoside (C3dG) and cyanidin-3-malonylglucoside (C3M).

#### Taste sensor measurements

Taste attributes were measured in 96 accessions using an electronic taste sensor (TS-5000Z, Intelligent Sensor Technology, Kanagawa, Japan) equipped with five sensor electrodes (CA, CT0, C00, AE and AP). Fruit tissue was homogenised with 4 volumes of distilled water and 50 mL of the resulting extract used per measurement. Eight taste dimensions—sourness, saltiness, bitterness, umami, astringency, bitter off-flavor, astringent stimulation and rich umami—were recorded as sensor output values, in triplicate for each accession.

#### Multivariate analysis of phenotypic traits

Hierarchical clustering and principal component analysis (PCA) were performed independently on each omics dataset (sugar–acid composition, volatile aroma profiles, colorimetric and anthocyanin parameters, and taste sensor measurements) in R v4.6.0. For clustering, a dissimilarity matrix was computed as d = 1 − r (r, Pearson correlation coefficient) and agglomerative clustering performed with the Ward.D2 method (hclust). PCA used scaled and centred data (prcomp). Heatmaps were generated with the ComplexHeatmap package^57^.

#### Genotype–phenotype concordance scan for parthenocarpy

To identify genomic regions associated with the capacity for parthenocarpic fruit set, we performed a genome-wide genotype–phenotype concordance scan contrasting non-Smyrna accessions, which set fruit without pollination, against Smyrna-type accessions, which abscise their syconia in the absence of caprification, guided by the classical +P (persistence) hypothesis for persistent versus caducous syconia^2,6,58^. The scan compared 118 non-Smyrna cases (114 Common-type and 4 San Pedro-type, comprising 101 *F. carica* and 17 *F. carica* F1) with 4 Smyrna-type controls (Fc_25, Fc_57, Fc_58, Fc_60). The remaining 360 resequenced accessions were not scored for fruit habit and were excluded, comprising male caprifigs, wild *Ficus* relatives, interspecific hybrid populations, unphenotyped lines, potential genotypic duplicates, and accessions of uncertain horticultural class (Supplementary Data 1). Because the control group was small and highly imbalanced relative to the cases, conventional association testing was considered underpowered; the concordance scan was therefore used as a candidate-locus discovery approach to identify regions showing complete allelic separation between the two groups. To evaluate selective sweep signatures at the chr04 concordant locus, Tajima’s D was calculated in non-overlapping 10 kb windows across the genome using VCFtools v0.1.16^59^, restricted to Common-type and Caprifig accessions carrying the d-type haplotype at the 8 concordant SNPs; windows with fewer than one segregating site were excluded, yielding 28,202 valid windows genome-wide (3,196 on chr04). Percentile ranks were calculated with scipy.stats.percentileofscore (Python 3). For comparison, GWAS/TWAS candidate loci for fruit quality traits (Supplementary Tables 1, 2; Supplementary Data 2) were deduplicated to unique genomic positions, mapped to their corresponding 10 kb windows in the same population-restricted dataset, and their Tajima’s D distribution compared with the genome-wide background using a two-sided Mann–Whitney U test (scipy.stats.mannwhitneyu).

The analysis was performed using SNP datasets mapped independently to the female-derived reference assembly FCAr2.3 and the male-derived draft assembly FCA6085_r1.1. Fruit habit was treated as a binary trait based on the clear qualitative distinction between Smyrna-type and non-Smyrna accessions under standard field conditions. Conventional association results were not used as the sole basis for candidate-locus identification because the small and imbalanced control set limited statistical power and robustness, particularly with respect to population structure.

For each SNP, genotype–phenotype concordance was evaluated under alternative dominant and recessive genotype models. Accessions were classified into genotype classes and compared with the binary fruit-habit phenotype, and the highest match rate between genotype class and phenotype class was retained as the concordance score; SNPs with a score of 1.0 were defined as perfectly concordant. Under the dominant model, dosage 0 (homozygous for the non-counted allele) was assigned to the case (non-Smyrna) class, and dosage 1 (heterozygous) or dosage 2 (homozygous for the counted allele) to the control (Smyrna) class. Concordance analyses were first performed on the imputed SNP datasets for each reference assembly to avoid artefactual complete matches caused by missing genotypes; perfectly concordant SNPs were then checked in the corresponding non-imputed datasets to confirm that they were present as observed rather than imputation-only variants.

To assess the statistical specificity of the chromosome 04 concordant haplotype, case–control labels were randomly permuted 10,000 times while maintaining the observed case–control ratio, and the number of perfectly concordant SNPs on chromosome 04 and across the genome was recorded for each permutation. Empirical P values were calculated as (k + 1)/(10,000 + 1), where k is the number of permutations yielding perfectly concordant SNP counts equal to or greater than the observed counts.

Candidate gene annotation within the concordant interval used the FCAr2.3 gene models, focusing on the *MYB101-like*-centred region and the adjacent 2-alkenal reductase gene. Read-based inspection of the *MYB101-like* region, including the upstream interval containing perfectly concordant SNPs, was performed by mapping reads from a representative Smyrna-type control (Fc_57, Sari Lop) and a representative non-Smyrna parthenocarpic accession to the FCAr2.3 reference assembly and visualising read alignment, local variants, and read depth in IGV. To infer the evolutionary polarity of alleles at the concordant SNPs, publicly available whole-genome resequencing reads from wild *Ficus* species^18,19^ (BioProject PRJNA684963; SRA run accessions SRR13279546–SRR13279552 and SRR13279570–SRR13279575; species listed in Supplementary Table 3) were mapped to FCAr2.3 and genotypes at the eight concordant positions called and compared with *F. carica*. Alleles shared between non-Smyrna *F. carica* and wild outgroup species were inferred as ancestral, and *F. carica*-specific alleles as derived (Supplementary Table 3).

#### Bayesian GWAS and TWAS for fruit quality traits

Genome-wide and transcriptome-wide association analyses were performed for all fruit quality traits using a Bayesian framework. We assumed that the phenotype of a fruit quality trait for the *i*th individual, yi, is expressed as

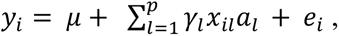

where *m* is a mean of the trait, *xil* is a covariate indicating the genotypes of the *l*th SNP of the *i*th individual in GWAS, taking values 1, 0 and -1 for a homozygous with a reference allele, heterozygous and a homozygous with alternative allele, or indicating standardized expression level of the *l*th gene of the *i*th individual in TWAS, respectively (*l*=1,2,…,*p*), *al* is an effect of the covariate *xil* on the phenotype, *gl* is an indicator variables taking values 1 and 0 corresponding to whether the SNP or gene is included or excluded in the model and *ei* is a residual of the model. Using the variational approximation method of Bayesian estimation, posterior means were approximately estimated for the model parameters following Hayashi and Iwata^60^. For the determination of whether each SNP or gene affects the phenotype, we evaluated posterior mean of *gl*, which are regarded as the approximate posterior probability of the *l*th SNP or gene being included in the model as a significant factor relevant to the phenotype (*l*=1,2,…,*p*).

Whole-genome variants were filtered by removing non-diploid calls, alleles with minor allele frequency < 0.05, and variants with a missing rate > 10%. For GWAS, each trait was analysed using a Bayesian sparse linear mixed model (BSLMM) implemented in GEMMA^61,62^, in which SNP effects were assumed to follow a mixture of a normal distribution and a point mass at zero, allowing simultaneous estimation of polygenic background and large-effect loci; population structure was accounted for by including a genomic relatedness matrix (GRM) computed from the filtered SNP data. For TWAS, the association between gene expression (TPM) and each trait was evaluated using a Bayesian linear mixed model with the GRM as a random effect. Posterior probabilities (PP) > 0.5 were used as the threshold for reporting candidate loci, and GWAS and TWAS results were integrated by identifying candidate pairs located within ±100 kb of each other on the same chromosome.

#### Multi-omics factor analysis

MOFA+ was applied to characterise the integrated architecture of fig fruit flavour, defined here as the multimodal sensory experience encompassing aroma, taste, and primary metabolite composition. Colorimetric and anthocyanin traits, which represent visual rather than direct flavour attributes, were excluded from the integration and analysed separately by GWAS and TWAS. A multi-layer dataset of 81 accessions with complete data across four omics layers—volatile aroma profiles (549 features; n = 121 in the full dataset), sugar–acid composition (41 parameters; n = 102), taste sensor measurements (8 dimensions; n = 96), and RNA expression (45,900 genes; n = 102)—was analysed using Multi-Omics Factor Analysis (MOFA+; MOFA2 R package v1.22.0 with the mofapy2 backend v0.7.4)^17^. Gaussian likelihoods were applied to all data layers. The model was trained with 15 latent factors using a single random initialisation (seed 42) and converged after 286 iterations; no post hoc factor-discard threshold based on variance explained was applied. Inter-layer connectivity networks were constructed in R using igraph and visualised with ggraph. Cross-modal modulation by individual volatile features was assessed using Lasso regression and ordinary least squares (OLS) regression, with taste sensor outputs as response variables and aroma features as predictors.

In addition, correlation network analysis was performed using ConfeitoGUI^25^ across the same four omics layers used in the MOFA+ analysis. Pairwise correlations were computed using three metrics (Pearson, Spearman, and cosine similarity), and community detection was applied to identify co-accumulation and co-expression modules across layers. Results from the Pearson-based network are presented as representative (Supplementary Fig. 9).

#### Gene functional annotation

Protein-coding genes in the FCAr2.3 reference assembly were predicted using the MAKER pipeline^63^ with Augustus and SNAP as described in Ikegami et al.^42^. Predicted protein sequences were used as queries for BLASTp-based similarity searches against the NCBI non-redundant protein (nr) database (E-value < 1×10⁻⁴), with additional pathway and orthology assignments from KEGG. Full gene-level annotation details for all 45,900 genes are provided in Supplementary Data 3.

## Data availability

The whole-genome resequencing data and SNP variant datasets (VCF files) analysed in this study were generated in Ikegami et al.^42^ and are available from the DDBJ Sequence Read Archive under accession numbers DRA016899, DRA016900 and DRA016910. The chromosome-scale reference genome assemblies used for read mapping and gene annotation—FCAr2.3 (female-derived Horaishi; DDBJ accession numbers AP028881–AP028893) and FCA6085_r1.1 (male-derived Caprifig6085; DDBJ accession numbers AP028894–AP028906)—are also described in Ikegami et al.^42^ and are accessible through the Plant Garden Database (*F. carica*: https://plantgarden.jp/en/list/t3494; *F. erecta*: https://plantgarden.jp/en/list/t66383). The RNA-seq reads generated in this study have been deposited in the DDBJ Sequence Read Archive under BioProject accession PRJDB42532.

Publicly available whole-genome resequencing data for wild *Ficus* species were obtained from NCBI BioProject PRJNA684963 (SRA run accessions SRR13279546–SRR13279552 and SRR13279570–SRR13279575). Publicly available hormone-treatment RNA-seq data for San Pedro-type fig were obtained from NCBI SRA accession SRP113799 (gibberellin treatment)^24^ and BioProject PRJNA397979 (cytokinin treatment)^47^. The primary metabolite, volatile, colorimetric, anthocyanin, taste sensor and normalised expression datasets underlying the analyses are provided in Supplementary Data 1–5. All other data supporting the findings of this study are available within the article and its Supplementary Information.

## Code availability

All analyses were performed using publicly available software: PLINK v1.9, ADMIXTURE v1.3.0, HISAT2, RSEM, edgeR, SnpEff, BCFtools, VCFtools v0.1.16, GEMMA, MAKER, IQ-TREE with ModelFinder, IGV and ConfeitoGUI, together with the R packages ape, ComplexHeatmap, MOFA2 v1.22.0 (mofapy2 backend v0.7.4), igraph and ggraph in R v4.6.0, and SciPy in Python 3. Custom scripts used for the genome-wide genotype–phenotype concordance scan, the phenotype-label permutation test, the Tajima’s D percentile and Mann–Whitney U comparisons, the MOFA+ workflow, and the Lasso and ordinary least squares cross-modal regression analyses are available from the corresponding author upon reasonable request.

## Supporting information

Supplementary Data 1: Accession composition and multi-omics analysis panel

Supplementary Data 2: Full list of GWAS and TWAS candidate loci for volatile aroma traits

Supplementary Data 3: Functional annotation of Ficus carica (FCAr2.3) gene models

Supplementary Data 4: Expression of MYB101-like and control genes in public RNA-seq data

Supplementary Data 5: TMM-normalized gene expression matrix

## Acknowledgements

This study was supported by JSPS KAKENHI (26850025, 16H04878, 16H06279 [PAGS], 19H02952, 22H05181)

We thank Hideyuki Nakayama and Shunichi Nakamura (Fukuoka Agriculture and Forestry Research Center Buzen Branch) for cultivation and field management of the fig germplasm collection, Rempei Kuwahara (Hirata Corporation) for the ConfeitoGUI correlation network analysis, and Atsushi Kurabayashi (Kazusa DNA Research Institute) for GC–MS technical support.

The authors used Claude to assist with language editing and manuscript formatting during preparation of this work. The authors reviewed and edited all AI-assisted output and take full responsibility for the content of the manuscript.

---

**Supplementary Table 1.**
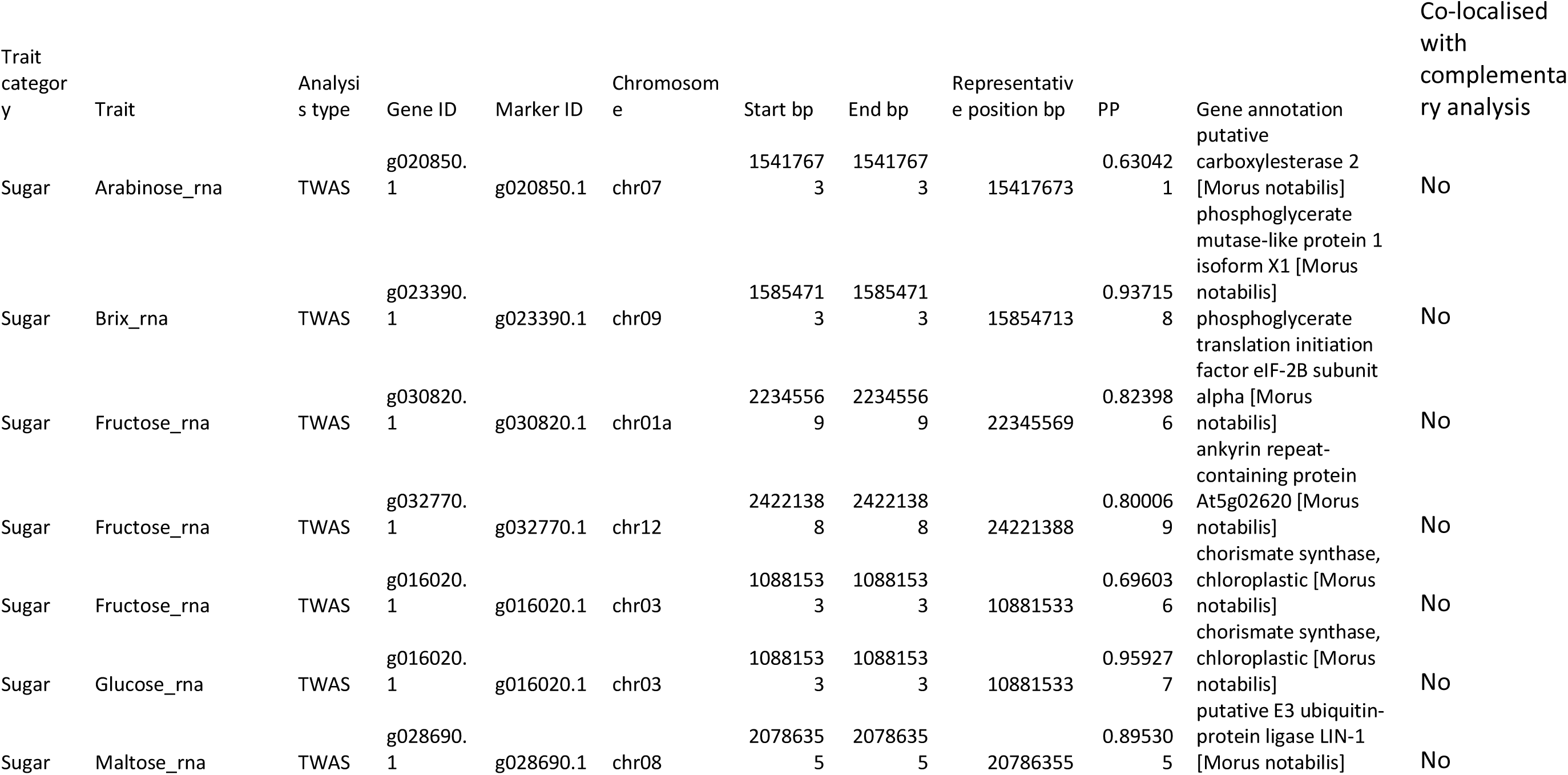
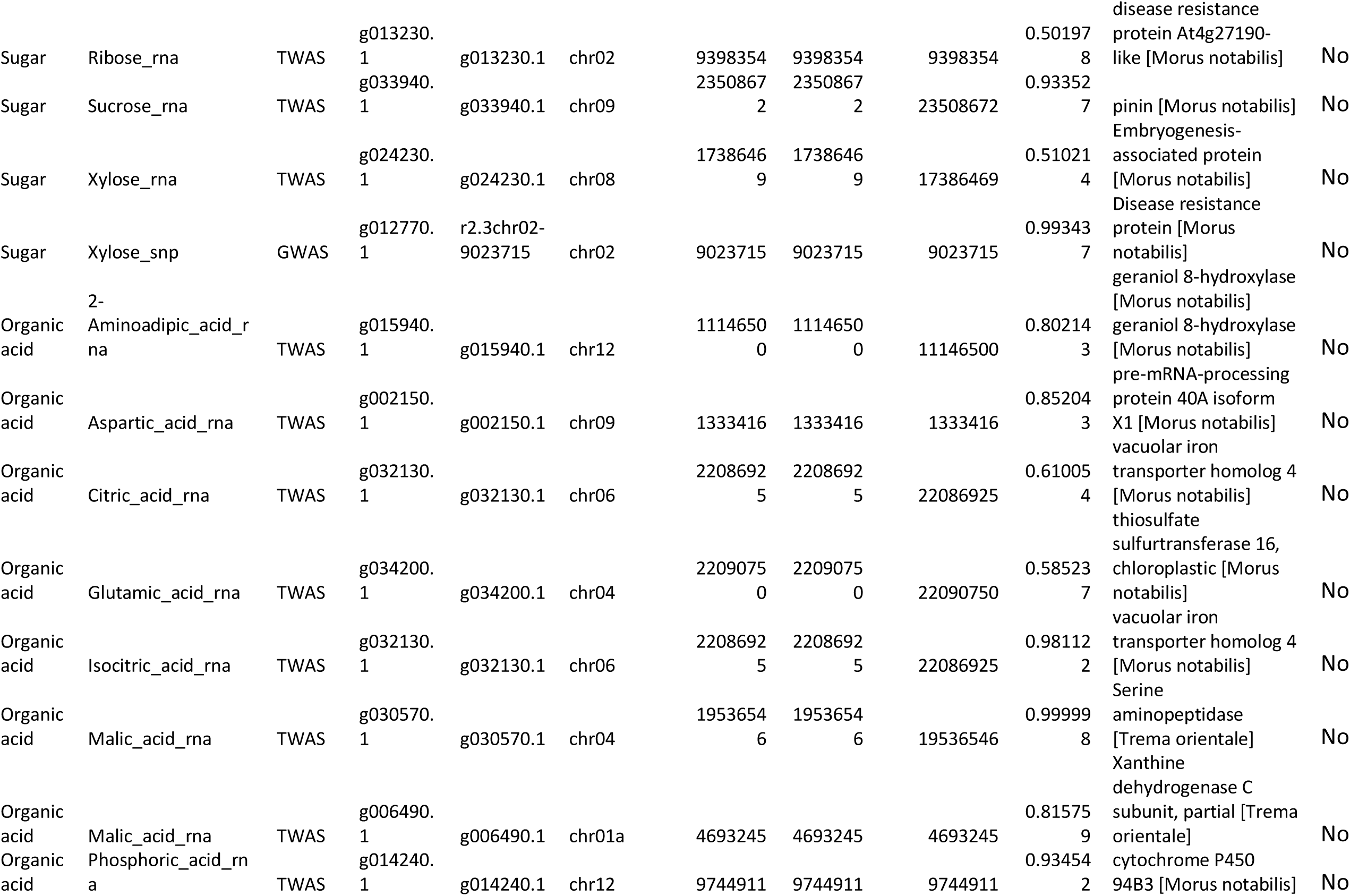
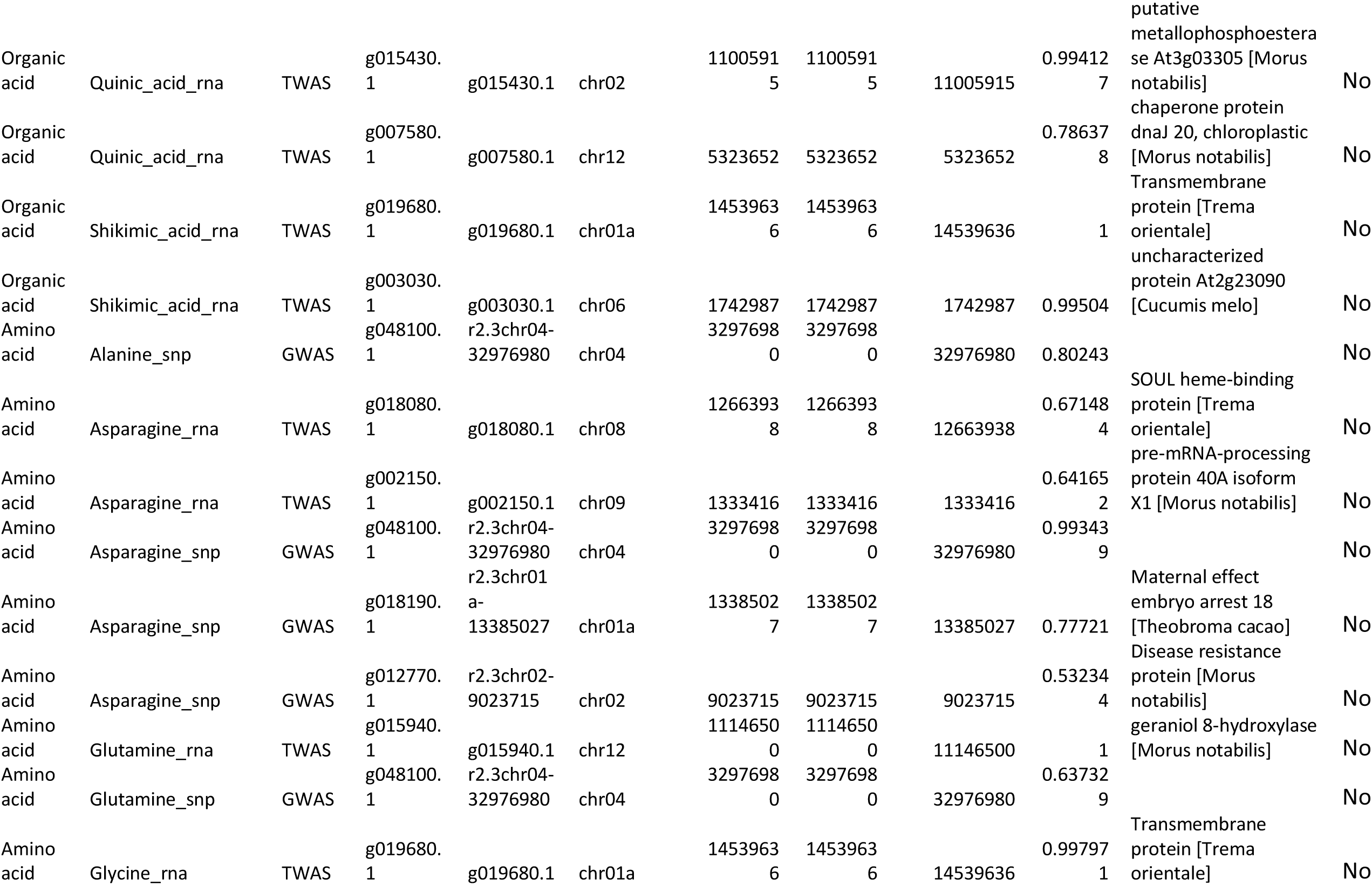
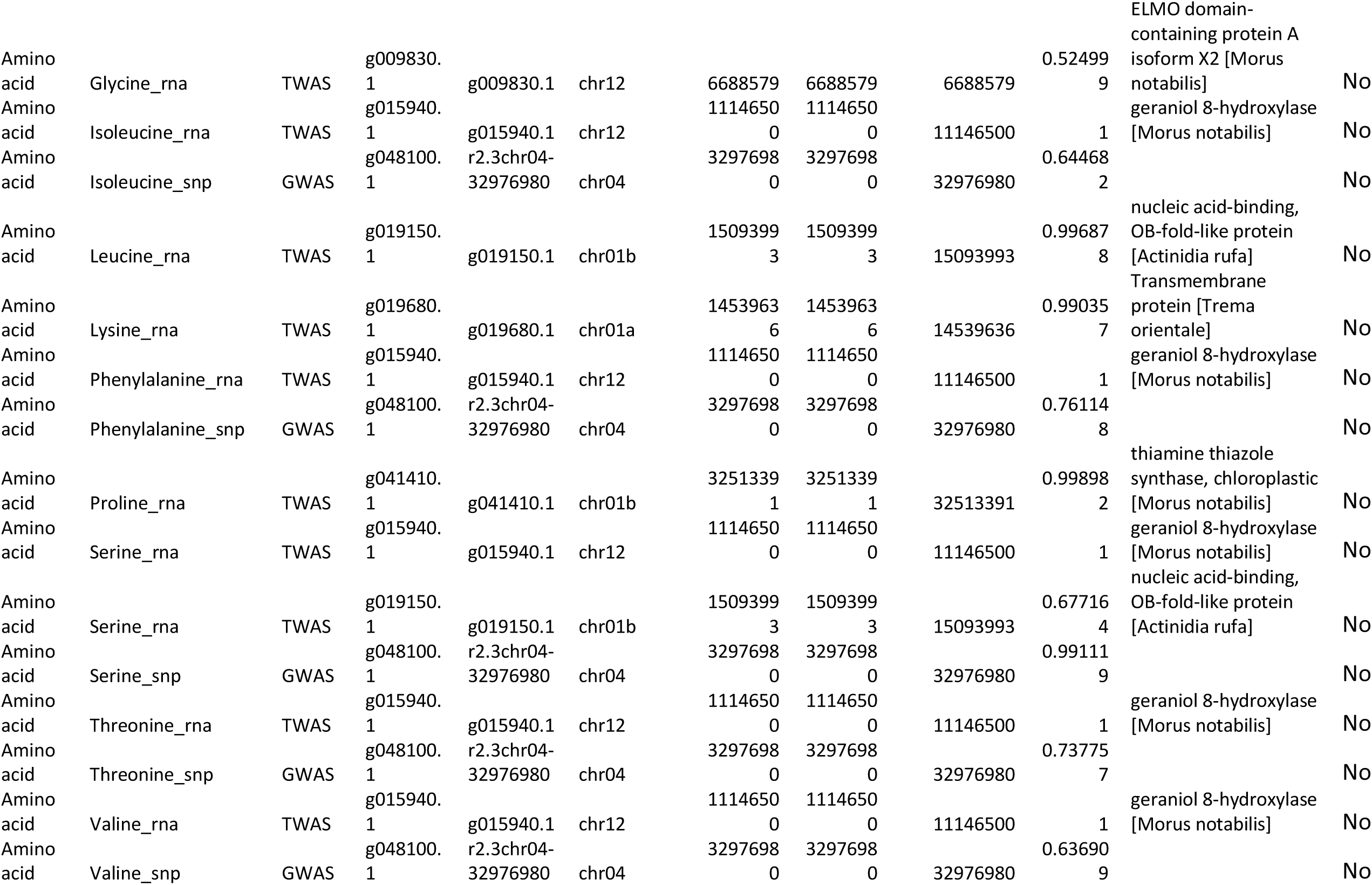
Full list of GWAS and TWAS candidate loci for primary metabolite traits. Candidate loci identified by Bayesian GWAS and TWAS for primary metabolite traits with posterior probability (PP) > 0.5 are listed (n = 47 entries total: 11 GWAS and 36 TWAS candidates across sugar, organic acid, and amino acid trait categories). For each candidate, the table reports trait category, associated trait, analysis type, gene ID, marker ID, chromosomal position (start bp, end bp, and representative position bp), PP value, gene annotation, co-localisation status with candidates from the complementary analysis within ±100 kb, matched GWAS/TWAS candidate ID, distance to matched candidate (bp), and overlap status with the chromosome 04 parthenocarpy candidate interval (13.785–13.810 Mb). One TWAS candidate, g034200.1 (chr04; Glutamic acid; PP = 0.585), was located within the chromosome 04 parthenocarpy candidate interval; this locus is distinct from the amino acid GWAS candidate g048100.1 and does not affect the genetic independence of primary metabolite loci from the parthenocarpy-associated haplotype. No GWAS–TWAS co-localised pairs were identified within ±100 kb for primary metabolite traits in this dataset. Key candidates include the TWAS Brix-associated candidate g023390.1 (chr09; PP = 0.937; phosphoglycerate mutase-like protein), the TWAS multi-amino-acid candidate g015940.1 (chr12; PP = 1.000 for glutamine, isoleucine, phenylalanine, serine, threonine, and valine; PP = 0.802 for 2-aminoadipic acid; geraniol 8-hydroxylase), and the GWAS multi-amino-acid locus g048100.1 (chr04; PP > 0.5 for eight amino acids). Candidates are sorted by trait category, trait, and descending PP value.

**Supplementary Table 2.**
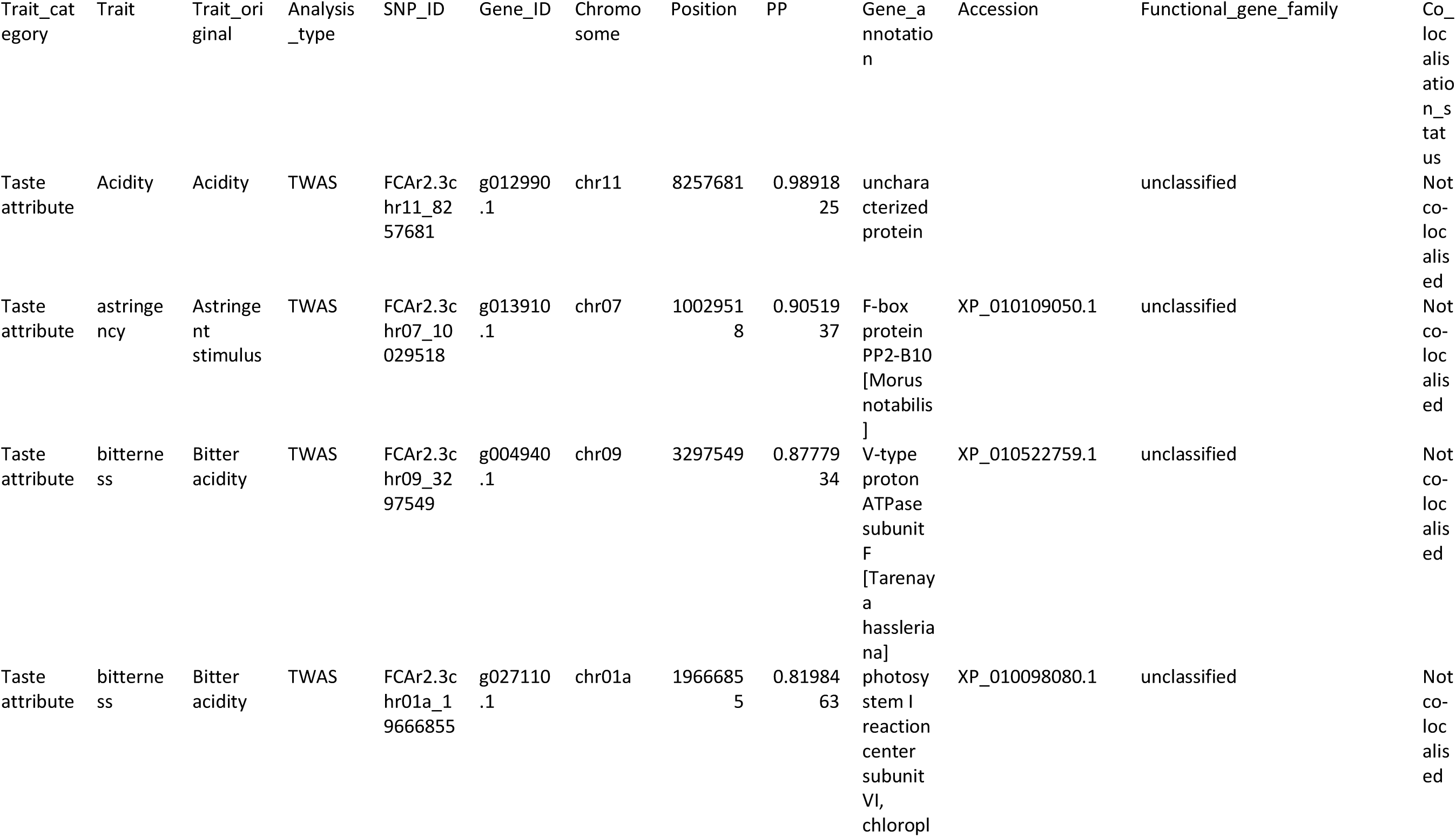
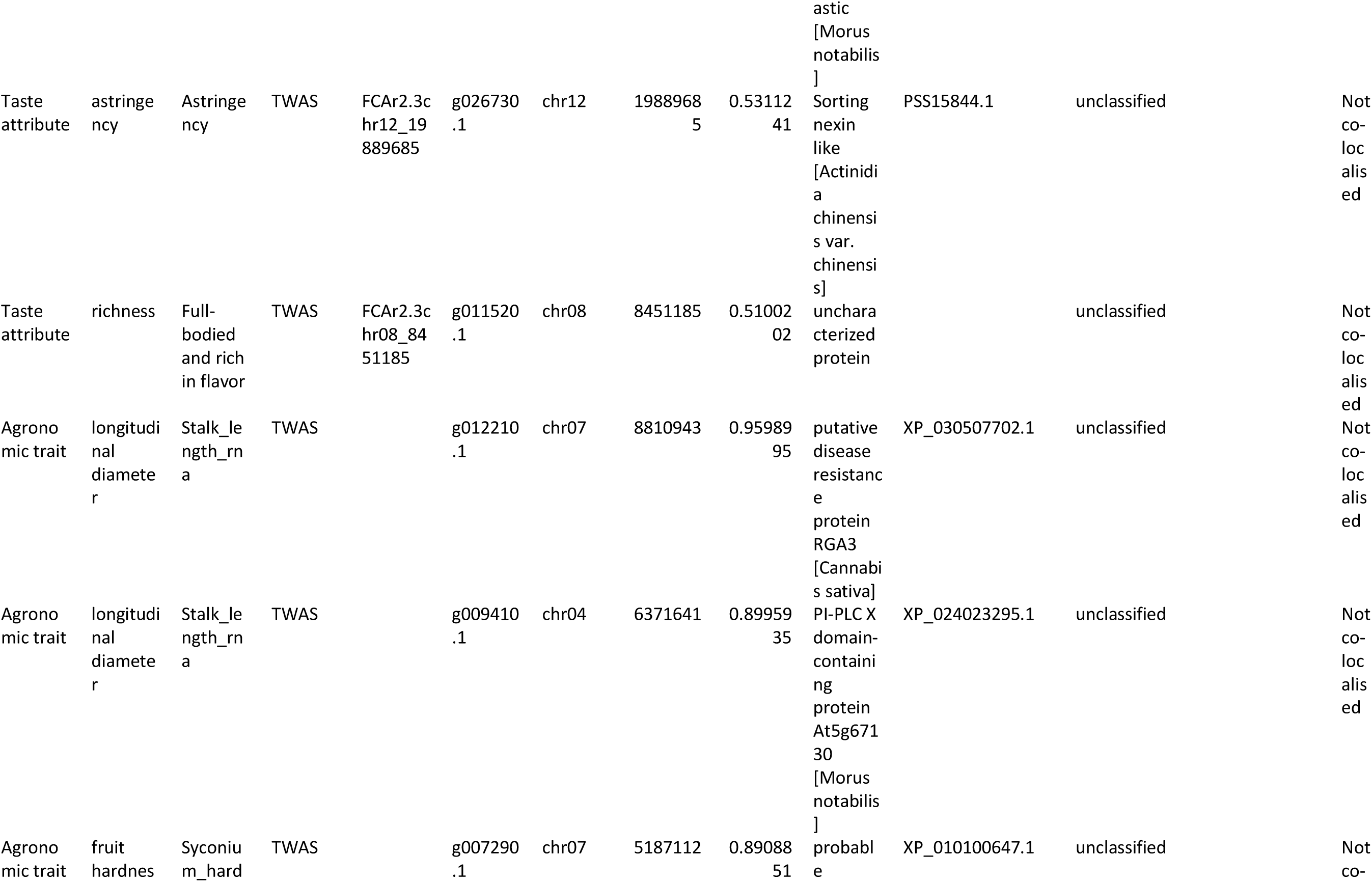
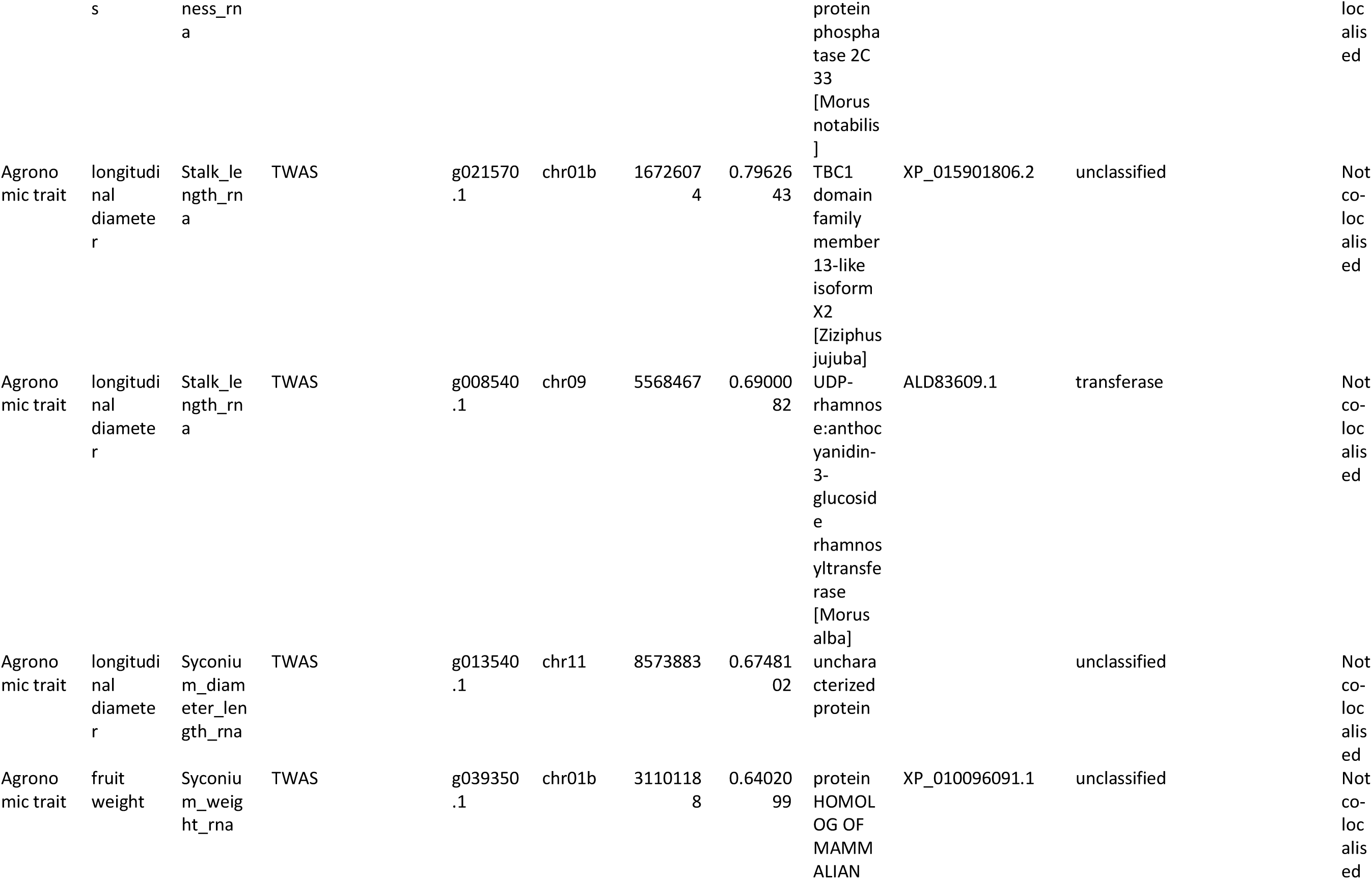
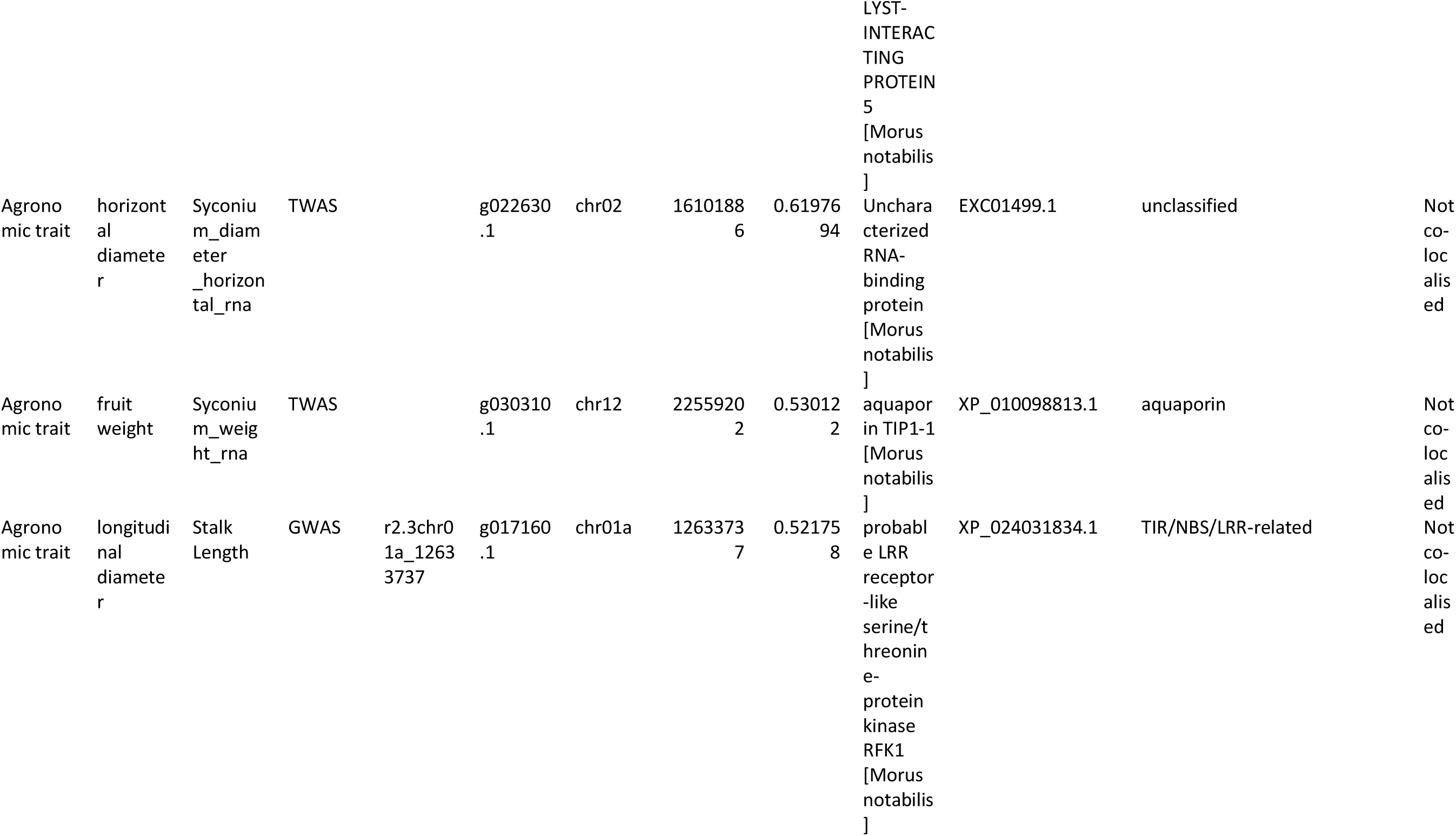
Full list of GWAS and TWAS candidate loci for taste attributes and agronomic quality traits. Candidate loci identified by Bayesian GWAS and TWAS for taste sensor attributes and agronomic quality traits with posterior probability (PP) > 0.5 are listed (n = 16 entries total: 6 taste attribute candidates and 10 agronomic trait candidates). For each candidate, the table reports trait category, associated trait, original trait label, analysis type, SNP ID or gene ID, chromosomal position, PP value, gene annotation, accession number, functional gene family classification, co-localisation status with candidates from the complementary analysis within ±100 kb, co-localised candidate ID, distance to complementary candidate (bp), hotspot flag, and source file. For taste attribute candidates (n = 6), all entries are TWAS-derived and include high-confidence candidates for acidity (g012990.1; chr11; PP = 0.99), astringent stimulus (g013910.1; chr07; PP = 0.91), bitter acidity (g004940.1; chr09; PP = 0.88; g027110.1; chr01a; PP = 0.82), astringency (g026730.1; chr12; PP = 0.53), and full-bodied and rich flavour (g011520.1; chr08; PP = 0.51). The chromosome 08 full-bodied flavour hotspot (∼8.45 Mb) is flagged in the hotspot_flag column. For agronomic quality trait candidates (n = 10), entries span fruit weight, longitudinal diameter, horizontal diameter, peduncle length, and fruit hardness. Key candidates include aquaporin TIP1-1 g030310.1 (chr12; fruit weight TWAS; PP = 0.530), putative disease resistance protein RGA3 g012210.1 (chr07; peduncle length TWAS; PP = 0.960), protein phosphatase 2C 33 g007290.1 (chr07; fruit hardness TWAS; PP = 0.891), and LRR receptor-like kinase g017160.1 (chr01a; peduncle length GWAS; PP = 0.522). Notably, UDP-rhamnose:anthocyanidin-3-glucoside rhamnosyltransferase g008540.1 (chr09; PP = 0.690)—also identified as the primary pigmentation candidate in Supplementary Data 2—appears as a TWAS candidate for longitudinal diameter, providing evidence for pleiotropic regulatory effects of this locus across secondary metabolic and morphological traits. Candidates are sorted by trait category and then by descending PP value within each category.

**Supplementary Table 3.**
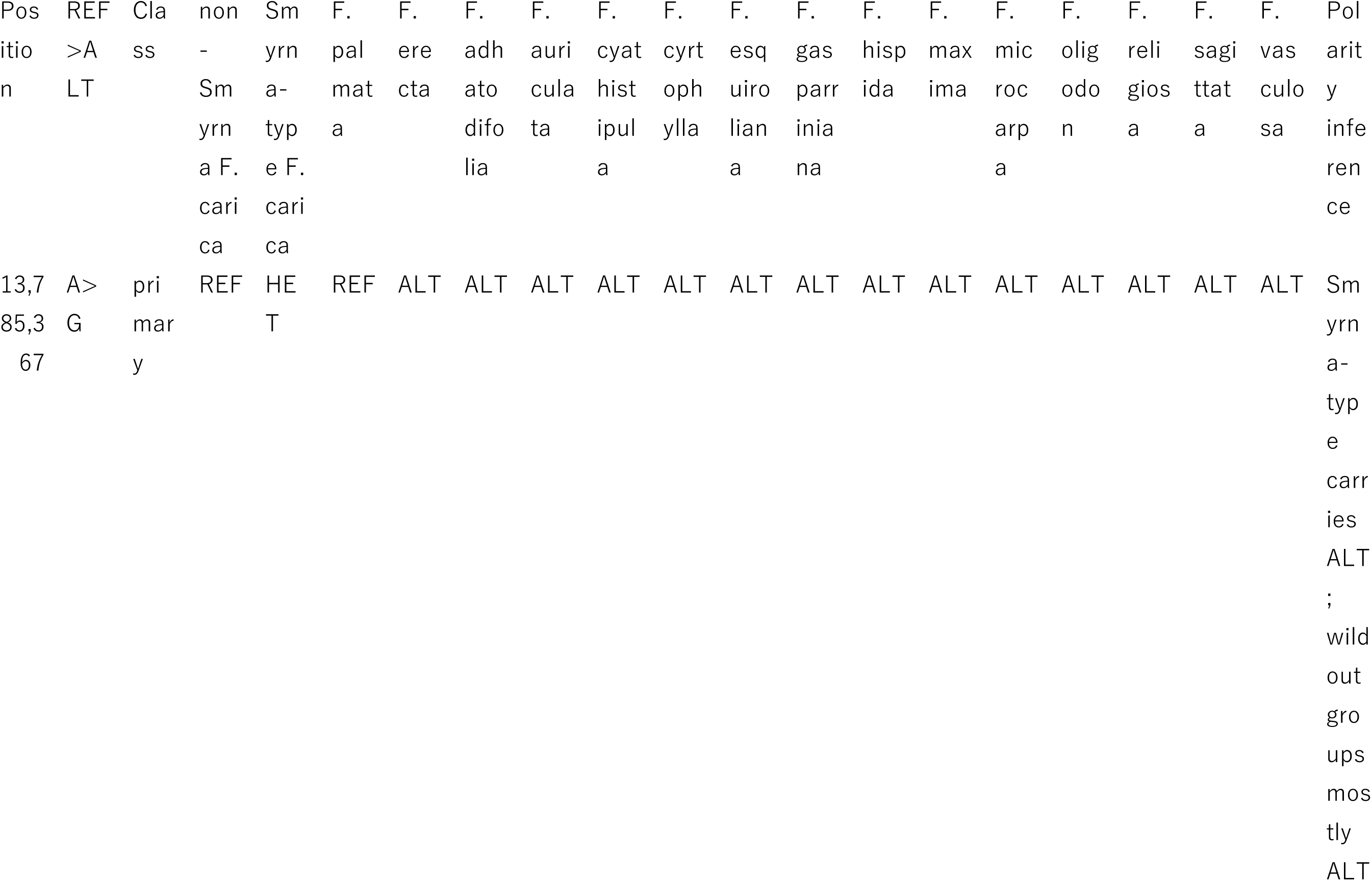
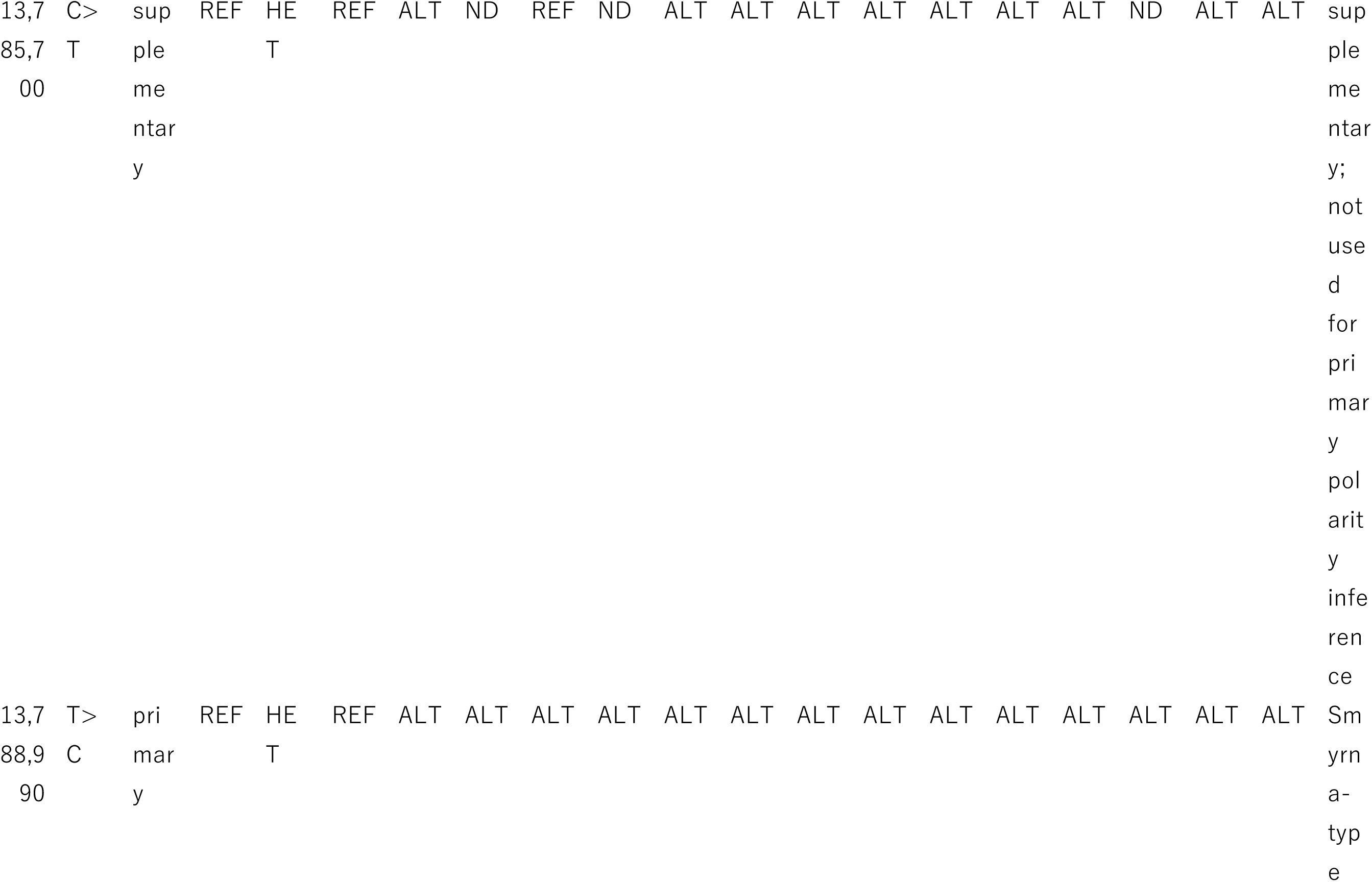
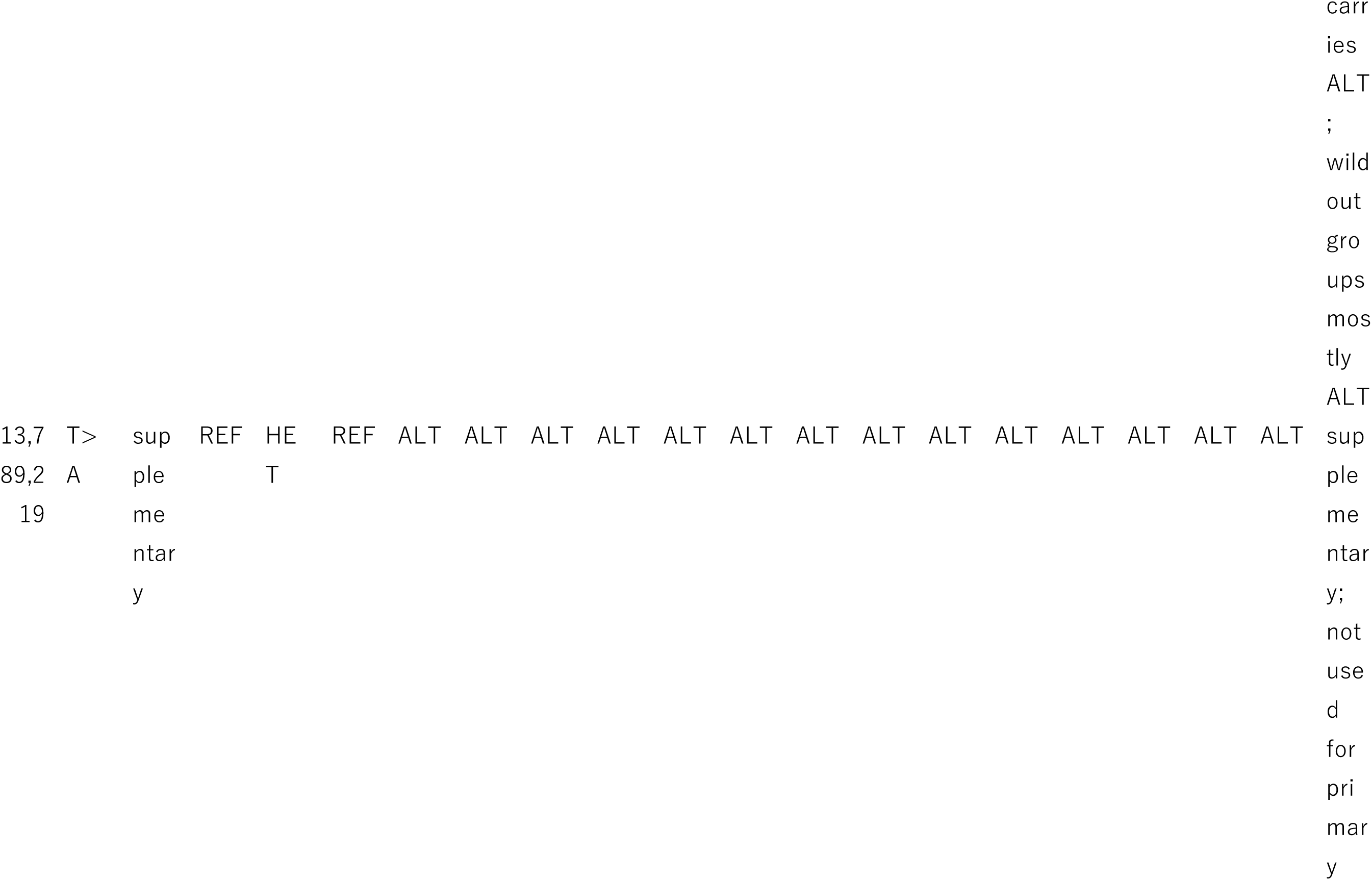
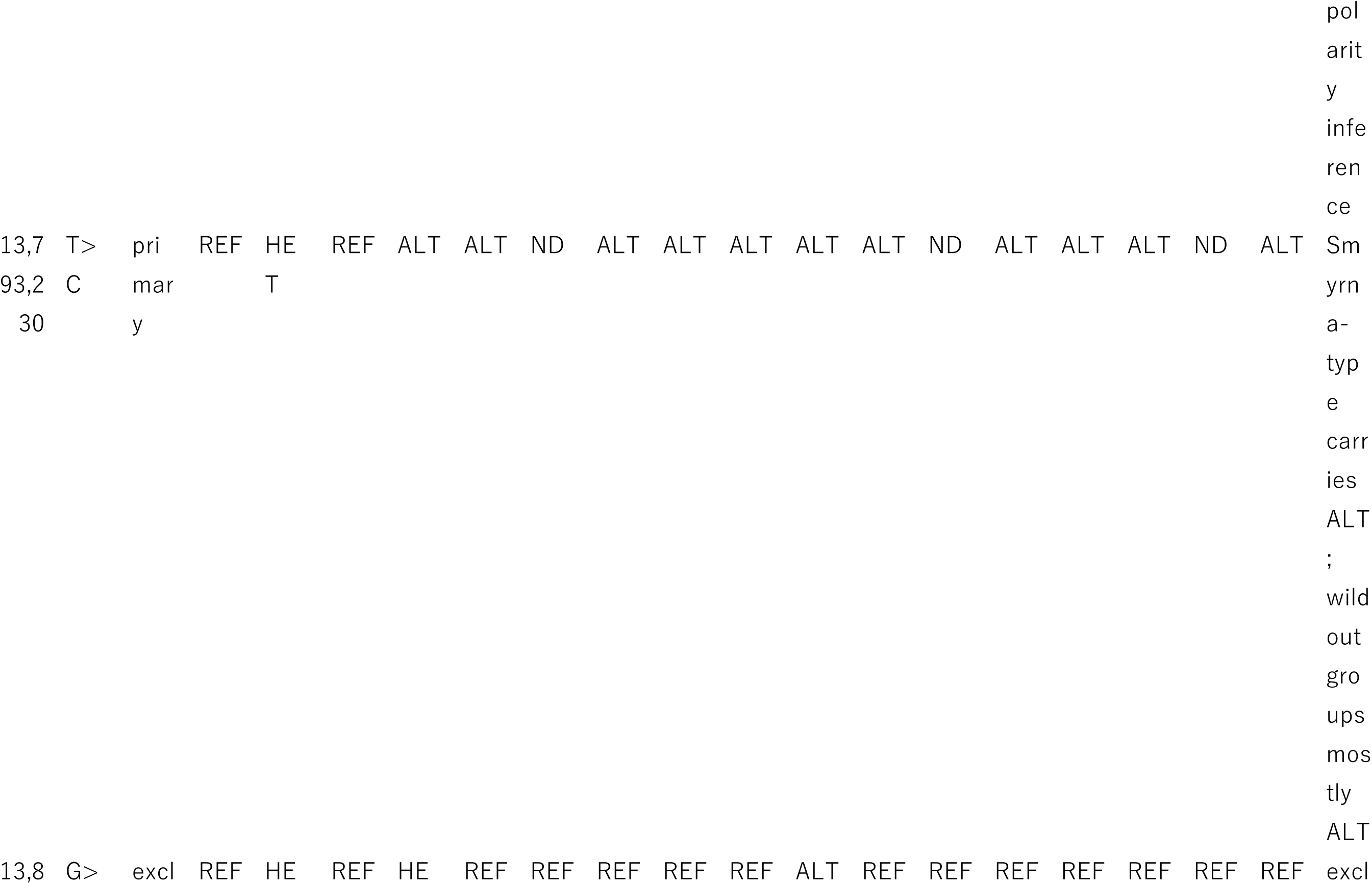
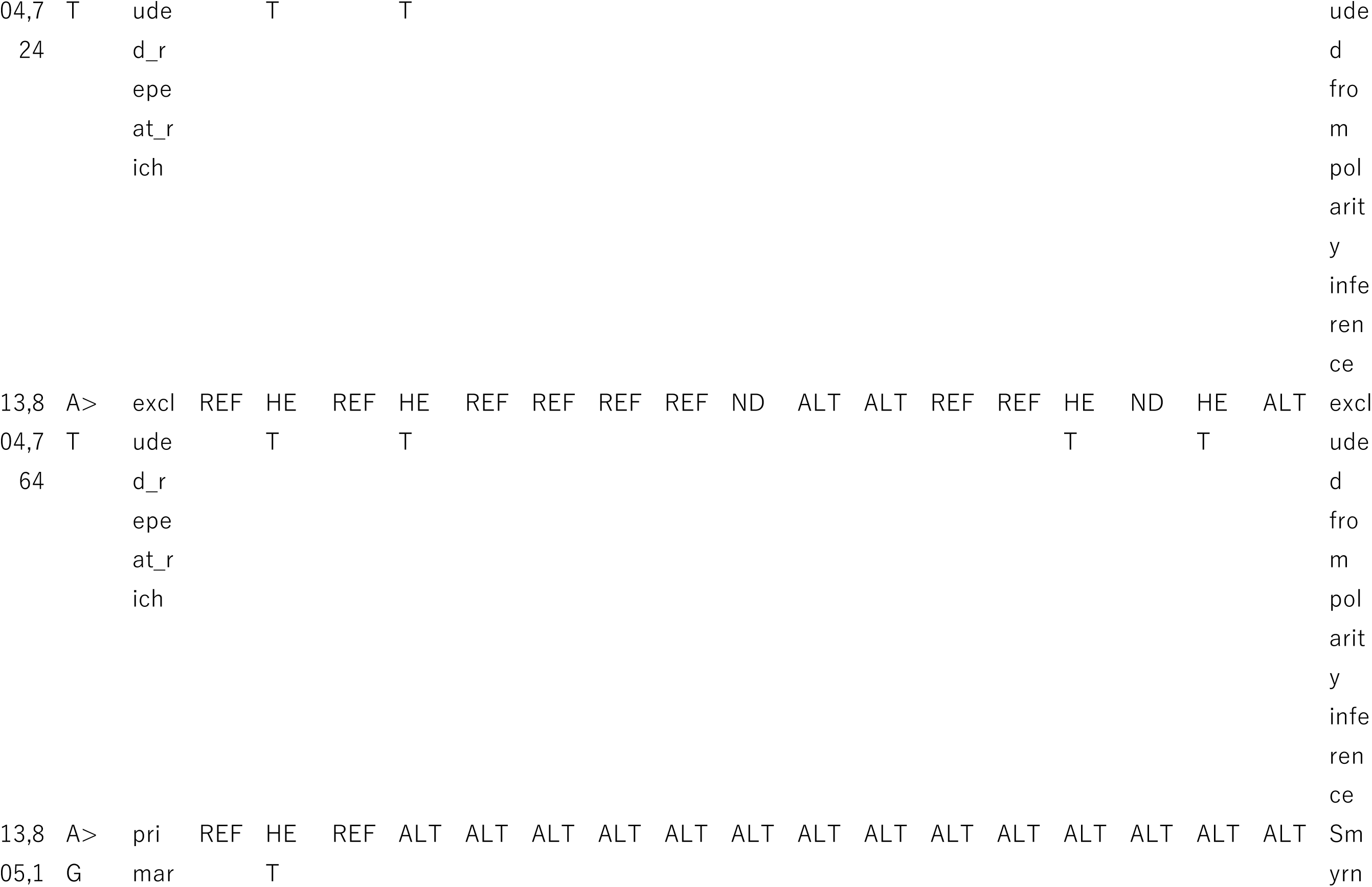
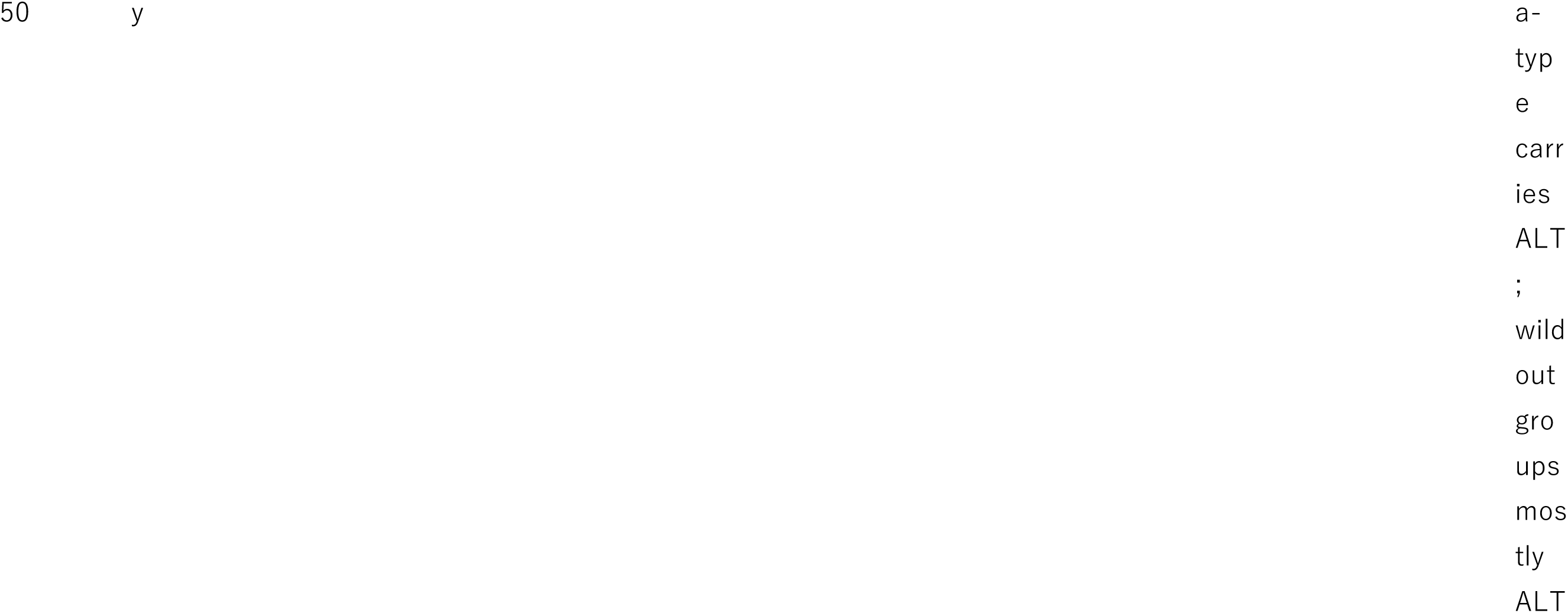
Genotypes at the eight concordant SNPs in *Ficus carica* and wild *Ficus* outgroup species. Genotype calls at the eight perfectly concordant SNPs defining the chromosome 04 *Eden* locus (chr04:13,785,367; 13,785,700; 13,788,990; 13,789,219; 13,793,230; 13,804,724; 13,804,764; 13,805,150), shown for non-Smyrna *F. carica*, Smyrna-type *F. carica*, the closely related wild species *F. palmata*, and fourteen more distantly related wild *Ficus* species (*F. erecta*, *F. adhatodifolia*, *F. auriculata*, *F. cyathistipula*, *F. cyrtophylla*, *F. esquiroliana*, *F. gasparriniana*, *F. hispida*, *F. maxima*, *F. microcarpa*, *F. oligodon*, *F. religiosa*, *F. sagittata*, and *F. vasculosa*). Genotypes were recovered by mapping publicly available whole-genome resequencing reads ^18,19^ (BioProject PRJNA684963) to the FCAr2.3 reference assembly…Four positions (chr04:13,785,367; 13,788,990; 13,793,230; 13,805,150) yielded genotype calls with adequate depth across all fourteen outgroup species and were used for polarity inference in the main text.

**Supplementary Table 4.**
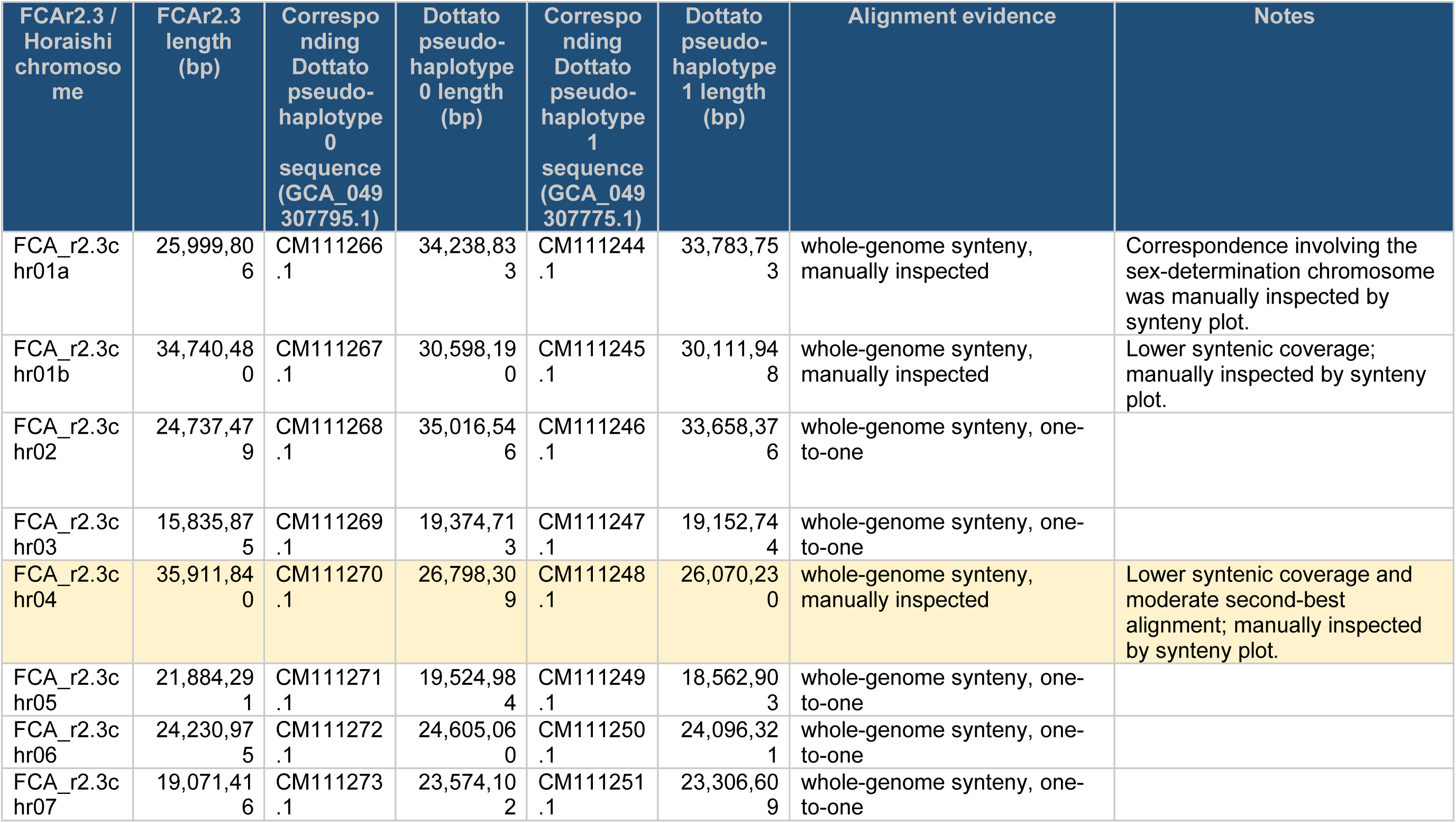
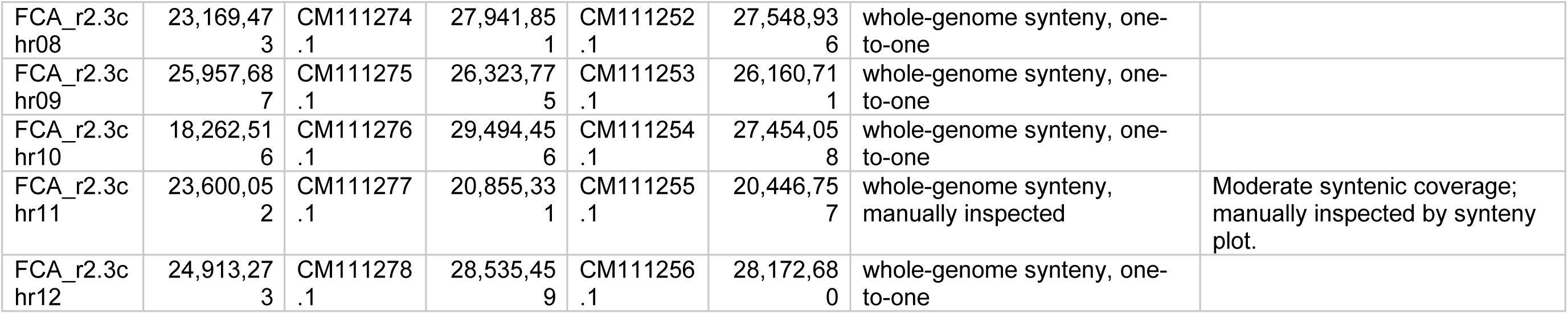
Correspondence between fig (*Ficus carica*) chromosome numbering systems. Chromosome numbers in the Horaishi-based reference assemblies^41,42^, used throughout this study, are shown alongside the corresponding sequences in the haplotype-phased Dottato assembly ^49,50^ (pseudo-haplotypes 0 and 1; GenBank assembly accessions GCA_049307795.1 and GCA_049307775.1). The two numbering systems reflect different naming conventions rather than differences in genome content: the Dottato assembly numbers chromosomes in descending order of physical length, whereas the Horaishi-based assemblies retain the numbering established in the original reference. In the Horaishi-based system, chromosome 1 is divided into two segments (chr01a and chr01b) following anchoring to the twelve chromosomes of the jujube (*Ziziphus jujuba*) genome, yielding thirteen reference entries. Correspondence was established by whole-genome alignment using minimap2 v2.28-r1209; pairs with lower syntenic coverage or a moderate second-best alignment—including the sex-determination chromosome (chr01a) and chr04—were additionally verified by manual inspection of synteny plots (see Notes). Chromosome lengths (bp) are given for reference; differences between systems reflect both cultivar divergence (Horaishi vs Dottato) and assembly version. This table enables cross-referencing of chromosome-level results between the two assemblies.

**Supplementary Figure 1.**
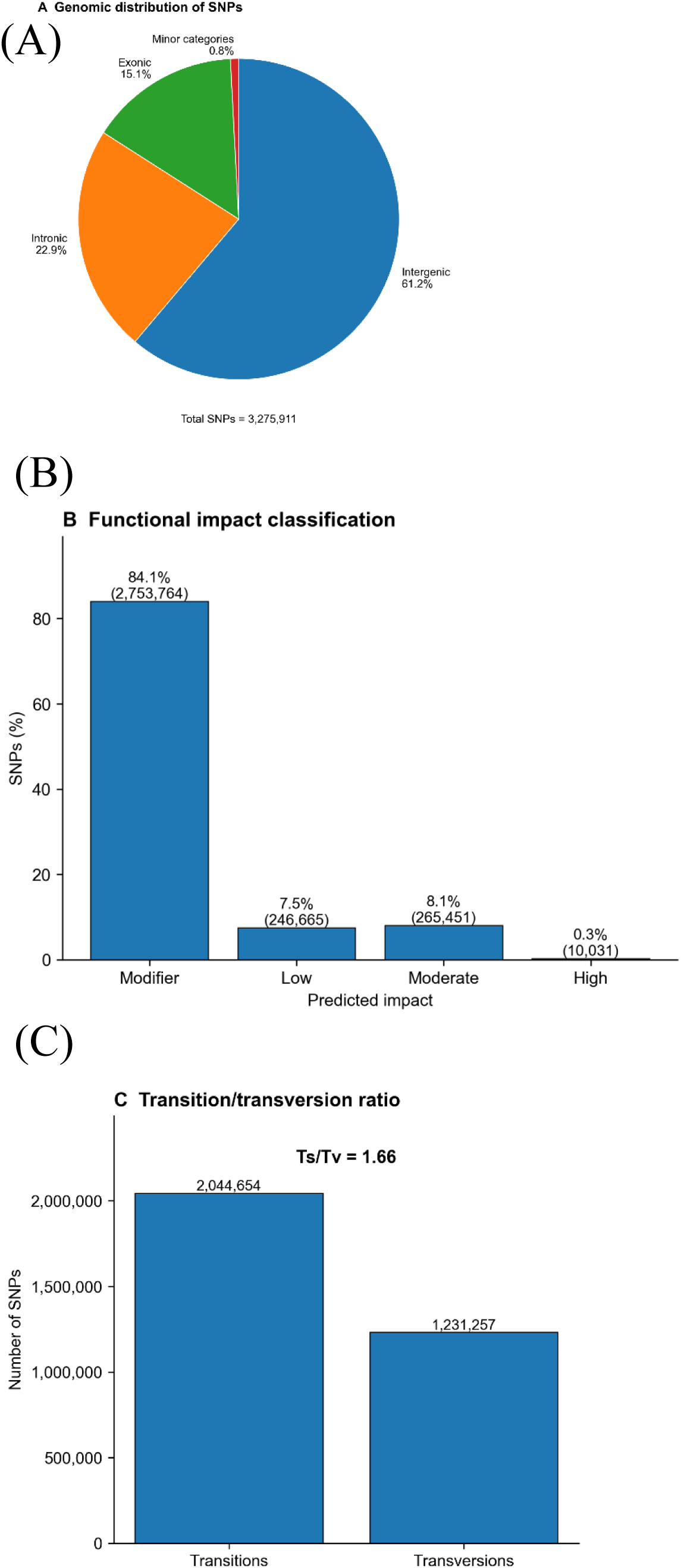
Genomic distribution and functional impact classification of SNPs identified across the *Ficus carica* accession panel. (A) Pie chart showing the proportional distribution of 3,275,911 SNPs across genomic compartments classified using SnpEff annotations: intergenic regions (61.2%), intronic regions (22.9%), exonic regions (15.1%), and minor categories (0.83%; splice-region variants, UTR/upstream/downstream variants, and other variants). For variants with multiple annotations, one representative genomic compartment was assigned according to the priority order of exonic, splice-region, intronic, UTR/upstream/downstream, intergenic, and other categories. (B) Bar chart showing the proportion of SNPs assigned to each predicted functional impact category by SnpEff: modifier (84.1%), moderate (8.1%), low (7.5%), and high (0.3%). For variants with multiple annotations, the strongest predicted impact was used for classification. (C) Transition-to-transversion ratio (Ts/Tv = 1.66), with transitions (2,044,654) and transversions (1,231,257) indicated. All statistics were derived from the filtered, non-imputed SNP dataset in FCAr2.3pmol.482.snpeff.vcf.gz, mapped to the female-derived reference assembly FCAr2.3.

**Supplementary Figure 2.**
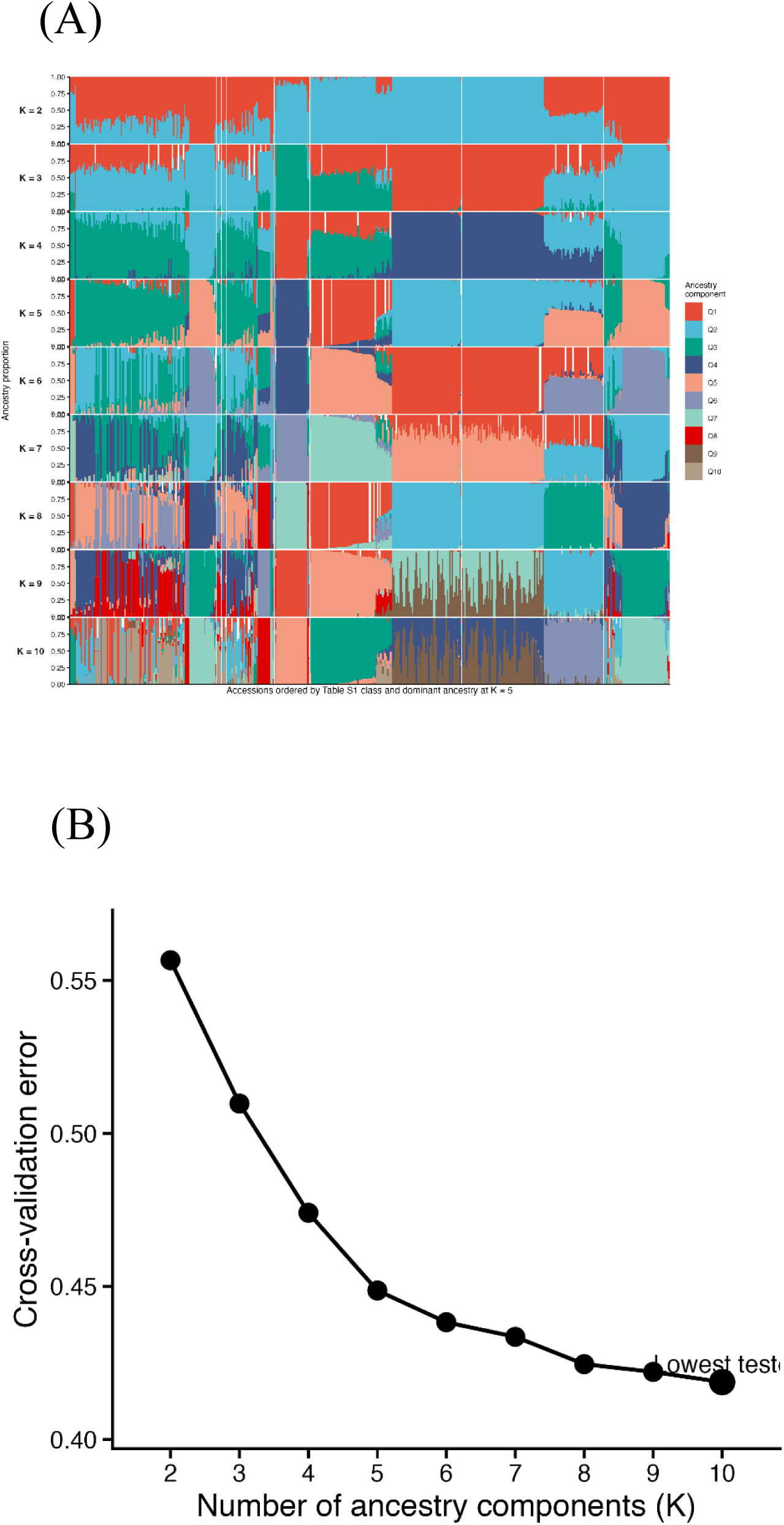
Extended ADMIXTURE analysis of the *Ficus* accession panel. (A) ADMIXTURE ancestry proportion plots for K = 2 to K = 10, based on the LD-pruned SNP dataset (164,933 SNPs). Each vertical bar represents one accession, and colours indicate inferred ancestry components within each K value. Accessions are shown in a fixed order across K values based on the sample order in the PLINK .fam file, allowing comparison of population structure across different numbers of ancestry components. (B) Cross-validation error across tested K values (K = 2–10) in the ADMIXTURE analysis. Cross-validation error decreased monotonically across the entire tested range, with the lowest value observed at K = 10 (the maximum tested K). The absence of a clear minimum within the tested range suggests that the accession panel harbours fine-scale population substructure that was not fully resolved at K ≤ 10, likely reflecting the continuous and admixed nature of genetic variation across the *Ficus* accession panel documented by PCA and neighbour-joining tree analysis.

**Supplementary Figure 3.**
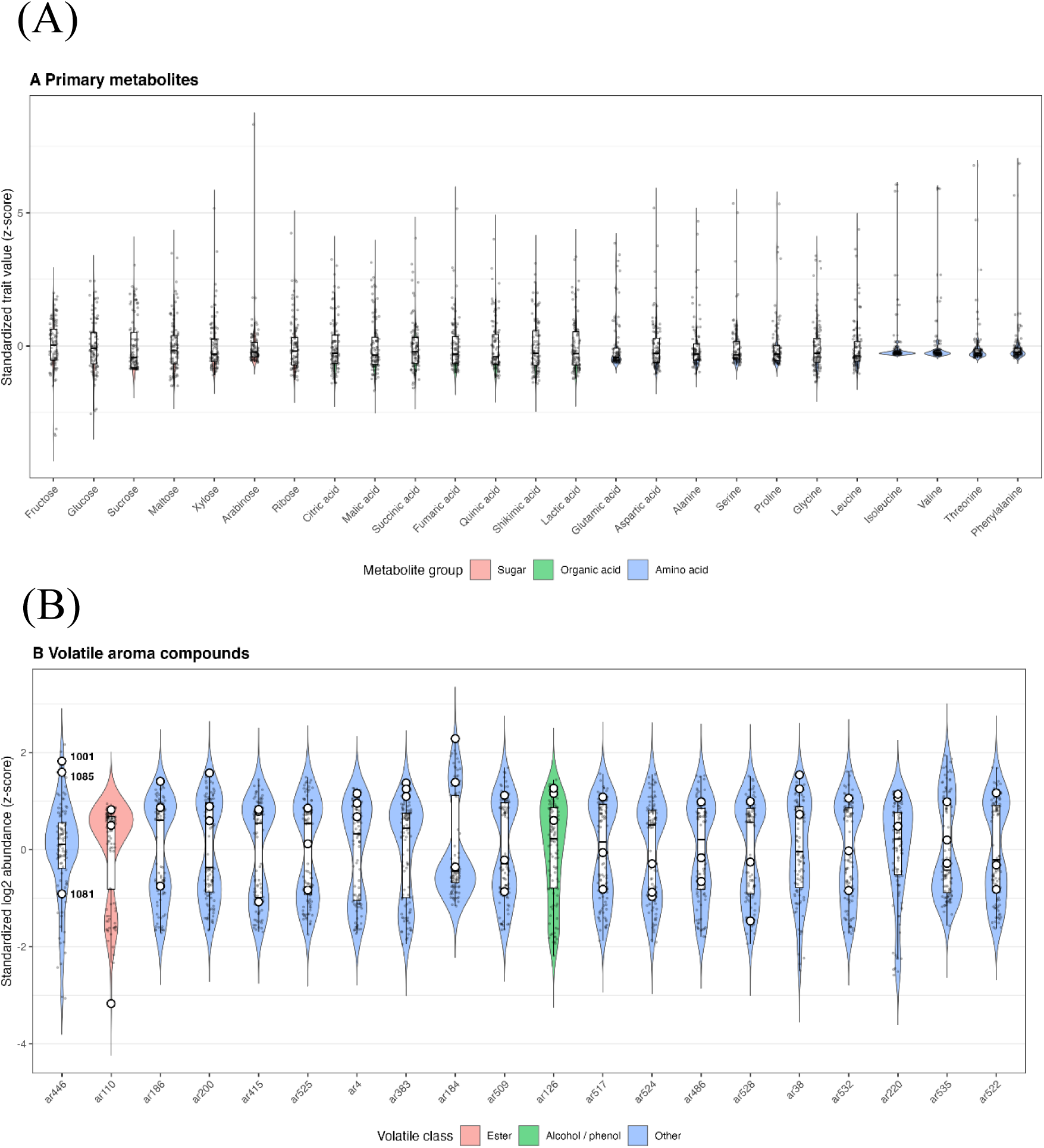
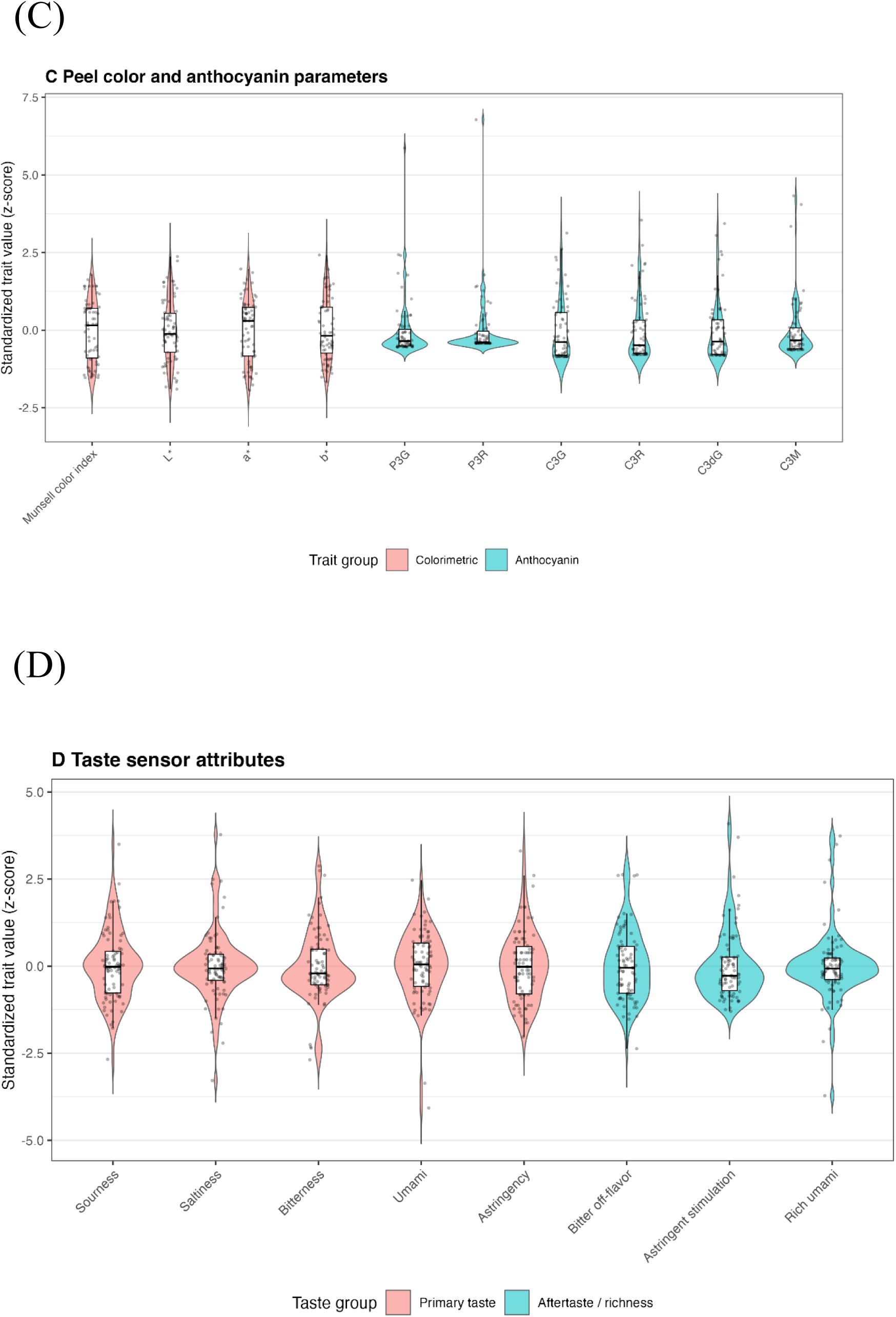
Distribution of phenotypic variation across four fruit quality omics layers in the *Ficus carica* accession panel. Violin plots and overlaid box plots showing the distribution of standardised trait values (Z-score) across accessions for representative features from each of the four omics layers. (A) Primary metabolites: distributions of sugars (fructose, glucose, sucrose, maltose, xylose, arabinose, ribose), organic acids (citric acid, malic acid, succinic acid, fumaric acid, quinic acid, shikimic acid, lactic acid, glutamic acid, aspartic acid), and amino acids (alanine, serine, proline, glycine, leucine, isoleucine, valine, threonine, phenylalanine), coloured by metabolite group (sugar, organic acid, amino acid). (B) Volatile aroma compounds: distributions of the top 20 most variable volatile features, including ar446. High-impact aroma variant accessions (1081, 1001, and 1085) are overlaid as individual labelled points. Features are coloured by volatile class (ester, alcohol/phenol, or other). (C) Peel colorimetric and anthocyanin parameters: distributions of Munsell color index, L*, a*, b*, and individual anthocyanin species concentrations (P3G, P3R, C3G, C3R, C3dG, C3M), coloured by trait group (colorimetric or anthocyanin). (D) Taste sensor attributes: distributions of all eight taste dimensions—sourness, saltiness, bitterness, umami, astringency, bitter off-flavor, astringent stimulation, and rich umami—coloured by taste group (primary taste or aftertaste/richness). This figure complements the PCA-based overview presented in Figure 2.

**Supplementary Figure 4.**
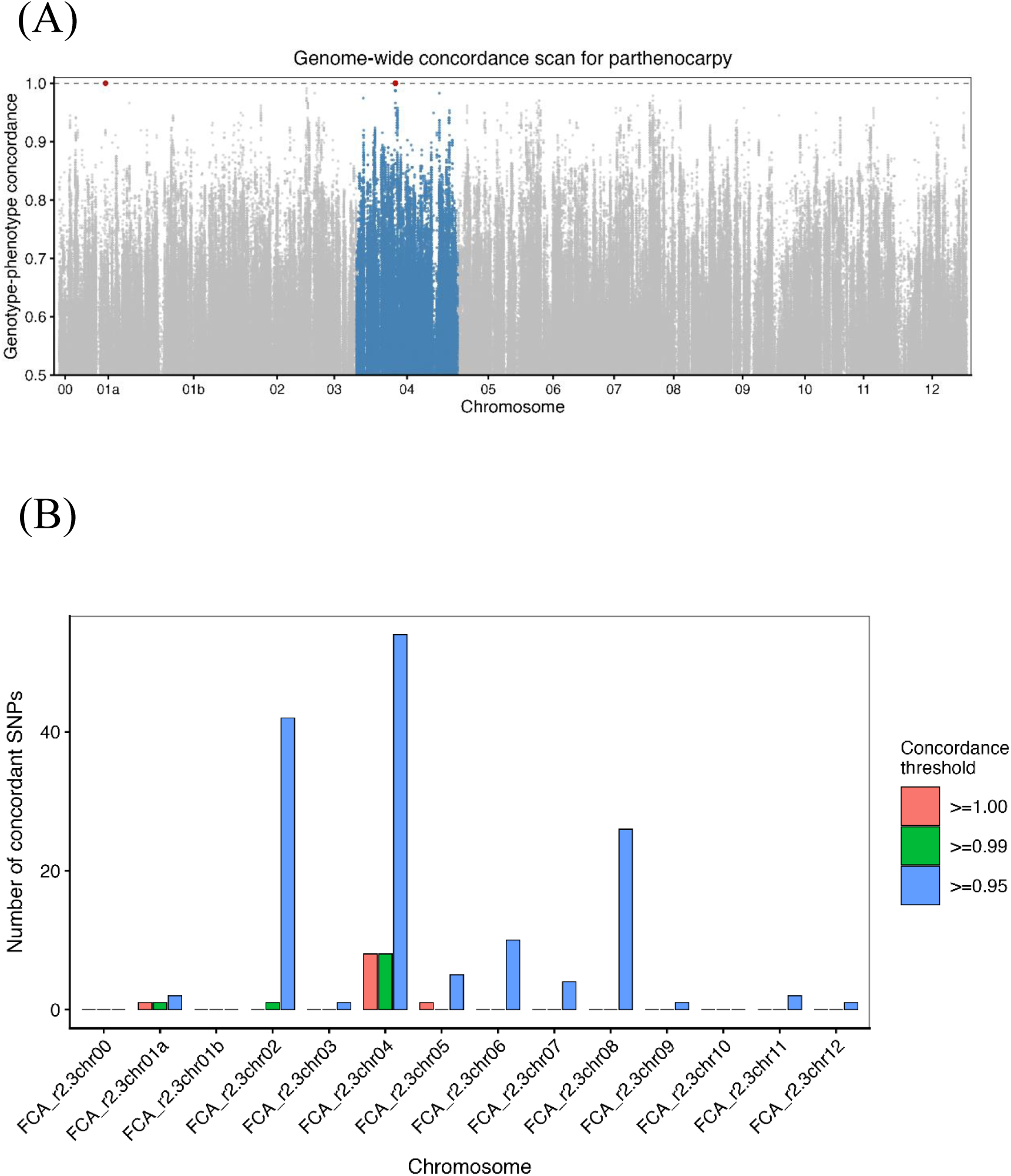
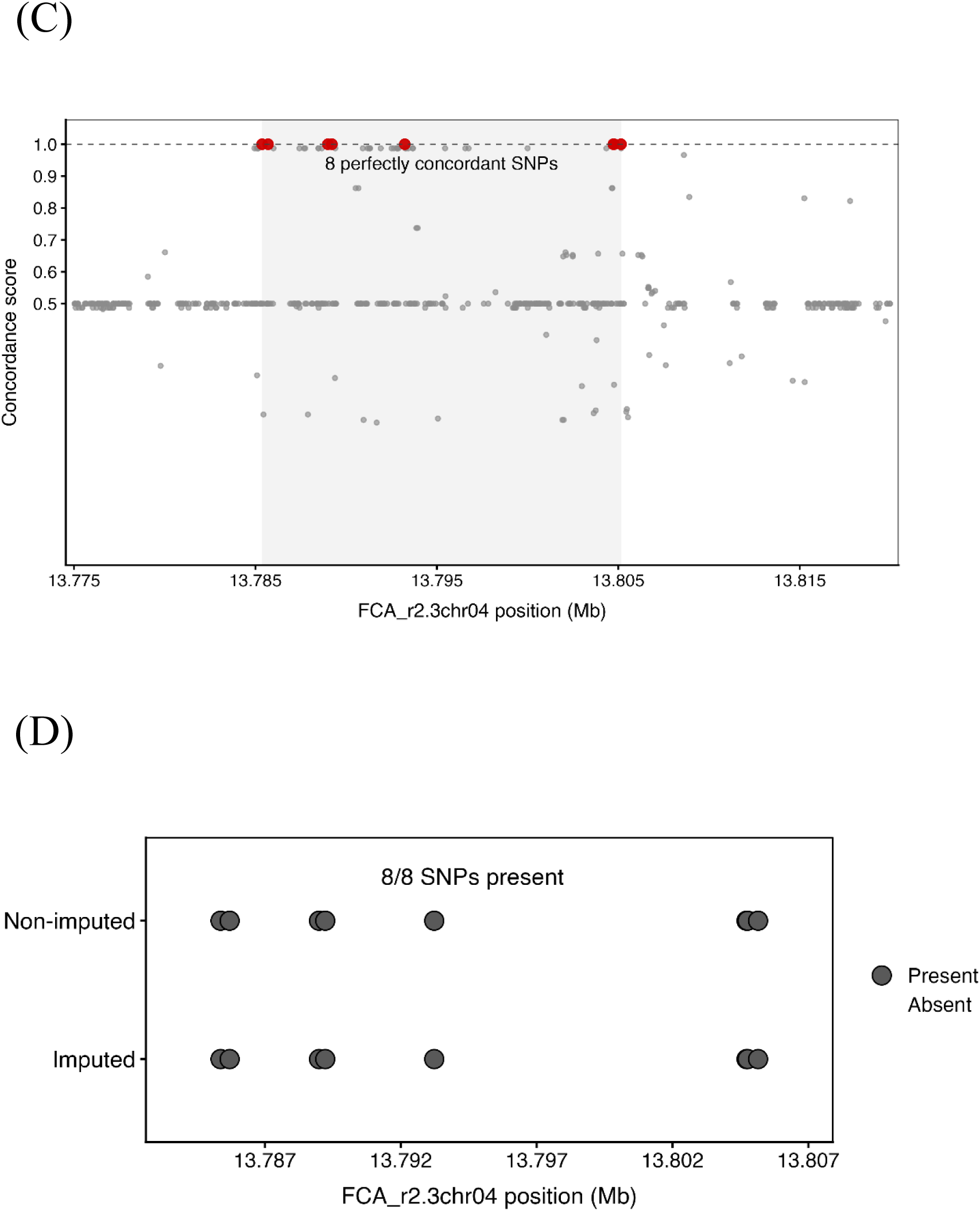
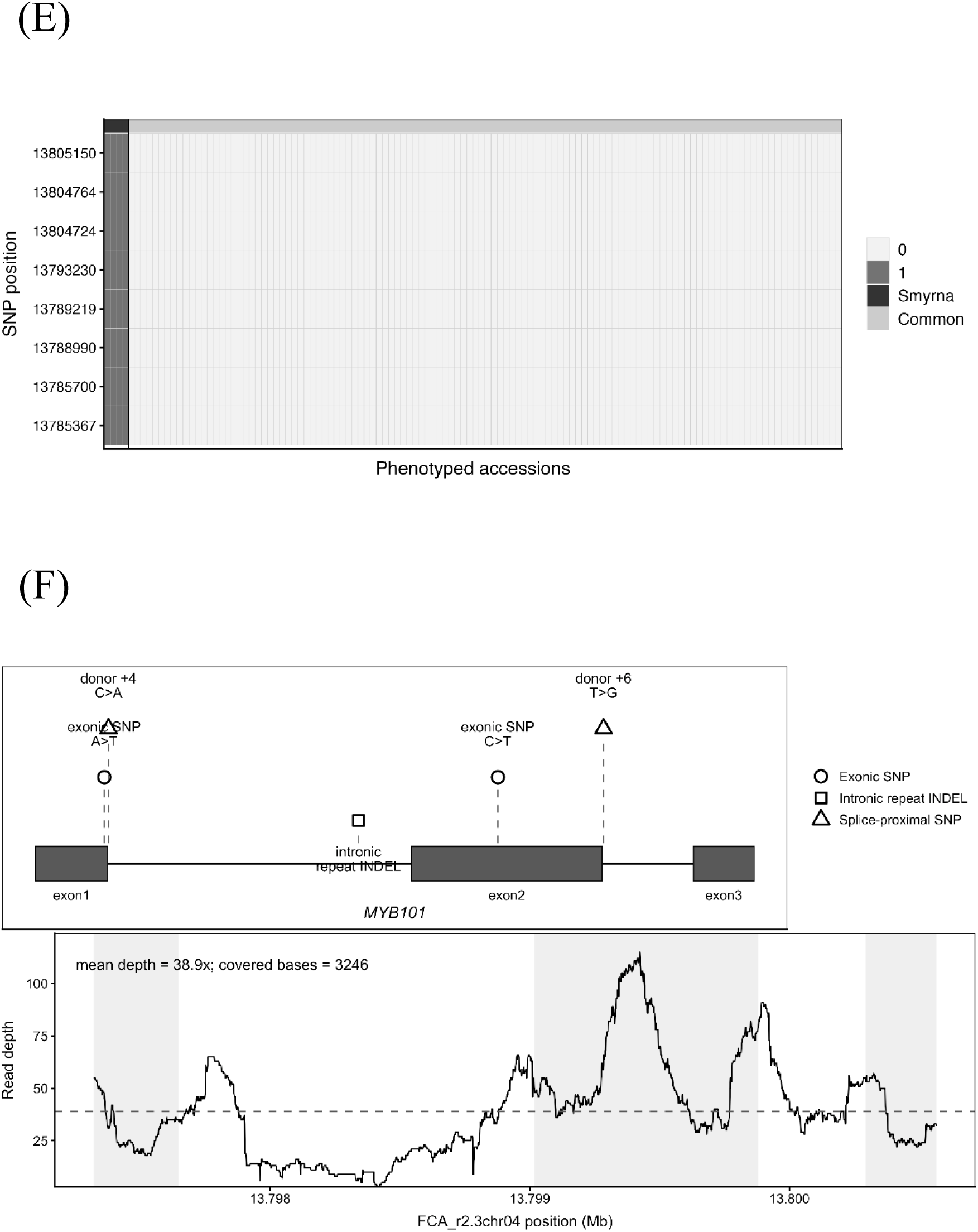
Supporting concordance, validation, and read-based inspection analyses for the chromosome 04 *MYB101-like*-centred parthenocarpy candidate interval. (A) Genome-wide genotype–phenotype concordance scan for parthenocarpy using the imputed FCAr2.3pmol SNP dataset. Each point represents a SNP plotted by chromosomal position and genotype–phenotype concordance score. The dashed horizontal line indicates complete concordance (score = 1.0). The *MYB101-like*-centred complete-concordance haplotype on chromosome 04 is highlighted in red. (B) Chromosome-level summary of perfectly concordant and highly concordant SNPs. The number of SNPs exceeding each genotype–phenotype concordance threshold (score = 1.00, ≥ 0.98, or ≥ 0.95) was counted for each chromosome. Eight perfectly concordant SNPs were located on chromosome 04, with one additional isolated perfectly concordant SNP on chromosome 01a and one on chromosome 05. This spatial concentration on chromosome 04, in contrast to the isolated signals on other chromosomes, supports chromosome 04 as the primary complete-concordance signal. (C) Detailed regional view of the chromosome 04 concordance interval (13.785–13.810 Mb) surrounding the *MYB101-like* candidate region. SNP-level concordance scores are shown together with local gene annotations indicating the relative positions of the perfectly concordant SNPs and the three annotated genes in the interval: (g020640.1), *MYB101-like* (g020650.1), and 2-alkenal reductase (g020660.1). The shaded region indicates the candidate interval (13.785–13.810 Mb). (D) Verification of the eight perfectly concordant chromosome 04 SNPs in the non-imputed FCAr2.3pmol SNP dataset. All eight SNPs identified from the imputed dataset were also present as observed variants in the non-imputed dataset (8/8 SNPs present), confirming that the concordant haplotype is not an artefact of genotype imputation. (E) Genotype matrix of the eight perfectly concordant chromosome 04 SNPs across phenotyped accessions. Rows represent the eight SNPs in the *MYB101-like*-centred candidate interval (positions in FCAr2.3 coordinates shown on the y-axis), and columns represent accessions grouped by reproductive phenotype (Smyrna-type controls on the left; Common and San Pedro-type parthenocarpic cases on the right). Cell colour indicates mean genotype dosage. The four Smyrna-type controls consistently carry dosage 1 (heterozygous), whereas all 118 Common and San Pedro-type accessions carry dosage 0, yielding complete genotype–phenotype separation at all eight SNP positions. (F) Read-based inspection of the *MYB101-like* gene body in a representative Smyrna-type control accession (Fc_57). The upper panel shows the *MYB101-like* gene model (three exons) and variants detected from Fc_57 read mapping: two exonic SNPs (A>T in exon 1; C>T in exon 2), two splice-proximal SNPs at the donor sites of exons 1 and 2 (donor +4 C>A; donor +6 T>G), and one intronic repeat-length INDEL. The lower panel shows read depth across the *MYB101-like* gene body (mean depth 38.9×; covered bases 3,246). No exon-disrupting frameshift INDEL was detected; however, the splice-proximal SNPs at donor +4 and donor +6 positions may affect splicing efficiency and warrant functional investigation.

**Supplementary Figure 5.**
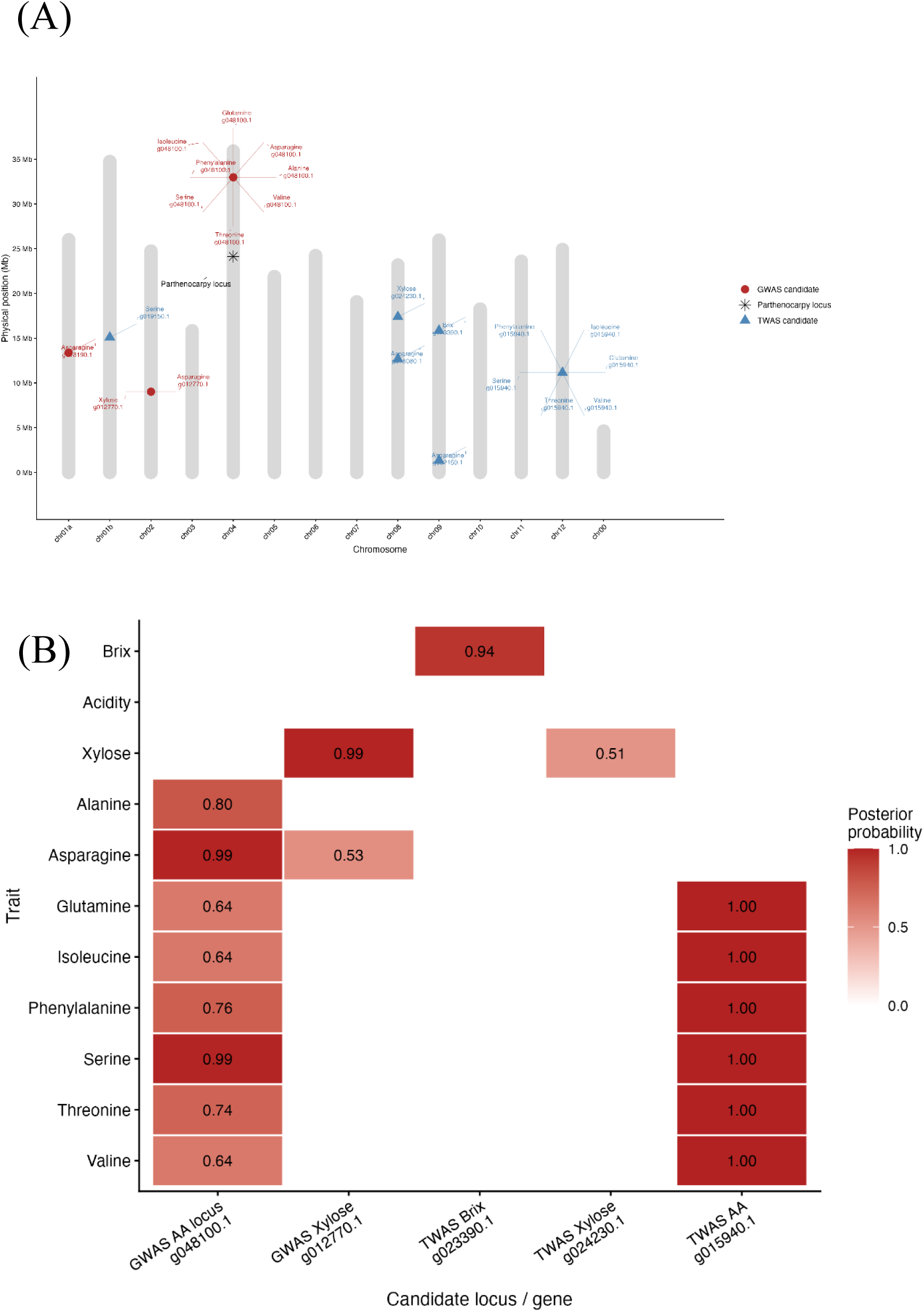
Chromosomal distribution and posterior probability summary of GWAS/TWAS candidate loci for primary metabolite traits. (A) Chromosomal map of high-confidence GWAS and TWAS candidate loci for primary metabolite traits. GWAS candidates are shown as circles, TWAS candidates as triangles, and the chromosome 04 parthenocarpy candidate locus (MYB101-like concordant haplotype; 13.785–13.810 Mb) as a star. Associated traits and candidate gene IDs are annotated around each locus. Chromosomal positions are shown in physical megabase coordinates based on FCAr2.3 pseudomolecule lengths. All primary metabolite-associated loci are spatially distinct from the parthenocarpy locus on chromosome 04. (B) Candidate–trait posterior probability heatmap summarising key GWAS and TWAS signals for primary metabolite traits. The TWAS candidate g023390.1 was associated with Brix (PP = 0.937). The GWAS amino acid locus g048100.1 showed high posterior probabilities across eight amino acid traits (PP = 0.64–0.99). The TWAS candidate g015940.1 showed PP = 1.000 for glutamine, isoleucine, phenylalanine, serine, threonine, and valine, and PP = 0.802 for 2-aminoadipic acid. Xylose-associated candidates include GWAS locus g012770.1 (PP = 0.993) and TWAS candidate g024230.1 (PP = 0.510).

**Supplementary Figure 6.**
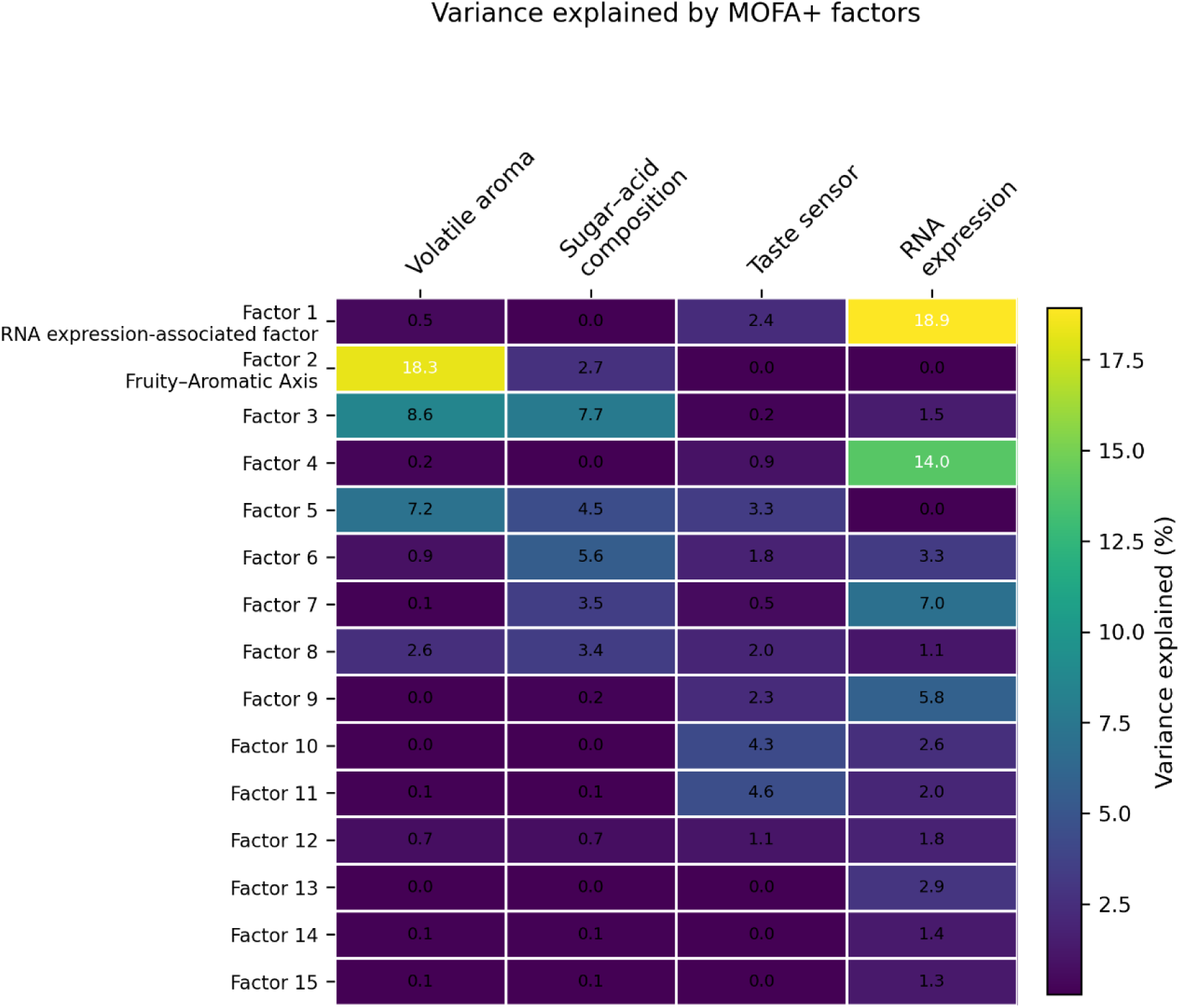
Variance explained by MOFA+ factors across four omics layers. Heatmap showing the proportion of variance explained by each of the 15 MOFA+ factors across four omics layers: volatile aroma, sugar–acid composition, taste sensor, and RNA expression. Rows represent MOFA+ factors and columns represent omics layers. Colour intensity represents the percentage of variance explained, with brighter colours indicating higher values. Factor 1 (RNA expression-associated factor) was dominated by RNA expression (18.9%), likely reflecting the high dimensionality of the transcriptomic layer. Factor 2 (Volatile Aroma Axis) was dominated by volatile aroma (18.3%) with secondary contribution from sugar–acid composition (2.7%), and corresponds to the primary sensory axis highlighted in Figure 5B and 4C. Factor 5, which integrates volatile aroma (7.2%), sugar–acid composition (4.5%), and taste sensor profiles (3.3%), corresponds to the Integrated Flavour Axis shown in Figure 5B. Factors are ordered by their assigned number; the two biologically interpreted factors highlighted in Figure 5 are labelled accordingly.

**Supplementary Figure 7.**
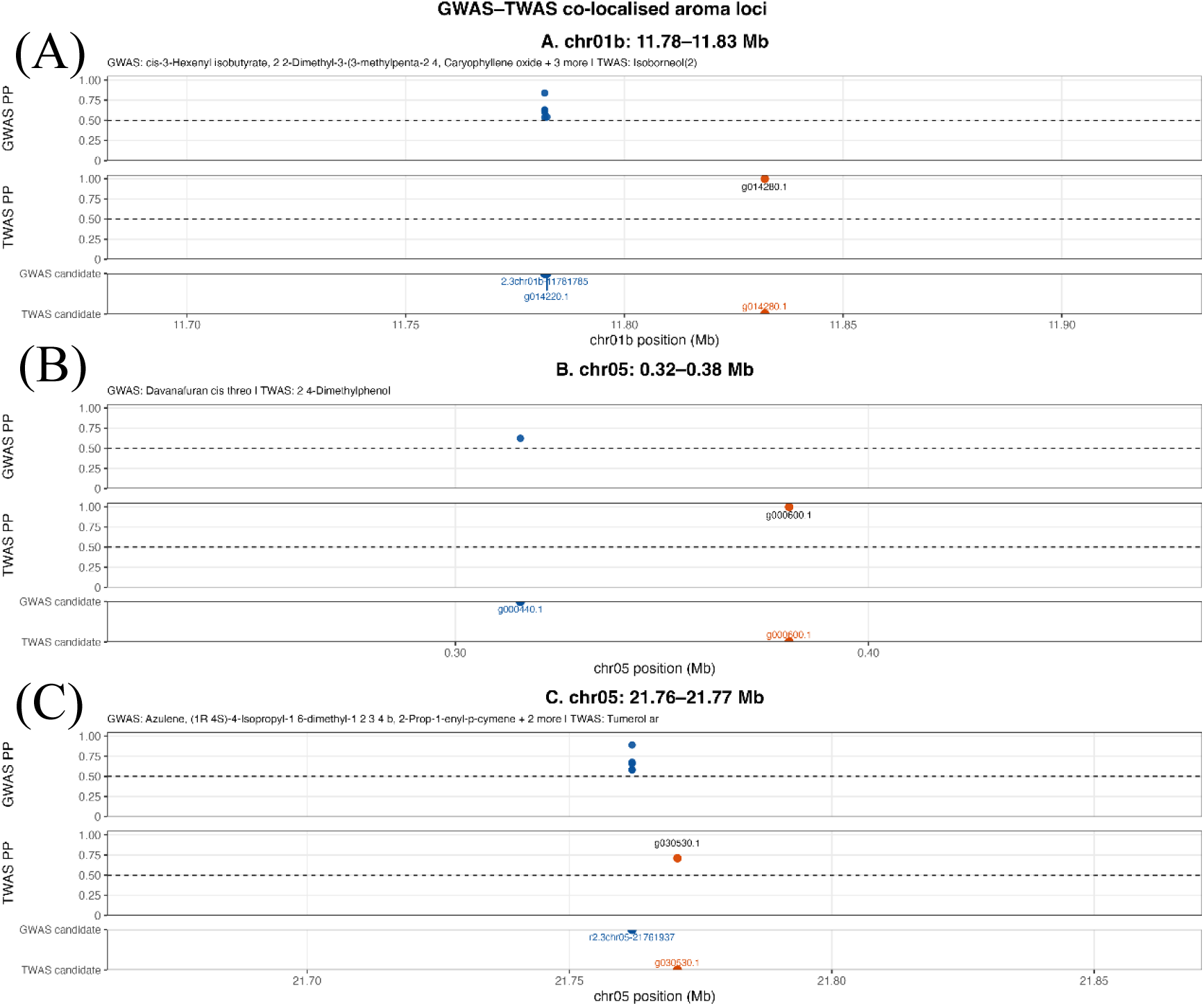
Regional views of GWAS–TWAS co-localised aroma loci. Regional association plots for three genomic loci in which GWAS and TWAS candidates for volatile aroma traits were located within ±100 kb. For each locus, the upper panel shows the GWAS posterior probability profile, the middle panel shows the TWAS posterior probability profile, and the lower panel shows the positions of GWAS (blue) and TWAS (orange) candidates. Dashed horizontal lines indicate the posterior probability threshold of PP = 0.5. (A) chr01b: 11.78–11.83 Mb. GWAS candidates for cis-3-Hexenyl isobutyrate, 2,2-Dimethyl-3-(3-methylpenta-2,4-dienyl)oxirane, Caryophyllene oxide, and related compounds (g014220.1) co-localised with the TWAS candidate g014280.1, associated with Isoborneol. (B) chr05: 0.32–0.38 Mb. GWAS candidate g000440.1 for Davanafuran cis-threo co-localised with the TWAS candidate g000600.1 associated with 2,4-Dimethylphenol. (C) chr05: 21.76–21.77 Mb. GWAS candidates for Azulene, (1R,4S)-4-Isopropyl-1,6-dimethyl-1,2,3,4b-tetrahydrophenanthrene, 2-Prop-1-enyl-p-cymene, and related compounds (r2.3chr05-21761937) co-localised with the TWAS candidate g030530.1 associated with Tumerol ar. This figure complements the genome-wide overview of aroma-associated loci shown in Figure 7A.

**Supplementary Figure 8.**
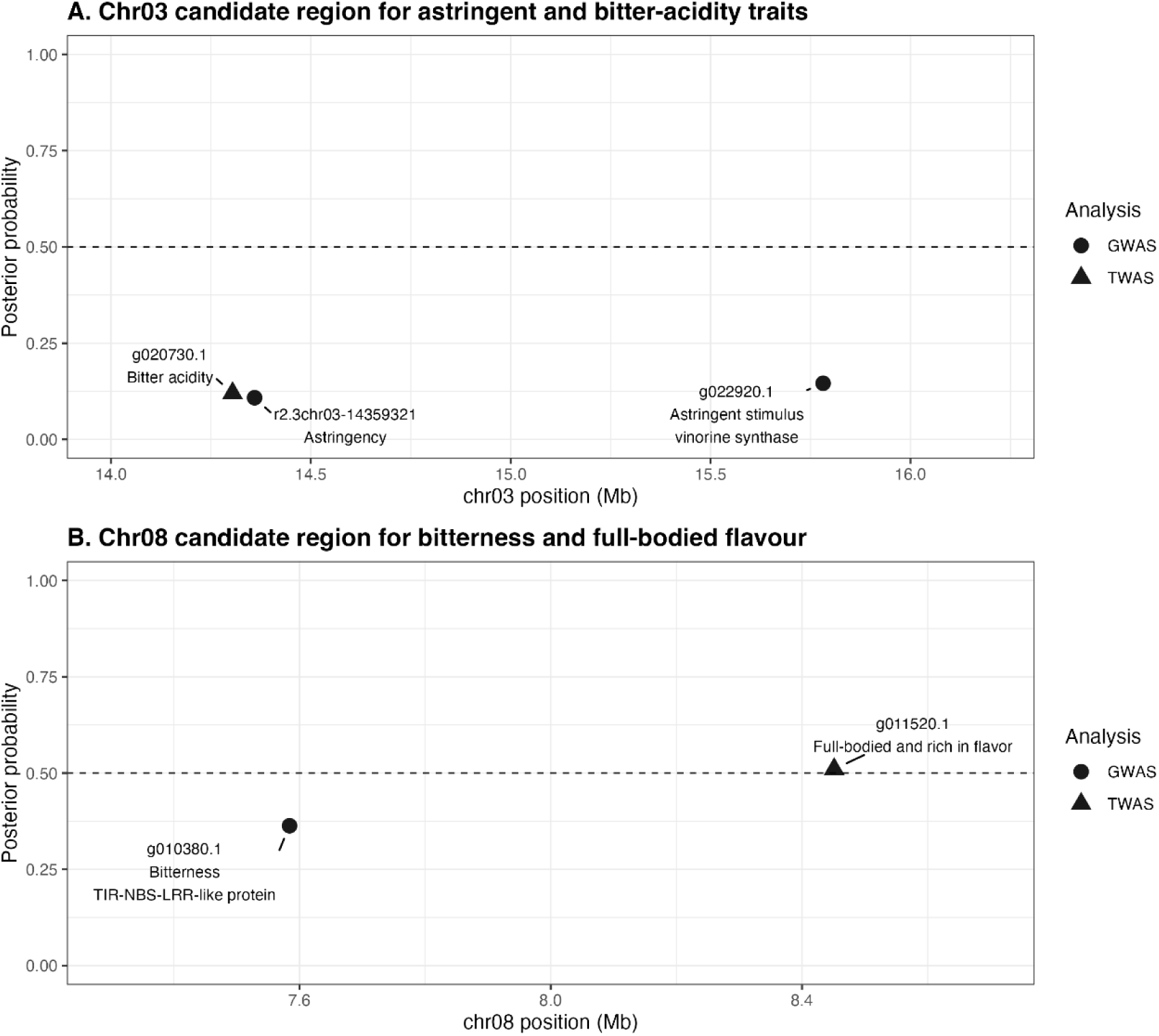
Regional association plots for taste attribute candidate regions on chromosomes 03 and 08. (A) Chromosome 03 candidate region (14.3–15.7 Mb) associated with astringent and bitter-acidity traits. Upper panel: GWAS posterior probabilities for bitter acidity (g020730.1) and astringency (r2.3chr03-14359321). Lower panel: TWAS posterior probabilities, with g022920.1 (vinorine synthase; Astringent stimulus) as the highest-confidence candidate (PP ≈ 0.12). Note that all candidates in this region fall below the PP = 0.5 threshold and are considered suggestive rather than high-confidence associations. (B) Chromosome 08 candidate region associated with bitterness and full-bodied flavour attributes. Upper panel: GWAS candidate g010380.1 (TIR-NBS-LRR-like protein; Bitterness; PP ≈ 0.35) at ∼7.6 Mb. Lower panel: TWAS candidate g011520.1 (uncharacterized protein; Full-bodied and rich in flavor; PP = 0.51) at ∼8.4 Mb. The GWAS and TWAS candidates are located ∼0.85 Mb apart within the same chromosomal region, suggesting a broader regulatory interval for complex flavour perception on chromosome 08. In each panel, the dashed horizontal line indicates PP = 0.5.

**Supplementary Figure 9.**
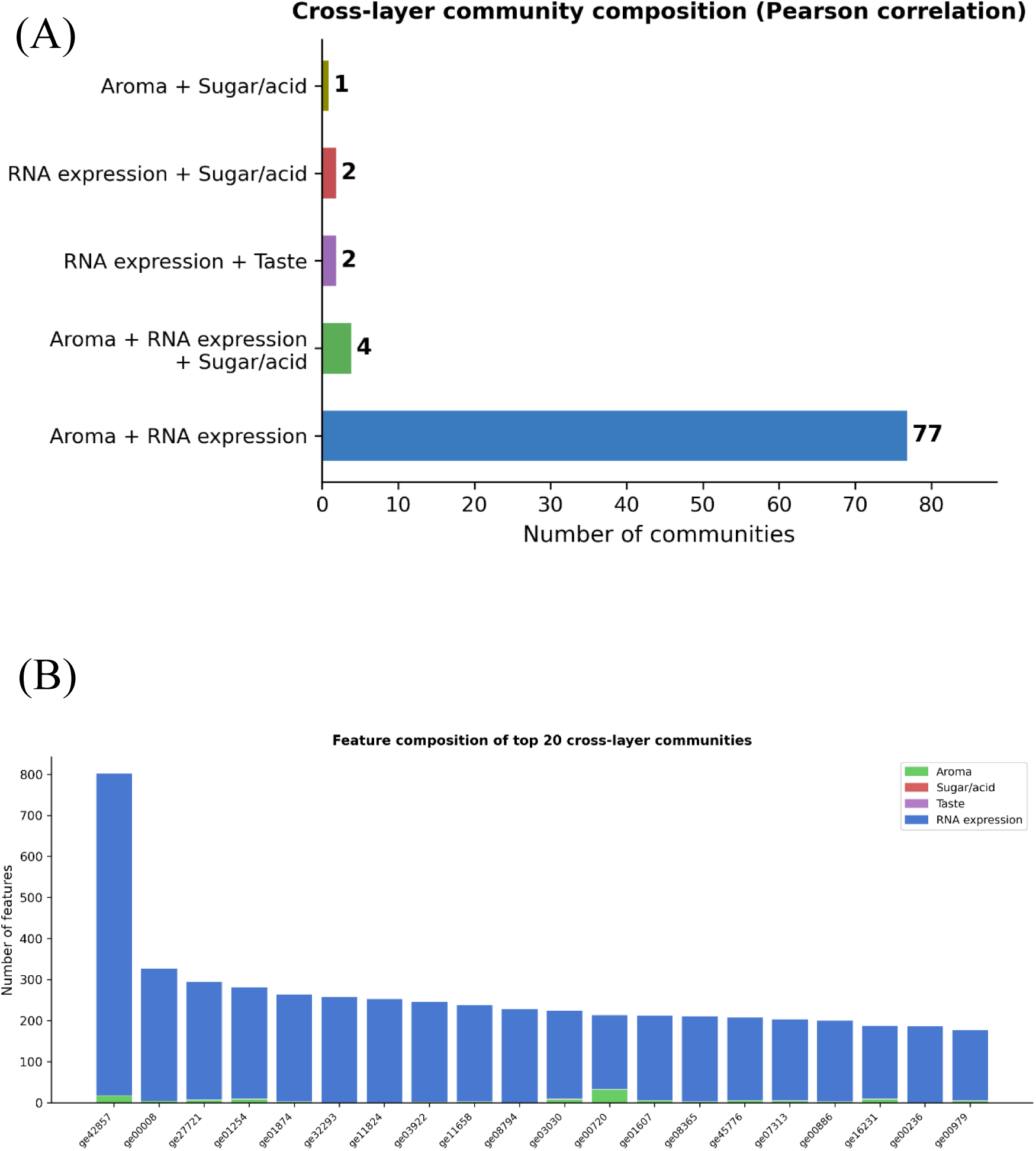
Cross-layer community structure across four omics layers revealed by ConfeitoGUI (Pearson correlation). Correlation network analysis was performed using ConfeitoGUI across the same four omics layers analysed by MOFA+ (volatile aroma, sugar–acid composition, taste sensor profiles, and RNA expression; n = 81 accessions). (A) Number of communities spanning two or more omics layers, classified by layer combination. Among 86 multi-layer communities, the most frequent combination linked aroma features with RNA expression (77 communities), with smaller numbers incorporating sugar–acid composition or taste sensor features. The predominance of multi-layer communities indicates that the four omics layers are interconnected rather than forming layer-specific modules. (B) Feature composition of the top 20 cross-layer communities ranked by community size. Each bar represents one community; colours indicate the omics layer of each constituent feature (Aroma, green; Sugar/acid, red; Taste, purple; RNA expression, blue). Community representative IDs are shown on the x-axis. The mixed-layer composition of these communities confirms that features from different omics layers co-occur within shared correlation modules, consistent with the integrated flavour structure identified by MOFA+.

**Supplementary Figure 10.**
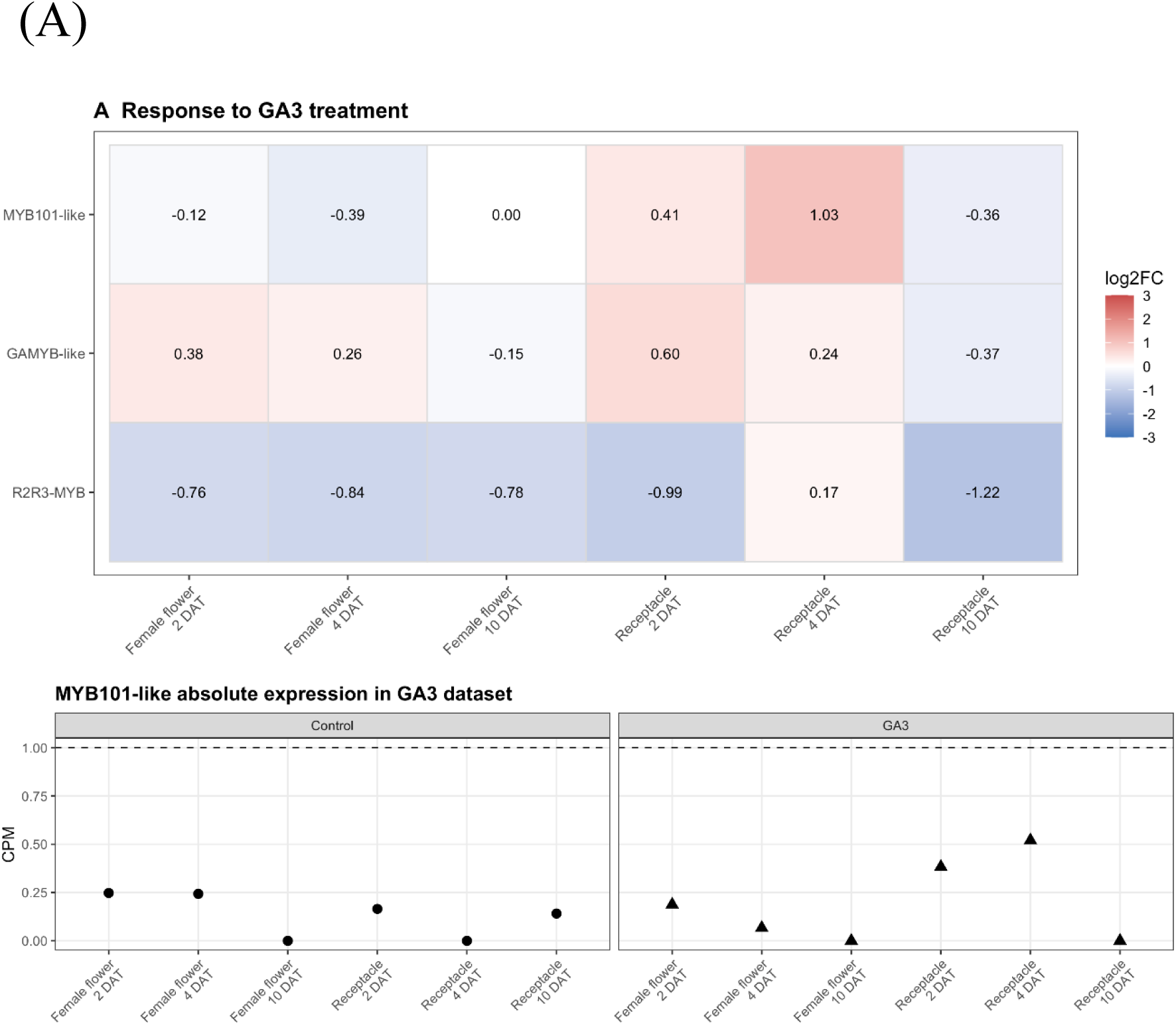
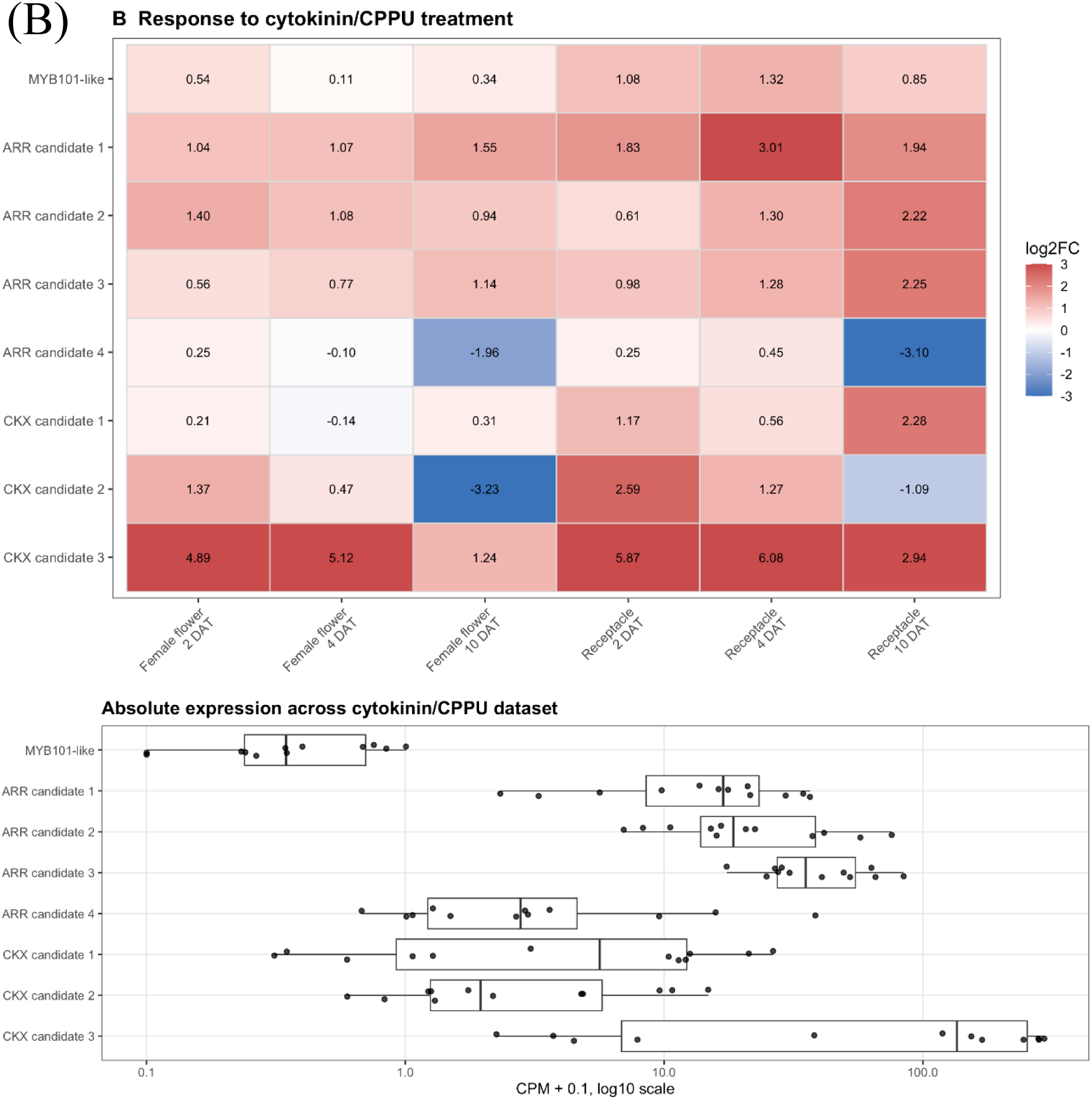
Exploratory expression analysis of *MYB101-like* in publicly available hormone-treatment RNA-seq datasets from San Pedro-type fig. Publicly available RNA-seq reads from San Pedro-type fig (*Ficus carica* L.) main crop treated with exogenous gibberellin (GA₃; SRA SRP113799)^24^ or cytokinin (CPPU; SRA PRJNA397979)^47^ were re-mapped to the FCAr2.3 reference assembly using a uniform pipeline, and expression of *MYB101-like* (Fca_r2.3chr04_g020650.1) was quantified alongside related and control genes. In both datasets, a single sequencing library is available per condition (tissue × treatment × timepoint); no biological replicates are deposited. Consequently, no replicate-based statistical testing could be performed, and all values shown are descriptive only. (A) Response to GA₃ treatment. log2 fold-change (GA₃ / control) for *MYB101-like* (Fca_r2.3chr04_g020650.1), its closest full-length GAMYB-like paralog (Fca_r2.3chr00_g005240.1), and a phylogenetically distinct R2R3-MYB (Fca_r2.3chr04_g042460.1), across female flower and receptacle tissues at 2, 4, and 10 days after treatment (DAT). *MYB101-like* transcript levels remained near the detection limit (normalized counts < 1 across all conditions; see side panel) and showed no coherent directional response to GA₃. (B) Response to cytokinin (CPPU) treatment. log2 fold-change (cytokinin / control) for *MYB101-like* and seven cytokinin-responsive positive control genes, identified as fig orthologs of characterized Arabidopsis type-A response regulators (ARR3/4/5/6/7/15/16) and cytokinin oxidase/dehydrogenase (CKX) family members by reciprocal sequence similarity, across the same six tissue × timepoint conditions. Positive control genes showed robust and largely consistent induction, confirming that the pipeline captures genuine cytokinin-responsive transcriptional signal. Against this background, *MYB101-like* showed a directionally consistent increase across all six comparisons, although its absolute expression remained two to three orders of magnitude lower than the positive controls. Gene identifiers refer to the FCAr2.3 annotation. Absolute expression values (counts per million, CPM) for all genes and conditions are provided in Supplementary Data 4. Because these observations derive from single-library comparisons, they are presented as exploratory, hypothesis-generating results and are not used to support statistical claims in the main text.

## Data File

**Supplementary Data 1. Accession composition and multi-omics analysis participation of the *Ficus* accession panel.**

The panel comprises 482 accessions subjected to whole-genome resequencing, including 217 newly sequenced *Ficus carica* accessions—165 cultivars and parental lines and 52 F1 progeny from crosses within *F. carica*; by horticultural class, 118 Common-type, 38 Caprifig, 4 San Pedro-type, 4 Smyrna-type, and 53 of unassigned class—together with 1 *F. palmata* (sister species of *F. carica* within subsect. *Ficus*; shares the parthenocarpic d-type haplotype and was excluded from the parthenocarpy concordance scan as it cannot be classified by *F. carica* horticultural criteria), 1 *F. erecta*, 1 *F. carica* × *F. erecta* interspecific hybrid (FEBN7), two *F. carica* × *F. erecta* interspecific hybrid populations (FcB1F1, n = 121; FcB2F1, n = 114), and 27 previously published *Ficus* genomes representing wild and closely related *Ficus* species^18,19^. For each accession, the table reports accession ID, sex phenotype, species/cross type, horticultural class or population, material origin, and participation status (Yes/No) in each analysis: WGS, fruit morphology/color assessment, sugar–acid profile, taste sensor profile, volatile aroma profile, RNA-seq, GWAS, TWAS, and MOFA+.

**Supplementary Data 2. Full list of GWAS and TWAS candidate loci for volatile aroma compounds, peel colorimetric parameters, and anthocyanin traits.**

Candidate loci identified by Bayesian GWAS and TWAS for volatile aroma compounds (n = 241 entries; 126 GWAS and 115 TWAS across 84 unique candidate genes), peel colorimetric parameters (n = 10 entries), and individual anthocyanin species (n = 7 entries) with posterior probability (PP) > 0.5 are listed. For each candidate, the table reports trait category, organ, associated trait, analysis type, gene ID or lead SNP locus, chromosomal position, PP value, gene annotation, GO functional category, KEGG functional category, hotspot or locus designation, co-localisation status with candidates from the complementary analysis within ±100 kb, co-localised partner ID, distance to complementary candidate (bp), and source file. For volatile aroma candidates, assignment to the chromosome 01b aroma hotspot, chromosome 05 aroma hotspot (Hub locus), chromosome 10 secondary locus, or other aroma locus is indicated in the hotspot_or_locus column. For peel colorimetric candidates, the dominant chromosome 09 locus containing g008540.1 (UDP-rhamnose:anthocyanidin-3-glucoside rhamnosyltransferase; PP = 1.0 for Munsell color index) is designated as the fruit colorimetric locus. For anthocyanin candidates, metabolite-specific loci for individual anthocyanin species (P3G, P3R, C3G, C3M, C3dG) are listed with their respective chromosomal positions. Candidates are sorted by trait category, trait, and descending PP value.

**Supplementary Data 3. Functional annotation of *Ficus* carica (FCAr2.3) gene models, with associated RSEM expected read counts.**

The table provides functional annotation for all 45,900 annotated gene models of the FCAr2.3 reference assembly. Columns include: gene identifiers (gene_id, gene shortname, GeneSymbol); gene structure (chromosome, feature type, terminal, strand, exon number, genomic coordinates); homology-based annotation from BLAST alignment against the NCBI non-redundant protein database (top-hit accession, query and subject length, alignment score, E-value, %identity, alignment length, product description, source database, rank); and pathway/ontology assignments from KEGG (KEGG Orthology ID, Enzyme Commission number, definition, gene symbol, pathway ID and name), Clusters of Orthologous Groups (COG), and Gene Ontology (GO terms). For reference, RSEM expected read counts^53^ across 110 sequencing libraries are appended as additional columns. Expected counts account for multi-mapping reads by expectation-maximization-based probabilistic assignment among candidate loci; most genes receive integer-valued counts (uniquely assignable reads), while genes sharing sequence similarity with other loci may receive fractional counts. These values are distinct from the filtered, replicate-averaged expression matrix used as MOFA+ input (Supplementary Data 5).

**Supplementary Data 4. Expression of *MYB101-like* and control genes in publicly available hormone-treatment RNA-seq datasets from San Pedro-type fig.**

Quantification of gene expression from publicly available RNA-seq datasets of San Pedro-type fig (*Ficus carica* L.) main crop treated with exogenous gibberellin (GA₃; SRA SRP113799)^24^ or cytokinin (CPPU; SRA PRJNA397979)^47^, re-mapped to the FCAr2.3 reference assembly using a uniform pipeline. The table lists, for each gene and each experimental condition, the raw read count, normalized expression (counts per million, CPM), and log2 fold-change relative to the corresponding control library.

Genes included are the parthenocarpy candidate *MYB101-like* (Fca_r2.3chr04_g020650.1), its closest full-length GAMYB-like paralog (Fca_r2.3chr00_g005240.1), a phylogenetically distinct R2R3-MYB (Fca_r2.3chr04_g042460.1), and seven cytokinin-responsive positive control genes identified as fig orthologs of characterized *Arabidopsis* type-A response regulators (ARR family) and cytokinin oxidase/dehydrogenase (CKX) family members. Each condition is defined by tissue (female flower or receptacle), treatment (control, GA₃, or cytokinin), and timepoint (2, 4, or 10 days after treatment, DAT).

For both datasets, only a single sequencing library is available per condition (tissue × treatment × timepoint); no biological replicates are deposited in the public repositories. Consequently, log2 fold-change values are descriptive comparisons between single libraries and are not accompanied by statistical significance estimates. Sequence Read Archive (SRA) run accessions corresponding to each library are provided in the table to enable reproduction. Values should be interpreted as exploratory only, as detailed in the main text and in the legend to Supplementary Fig. 10.

**Supplementary Data 5. TMM-normalized gene expression matrix used as the RNA-expression layer input for MOFA+ multi-omics integration.**

The table reports TMM-normalized expression values (edgeR; Robinson et al., 2010), derived from RSEM expected counts^53^, for 30,081 expressed genes (read count ≥ 1) out of 45,900 annotated FCAr2.3 gene models, across 81 fig (*Ficus* carica) accessions following aggregation of technical/biological replicates. This matrix constitutes the RNA-expression layer used as input for MOFA+ multi-omics factor analysis (Figure 5, Figure 9A) and for transcriptome-wide association analysis (TWAS).

## Additional Resources

Plant Garden Database (*Ficus carica*): https://plantgarden.jp/en/list/t3494

Plant Garden Database (*Ficus erecta*): https://plantgarden.jp/en/list/t66383

